# Chromosome gigantism and auxin deconjugation underpin gall induction in a horned gall aphid

**DOI:** 10.64898/2026.06.08.730764

**Authors:** Qin Lu, Weiwei Wang, Xumei Chen, Juan Liu, Yiyuan Pan, Xin Zhang, Chenjing Ma, Hang Chen

## Abstract

Insect galls are extended phenotypes built through sustained reprogramming of host development, yet the genomic innovations and effector mechanisms that enable this process remain poorly resolved. Here we generated chromosome-level genomes for the horned gall aphid *Schlechtendalia chinensis* and its host tree *Rhus chinensis*, and integrated these resources with morph-resolved salivary proteomics, metabolomics and functional assays. The aphid genome contains an anomalously expanded chromosome 1 spanning 93.61 Mb that carries 33.08% of anchored genes and concentrates 70.47% of genome-wide segmental duplications, together with elevated transposable-element load and multiple tandem gene clusters. We identified a salivary M20-family enzyme, ACY1, that consistent with an auxin-conjugate hydrolase. Loss- and gain-of-function assays show that ACY1 hydrolyzes stored, inactive auxin–amino acid conjugates in host tissue, increasing local active auxin levels. This hormone manipulation reprograms plant cell development to form galls. Our findings demonstrate how concentrated genomic duplication can drive the evolution of an effector that controls host physiology, suggesting a potential principle in the evolution of parasitic traits.

## 1 Introduction

Insect galls represent striking examples of sustained developmental reprogramming across kingdoms[1]. They are not passive wound responses, but newly built plant organs, with growth, patterning and physiology redirected by the inducing insect[2, 3]. Among galling insects, aphids occupy a particularly distinctive niche because they can maintain enclosed galls that support multiple generations within a single host structure[4]. *Schlechtendalia chinensis* represents an especially tractable system in this regard. It induces elaborate horned galls on *Rhus chinensis*, and only two morphs in its life cycle, the fundatrix and fundatrigenia, actively participate in gall establishment and maintenance [5, 6].

How such a system evolves and how it operates mechanistically remain open at two linked levels. One concerns genome architecture. Aphid chromosomes are holocentric and can tolerate extensive structural rearrangement[7], yet it is still unclear how particular chromosomal compartments become disproportionately enlarged and functionally specialized in galling lineages. The other concerns the effector layer at the aphid–plant interface. Gall induction clearly requires persistent manipulation of host developmental programs[8], but only a small number of insect-derived factors have been connected to that process with functional evidence[9, 10].

Auxin (indole-3-acetic acid, IAA) homeostasis is central to this problem. Because IAA can generate stable local maxima through polar transport or homeostatic shifts alone, it is a primary target for gall-inducing organisms[11]. Current evidence points to at least five non-exclusive routes by which gallers manipulate host auxin level. (i) Some species synthesize IAA *de novo* and inject it directly via salivary secretions [12, 13]. (ii) Gallers can transcriptionally upregulate host IAA biosynthesis genes [14, 2, 15]. (iii) Polar-transport machinery can be co-opted to build local IAA peaks [16]. (iv) IAA-mediated suppression of jasmonate/salicylate defence can redirect the host growth–defence trade-off towards organ-building programmes that favour gall maintenance[17]. (v) Finally, and least resolved, gallers may directly alter host IAA conjugation–deconjugation metabolism, converting stored conjugates into free active auxin. In plants, GH3 acyl-amido synthetases convert free IAA into amino-acid conjugates (IAA-AA) that serve as a buffered storage pool, and M20-family amidohydrolases such as ILR1/ILL/IAR3 reverse this reaction to release active IAA[18, 19]. Although conjugate–deconjugate cycling is widely recognized as a key reversible node in IAA homeostasis[20], no galling species has yet been shown to directly exploit this reservoir through a secreted effector.

Here we combine chromosome-level genome assemblies for *S. chinensis* and *R. chinensis* with comparative genomics, morph-resolved salivary proteomics, metabolomics and functional to reveal an unusually expanded aphid chromosome 1 (chr1) that concentrates duplication, transposons and multiple tandem gene clusters. It provides an evolutionary testing ground for the rapid generation of new gene copies and functional innovations. A salivary M20 metallopeptidase, ACY1, is identified as a candidate effector linked to local auxin remodeling at the feeding site. This strategy of insects utilizing the resource pool of host plants to build an expanding refuge (galls) for themselves may be widely present in other herbivorous insects and host plants, providing new ideas for studying the interaction and co-evolution between insects and plants.

## 2 Materials and methods

Detailed protocols, software versions and extended parameter settings are provided in the Supplementary Methods. The main procedures used in the present study are summarized below.

### 2.1 Sample collection

*S. chinensis* was collected from naturally formed galls on *R. chinensis* in Yunnan, China (E102°44′49″, N25°03′50″), across two field seasons. Three aphid morphs were sampled: fundatrices, fundatrigeniae and sexuparae. Young expanding *R. chinensis* leaves from the same host trees were used for host genome sequencing and for downstream functional assays. Salivary glands (SGs) were dissected from defined aphid morphs under ice-cold conditions and stored at −80 °C until analysis. Saliva was collected using an artificial membrane feeding system. Briefly, aphids were placed on Petri dishes sealed with stretched Parafilm membrane over the collection fluid and allowed to feed for 12 h under controlled environmental conditions. For proteomic analysis, the collection fluid consisted of a chemically defined artificial diet containing amino acids and 15% (w/v) sucrose at pH 7.0. For metabolomic analysis, the collection fluid consisted of 15% (w/v) sucrose solution prepared with LC–MS-grade water. After feeding, the Parafilm membranes were removed, and the diet solutions containing secreted watery saliva were recovered, passed through 0.22-μm filters to remove cellular debris, concentrated by ultrafiltration (3-kDa molecular weight cut-off), and stored at −80 °C. Five sample groups were designated as follows: fundatrix salivary gland (Fx-SG), fundatrigenia salivary gland (Fa-SG), sexuparae salivary gland (Se-SG), fundatrix saliva (Fx-S) and fundatrigenia saliva (Fa-S).

### 2.2 Genome sequencing, assembly and annotation

High-molecular-weight DNA from *S. chinensis* and *R. chinensis* was sequenced using PacBio HiFi long reads and Illumina short reads, and chromosome scaffolding was achieved with Hi-C. Assemblies were generated with hifiasm [21, 22], polished with long- and short-read evidence, and scaffolded with Juicer and 3D-DNA [23, 24] followed by manual curation. Assembly completeness was evaluated with Benchmarking Universal Single-Copy Orthologs (BUSCO) [25], and chromosome numbers were independently validated by DAPI-FISH and Giemsa staining. Protein-coding genes were annotated using a combined *ab initio*, transcript-supported and homology-based workflow, and functional annotations were assigned against NR, Swiss-Prot, Kyoto Encyclopedia of Genes and Genomes (KEGG), KOG/eggNOG and InterPro databases. Repetitive sequences were identified through *de novo* and homology-based approaches.

### 2.3 Comparative genomics and chromosome-level analyses

Orthogroups were inferred with OrthoFinder [26], and gene-family expansion and contraction were estimated with CAFE [27]. Positive selection was tested with branch-site models in codeml from PAML [28] using filtered codon alignments of single-copy orthologues. Intra- and interspecies synteny were inferred with MCScanX [29]. Segmental duplications (SDs) were called from all-versus-all protein similarity searches with stringent identity and alignment thresholds and were then summarized by chromosome. Chr1-focused analyses included category-specific density comparisons, KEGG enrichment of chr1-resident genes, manual re-annotation of genes assigned to lysine degradation, and mapping of tandem clusters such as opsins, crystallin/small heat shock protein 20 (sHSP20) genes and PIF1 helicases.

### 2.4 Salivary proteomics

The five aphid sample groups (Fx-SG, Fa-SG, Se-SG, Fx-S and Fa-S) were profiled. Proteins were extracted in SDS–DTT–Tris-HCl buffer (SDT buffer), digested by filter-aided sample preparation and analysed by liquid chromatography–tandem mass spectrometry (LC–MS/MS) on an Orbitrap Exploris 480 platform. Raw files were searched with MaxQuant [30] against the predicted *S. chinensis* proteome, using a peptide and protein false discovery rate (FDR) below 1%. Label-free quantification (LFQ) was performed with MaxLFQ. Differential abundance was assessed from log_2_-transformed LFQ intensities using empirical Bayes moderated t-tests in limma, with Benjamini–Hochberg correction; proteins with |log_2_FC| > 1 and adjusted *P* < 0.05 were considered significantly differential.

### 2.5 Untargeted metabolomics

Metabolites from the same saliva and SG sample groups were extracted in 80% methanol and analysed on a Vanquish UHPLC system coupled to a Q Exactive HFX Orbitrap mass spectrometer in both ionization modes. Peaks were aligned and annotated in Compound Discoverer using mzCloud, HMDB, KEGG and ClassyFire-supported workflows. Differential metabolites were defined using |log2FC| > 1 and *P* < 0.05 with correction for multiple testing. Pathway enrichment was assessed against KEGG annotations.

### 2.6 RNA interference and targeted auxin profiling

A dsRNA fragment targeting *ACY1* was synthesized *in vitro* and delivered to newly born fundatrix nymphs through an artificial diet for 24 h. Aphids were then transferred to detached young *R. chinensis* leaves for gall-induction assays. Galling rate and mortality were scored at 25 d. Knockdown efficiency was quantified by RT–qPCR, with primer information provided in Table S1. For auxin profiling, tissue surrounding the aphid settlement site was excised and analysed by targeted LC–MS/MS with stable-isotope internal standards to quantify free IAA and seven conjugated IAA species.

### 2.7 Cell-free expression and delivery

The full-length ACY1 coding sequence was expressed in a wheat-germ cell-free system using the pEU-E01-MCS-N-His expression vector together with the WEPRO7240H bilayer reaction format (CellFree Sciences). The resulting crude lysate contained recombinant ACY1 alongside the endogenous wheat-germ proteins required for translation; total protein concentration was therefore measured by BCA assay, and apparent ACY1 concentrations are reported throughout the cell-free experiments as the BCA-based total-protein mass divided by the predicted molecular mass of ACY1 (∼45.8 kDa), rather than as concentrations of affinity-purified enzyme (see Supplementary Methods 8.1). Heat-inactivated ACY1 (HI-ACY1) and empty-vector lysate (EV), each matched in total-protein concentration to the corresponding active treatment, served as controls. For in planta validation, young R. chinensis leaves were first punctured with a fine needle to create small wounds mimicking aphid stylet penetration, then vacuum-infiltrated with lysate diluted to apparent ACY1 concentrations of 1, 2, 4 and 8 μM. Infiltrated leaves were placed in humidity chambers and incubated for 12 h before sampling for targeted auxin profiling.

### 2.8 Histology

Settlement-zone tissues from control and *ACY1*-RNAi treatments were fixed in FAA, paraffin-embedded, sectioned and stained with Safranin O–Fast Green. Cell diameters were measured in epidermal, palisade mesophyll and spongy mesophyll tissues using ImageJ.

### 2.9 Statistics and reproducibility

All tests were two-sided. Multi-group comparisons were analysed by one-way ANOVA followed by Tukey’s HSD or Holm-adjusted *post hoc* testing as appropriate. Pairwise comparisons in the cell-free assays used Welch’s *t*-tests with Holm correction. Correlations were evaluated using Pearson’s *r* with Benjamini–Hochberg FDR correction. Histological measurements were compared with Mann–Whitney *U* tests. Effect sizes for the cell-free assays were expressed as Hedges’ *g*. Unless otherwise stated, data are presented as mean ± s.d., and all functional experiments included at least six biological replicates.

## 3 Results

### 3.1 A gigantic, duplication-driven chromosome 1 in S. chinensis

We generated chromosome-level assemblies for both partners in the *S. chinensis*–*R. chinensis* interaction. The aphid assembly resolved 16 chromosomes and 19,109 predicted protein-coding genes, with 18,096 genes anchored to chromosomes, and cytogenetic validation confirmed a diploid chromosome number of 2*n* = 32 (Fig. 1a, b; Fig. S1, Tables S2–S12). The most striking feature of the aphid genome is the scale of chr1. At 93.61 Mb, chr1 is far larger than any other aphid chromosome in the assembly and contains 5,986 genes, corresponding to 33.08% of all chromosome-anchored genes (Fig. 1c; Table S3). Hi-C contact patterns remain continuous across its full length, indicating that this chromosome is a coherent chromosomal unit rather than a misjoined scaffold (Fig. S2).

**Fig. 1.**
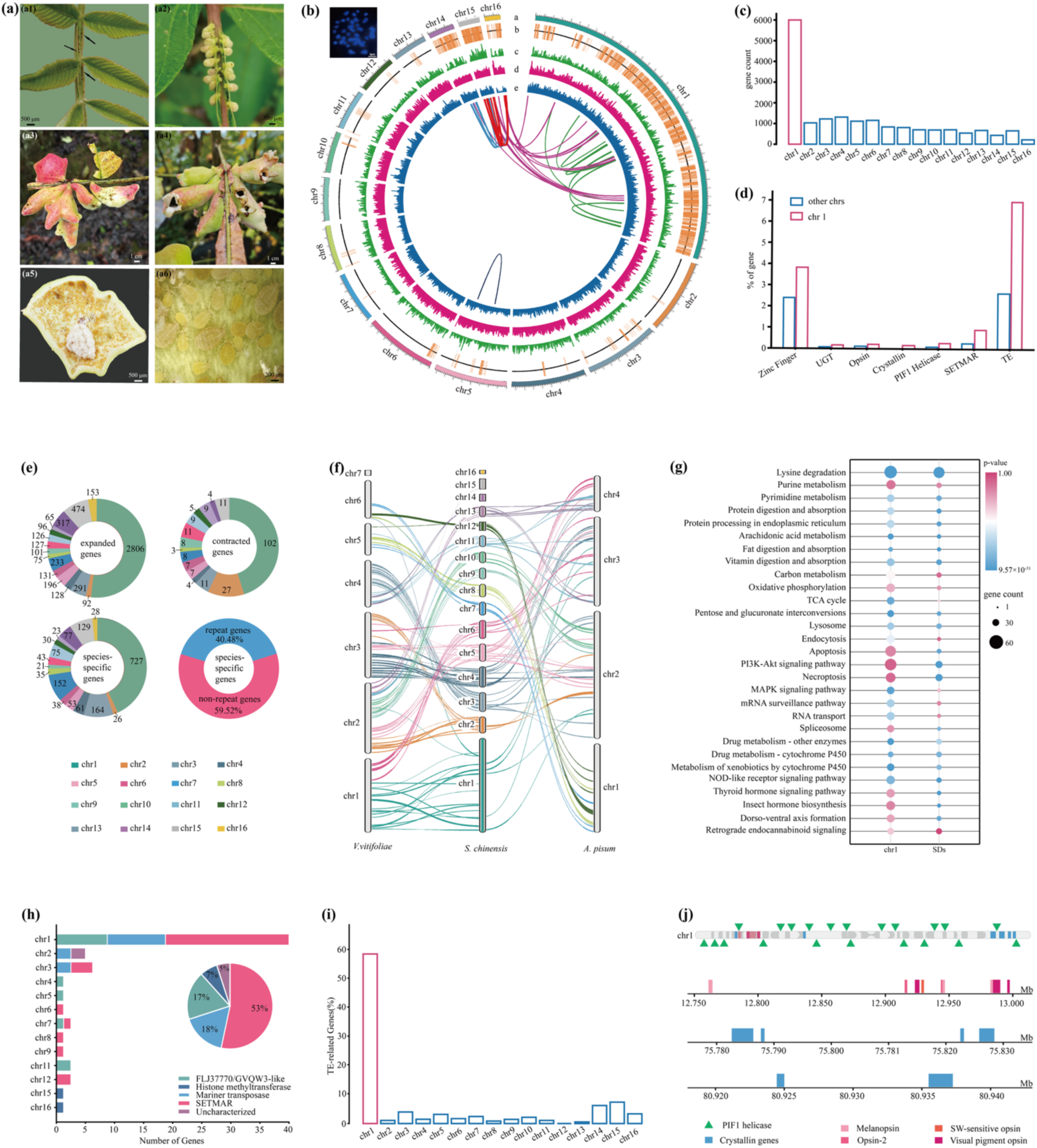
Chromosome-level genome assembly and structural features of *S. chinensis*. (a) Photographs of horned galls on *R. chinensis* leaves at different developmental stages. (a1) Fundatrix feeding on a young Rhus chinensis leaf wing. (a2) Early stage of horned gall development. (a3) Late stage of horned gall development. (a4) Dehisced (burst) horned gall. (a5) Interior of a horned gall. (a6) Fundatrigeniae inside the gall. (b) DAPI-FISH karyotype (2*n* = 32) and Circos plot of the *S. chinensis* genome. The Circos plot showing chromosome ideograms, gene density, GC content, repeat density, and intra-genomic synteny (tracks a–e from outer to inner). (c) Gene count per chromosome; chr1 harbours 5,986 genes (33.08% of of anchored genes). (d) Proportion of selected gene categories (zinc finger, UGT, opsin, crystallin, *PIF1* helicase, *SETMAR* and TE-related) on chr1 versus other chromosomes. (e) Chromosomal distribution of expanded gene families, contracted gene families, species-specific genes and the repeat portion of species-specific genes. (f) Three-way synteny comparison among *V. vitifoliae*, *S. chinensis* and *A. pisum*, showing that chr1 shares syntenic blocks with multiple chromosomes in each outgroup. (g) KEGG pathway enrichment of chr1-resident genes (left) and of genes overlapping segmental duplications (SDs) (right). (h) Composition of genes annotated to the lysine-degradation pathway by chromosome, revealing that the majority are *SETMAR*/Mariner-derived sequences. (i) Proportion of TE-related genes per chromosome, expressed as a percentage of the genome-wide total; chr1 alone harbours 58.87% of all TE-related genes. (j) Representative tandem gene clusters on chr1, including PIF1 helicases (green triangles), crystallin/sHSP20 genes (blue boxes) and opsin genes of four spectral classes (melanopsin, opsin-2, SW-sensitive opsin and visual pigment opsin), each shown as a distinctly coloured bar.

To place chr1 in an evolutionary context, we compared the aphid assembly with *Viteus vitifoliae* (galling aphid) and *Acyrthosiphon pisum* (non-galling aphid). Chr1 shares syntenic blocks with multiple chromosomes in both outgroups (Fig. 1f), indicating that it contains material corresponding to several ancestral chromosomal elements. SDs are overwhelmingly concentrated on chr1, which harbours 70.47% of all genome-wide SD events (Table S13). Repeat-associated features are similarly concentrated. Chr1 harbours 1,682 TE-related genes — 58.87% of all TE-related genes in the genome (Fig. 1i; Table S14). Together, these patterns indicate that chr1 gigantism is maintained primarily by ongoing duplication and repeat accumulation rather than by a single historical rearrangement alone.

The enlarged chromosome is not simply gene-rich; it is functionally structured. Several categories of adaptive gene families show pronounced enrichment on chr1 (Fig. 1d). In addition, lineage- and species-specific genes are also disproportionately mapped to chr1 (Figs. S4–S6). All nine genome-wide crystallin/sHSP20 genes are confined to chr1 and arranged in tandem clusters (Fig. 1j), an organisation consistent with repeated expansion of stress-responsive chaperone loci. A second prominent feature is a dense opsin cluster spanning roughly 240 kb and containing nine tandemly arrayed opsin genes representing four spectral classes (Fig. 1j).

Chr1 also carries marked amplification of *PIF1* helicases and *SETMAR*-related loci (Fig. 1d). This is important because the strongest apparent KEGG signal on chr1, lysine degradation, does not mainly reflect canonical catabolic enzymes (Fig. 1g). Manual re-annotation showed that most of the genes contributing to this signal are in fact Mariner-derived sequences or SETMAR fusions rather than bona fide lysine-degradation genes (Fig. 1h). The chromosome therefore appears to combine repeat expansion with a parallel increase in loci implicated in replication stability and epigenetic control.

Gene-family analysis further underscored the distinctive evolutionary behaviour of chr1. Across the aphid genome, 2,806 gene families were expanded and 727 genes were species-specific, with a disproportionate fraction mapping to chr1 (Fig. 1e; Figs. S7–S9, Table S15). By contrast, positive selection showed a different distribution. Of the 308 positively selected genes identified genome-wide, only 13.31% mapped to chr1, substantially below the chromosome’s 33.08% share of anchored genes (Fig. 1e; Fig. S10, Table S16). This decoupling suggests that the dominant mode of chr1 innovation is not widespread amino-acid diversification. Instead, the chromosome appears to have accumulated adaptive potential primarily through copy-number expansion, tandem amplification and repeat-associated restructuring.

### 3.2 The host genome defines a broad response landscape for gall development

We also assembled a chromosome-level genome for *R. chinensis*, resolving 15 chromosomes and 23,451 predicted protein-coding genes, with a diploid chromosome number of 2*n* = 30 confirmed cytologically (Fig. 2a, b; Fig. S11, Tables S17–S27). In contrast to the aphid genome, host genes are distributed relatively evenly across chromosomes. Even so, the host genome bears clear signatures of functional specialization relevant to the aphid–plant interface. Expanded gene families are enriched for pentose and glucuronate interconversions, xenobiotic metabolism by cytochrome P450, drug metabolism and transporter functions (Fig. 2c–g; Figs. S12–S19, Table S28). Cell-wall remodelling genes are especially prominent, including a large set of pectin methylesterases. Detoxification-related superfamilies, including glutathione S-transferases (GSTs), UDP-glucuronosyltransferases (UGTs), ABC transporters and cytochrome P450 enzymes (CYP450s), are also broadly expanded. A notable example of tandem amplification is a cluster of six squalene epoxidase (*SQLE*) genes arranged within an approximately 80-kb segment on *R. chinensis* chr1 (Fig. 2h). Positive selection in plant hormone signal transduction and plant–pathogen interaction pathways further indicates that the host genome has been shaped by sustained pressure at the interaction interface.

**Fig. 2.**
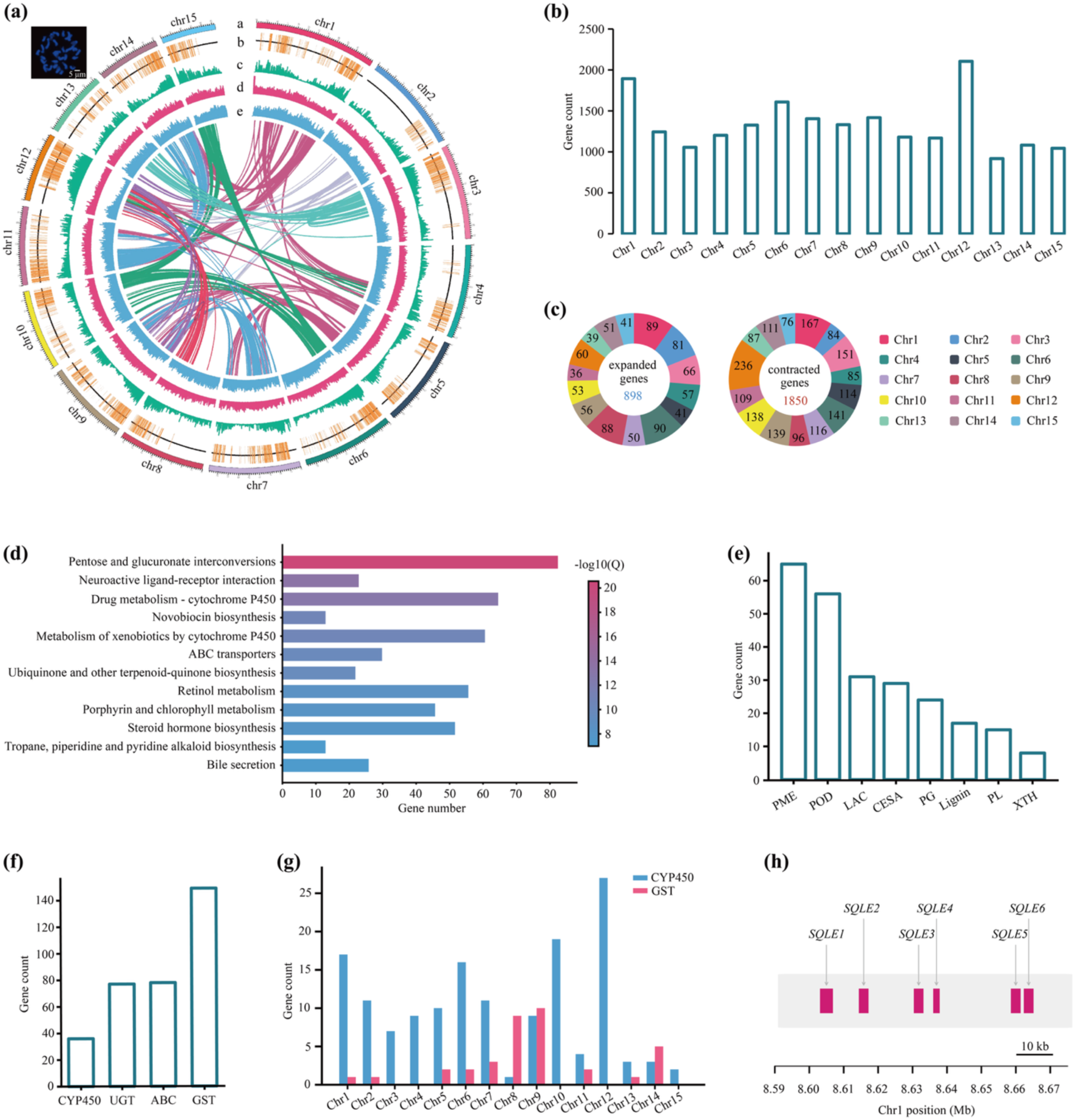
Chromosome-level genome assembly and genomic features of *R. chinensis*. (a) Circos plot of the host genome showing chromosome ideograms, gene density, GC content, repeat density, and intra-genomic synteny (tracks a–e from outer to inner). (b) Gene count per chromosome, showing a relatively even distribution. (c) Chromosomal distribution of expanded and contracted gene families. (d) KEGG pathway enrichment of expanded gene families, with pentose and glucuronate interconversions and cytochrome P450-mediated metabolism as the most enriched categories. (e) Gene counts of cell-wall remodelling families including pectin methylesterases (PME), peroxidases (POD), laccases (LAC), cellulose synthases (CESA), polygalacturonases (PG), lignin-related enzymes (Lignin), pectate lyases (PL) and xyloglucan endotransglucosylases (XTH). (f) Gene counts of major detoxification superfamilies (CYP450, UGT, ABC, GST). (g) Chromosomal distribution of CYP450 and GST genes. (h) A tandem array of six *SQLE* genes within an ∼80-kb segment on chr1, consistent with local duplication of a triterpenoid biosynthesis gene cluster.

### 3.3 Morph-resolved salivary proteomics prioritizes ACY1 as a gall-associated M20 candidate

To identify candidate effectors, we profiled salivary glands (Fig. 3a) and saliva from three aphid morphs: the fundatrix, which induces gall formation, has a stylet length of 167.52 ± 14.86 μm (Fig. 3b, c); the fundatrigenia, which maintains gall growth, has a stylet length of 127.34 ± 17.28 μm (Fig. 3d); and the sexupara, which does not induce galls, has a vestigial stylet (Fig. 3e).

**Fig. 3.**
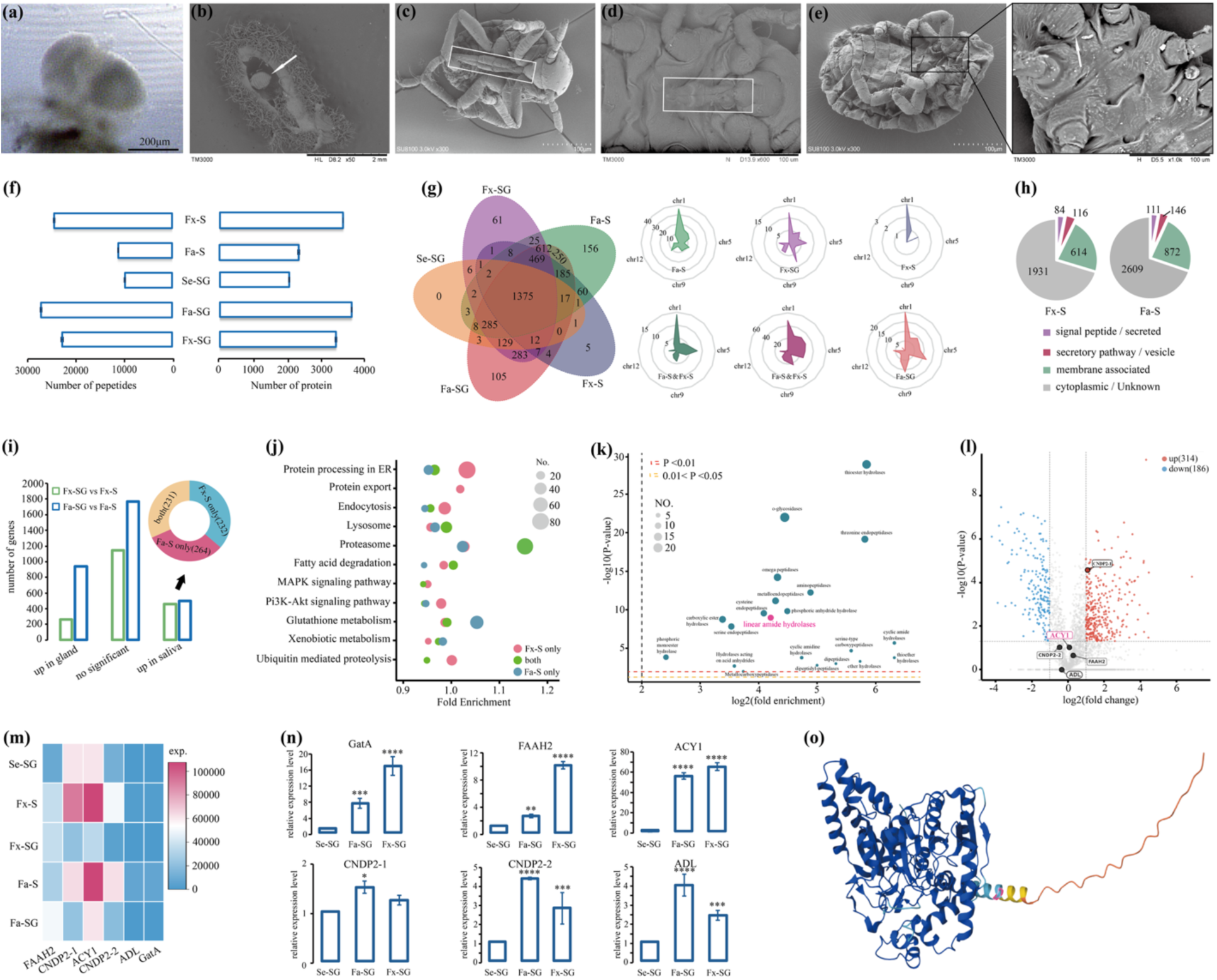
Morph-resolved salivary proteomics of *S. chinensis*. (a) A salivary gland of fundatrigenia. (b) A fundatrix inside the gall. (c–e) Scanning electron micrographs of stylet from fundatrix, fundatrigenia and sexuparae morphs, stylet absent in sexuparae morphs. (f) Number of peptides (left) and proteins (right) identified in each sample group. (g) Five-set Venn diagram showing protein overlaps among Fx-SG, Fa-SG, Se-SG, Fx-S and Fa-S; 1,375 proteins are shared across all groups and no morph-specific proteins were identified in the non-galling sexuparae. Radar plots show chromosomal distribution of proteins in each group. (h) Predicted secretion classification of saliva proteins. (i) Numbers of differentially abundant proteins between gland and saliva comparisons (upregulated in gland versus saliva). The donut chart shows that among the upregulated proteins, 231 were shared between fundatrix and fundatrigenia, 232 were specific to fundatrix (Fx-S only), and 264 were specific to fundatrigenia (Fa-S only). (j) KEGG pathway enrichment of Fa-S versus Fx-S differentially abundant proteins. (k) Bubble plot of enzyme-class enrichment in Fa-S and Fx-S -enriched proteins, highlighting linear amide hydrolases. (l) Volcano plot of Fa-S versus Fx-S comparison, with M20-family candidates ACY1, CNDP2-1, CNDP2-2, ADL and FAAH2 labelled. (m) Expression heatmap of six M20/amidohydrolase candidates across all five sample groups. (n) RT–qPCR validation showing that *ACY1* transcript abundance is highest in Fa-SG and Fx-SG relative to Se-SG. (o) AlphaFold2-predicted monomeric structure of ACY1. Blue, protein chain; orange, signal peptide.

Quantitative proteomics identified 4,142 proteins across the five sample groups (Fig. 3f; Figs. S20–S24, Table S29). Of these, 1,375 were shared across all groups, whereas each saliva and gland type also retained a distinct morph-specific component besides Se-SG (Fig. 3g). In both gall-inducing morphs, saliva proteins were predominantly classified as cytoplasmic/unknown — i.e., lacking a classical secretion signal — accounting for 70.35% of the saliva proteome in Fx-S and 69.80% in Fa-S, followed by membrane-associated proteins at 22.37% and 23.33%, respectively, whereas only 3.06% and 2.97% were predicted to be classically secreted through a signal peptide, and a further 4.23% and 3.91% fell into the secretory-pathway/vesicle category (Fig. 3h). The comparison most relevant to gall maintenance, Fa-S versus Fx-S, revealed 314 upregulated and 186 downregulated proteins (Fig. 3l). Detailed profiling of salivary gland and saliva proteins across the three morphs revealed both shared and morph-specific features (Fig. 3g, i, Figs. S25–S49). In all five sample groups, the secretome was dominated by proteins involved in protein processing and export, lysosomal function, endocytosis and proteasome-mediated turnover, consistent with the high secretory activity of aphid salivary glands. Metabolic enzymes related to fatty acid degradation, glutathione metabolism and xenobiotic processing were also broadly represented, suggesting that the salivary apparatus serves not only in effector delivery but also in detoxification at the feeding interface. Among the morph-specific differences, gall-associated morphs (fundatrix and fundatrigenia) showed a pronounced enrichment of hydrolytic enzymes, particularly peptidases, amidohydrolases and carboxylesterases, relative to the non-galling sexuparae. A total of 961 hydrolases were detected in the *S. chinensis* proteome, broadly involved in apoptosis, autophagy, proteasomal degradation, lysosomal digestion, and multiple signal transduction pathways, with saliva-enriched linear amide hydrolases and M20 metallopeptidase family proteins identified as key candidate effectors for manipulating host plant auxin metabolism. Chromosome 1, a giant chromosome harboring 33.08% of all predicted genes, concentrated 20.08% of the detected hydrolases (193 genes), whose KEGG pathways were predominantly enriched in apoptosis, autophagy, PI3K-Akt and mTOR signaling, phospholipase D signaling, and oxidative phosphorylation(Table S30, Fig. S49) — consistent with chr1’s role as an evolutionary hotspot of transposon-driven gene family expansion and reflecting its functional specialization in programmed cell death regulation, protein turnover, and energy metabolism.

KEGG pathway enrichment of these Fa-S versus Fx-S differentially abundant proteins highlighted protein processing, lysosome and proteasome (Fig. 3j). Functional enrichment converged on peptidases and amidohydrolases (Fig. 3k, l). Among M20-domain candidates, ACY1 was particularly notable because it was abundant in gall-associated saliva and showed elevated transcript levels in Fa-S and Fx-S (Fig. 3m, n; Table S31). Structural modelling of ACY1 revealed a monomeric metallopeptidase fold characteristic of the M20 family, with a conserved binuclear zinc-binding active site positioned to accommodate aminoacyl substrates (Fig. 3o).

### 3.4 Salivary metabolomics reveals a substrate-rich chemical environment in gall-inducing morphs

Untargeted metabolomics of the same five sample groups identified 1,475 metabolites in total (Fig. 4a, b; Figs. S50–S53; Table S32). At the superclass level, of the 1,313 metabolites assigned to a superclass, organic acids and derivatives constituted the largest category (327; 24.90%), followed by lipids and lipid-like molecules (305; 23.23%) and organoheterocyclic compounds (196; 14.93%) (Fig. 4a; Figs. S52, S53). Class-level annotation identified carboxylic acids and derivatives (19.05%), fatty acyls (9.36%) and organooxygen compounds (8.95%) as the most represented classes (Fig. S52). Volcano plots for individual comparisons further illustrated broad differential distributions among salivary glands of different morphs (Figs. S54–S56), and KEGG pathway enrichment consistently highlighted amino acid metabolism and biosynthesis pathways across multiple comparisons (Figs. S57–S63).

**Fig. 4.**
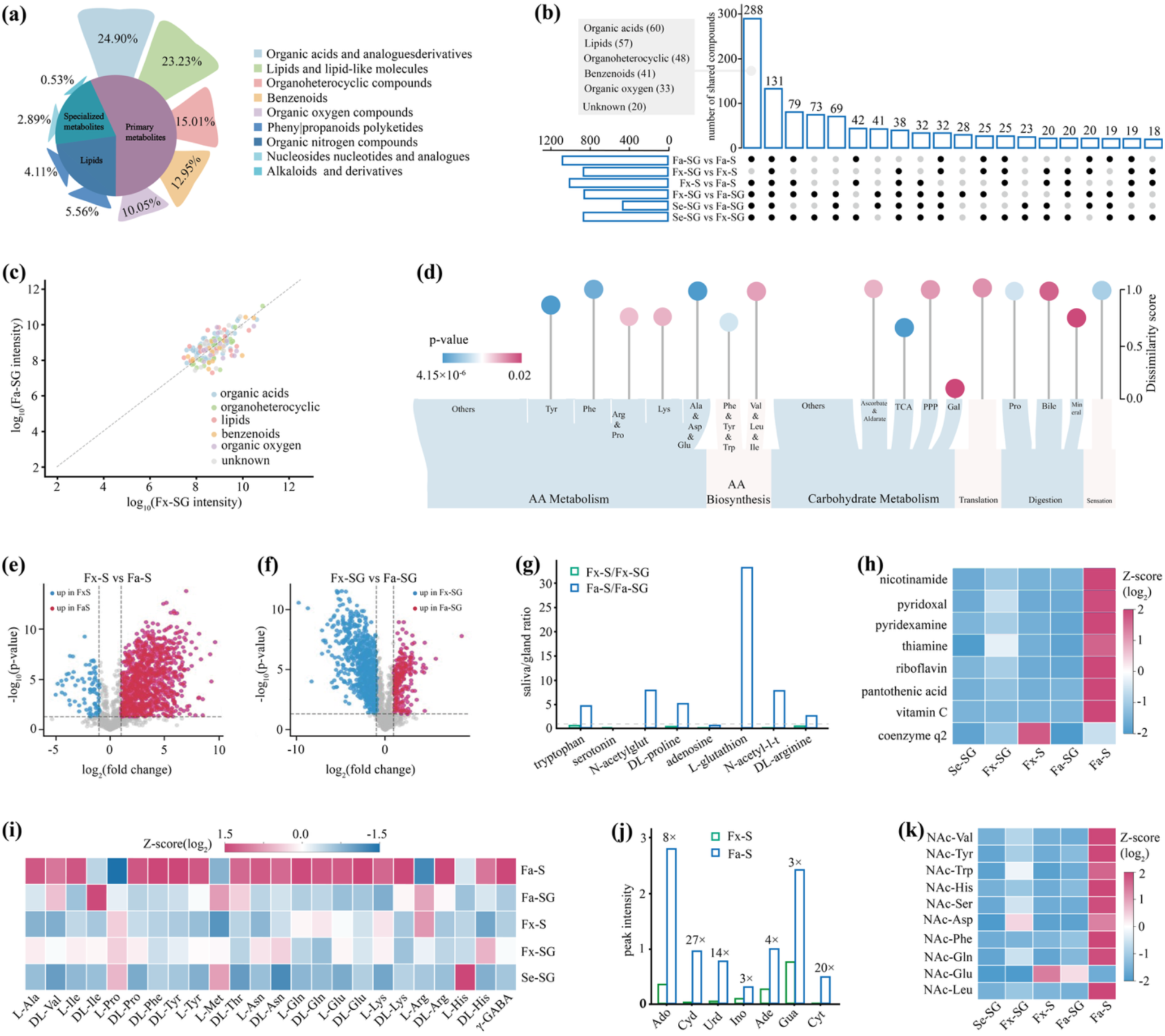
Untargeted metabolomics of aphid saliva and salivary glands. (a) Chemical superclass composition of the 1,313 metabolites assigned to a superclass (out of 1,475 detected in total); organic acids and lipids are the top two categories. (b) UpSet plot showing the intersection of differential metabolites across six pairwise comparisons; 288 metabolites differ in five comparisons and 131 in all six. (c) Scatter plot comparing log-transformed metabolite intensities between Fx-SG and Fa-SG. (d) KEGG-based metabolic pathway enrichment of differential metabolites, including amino acid metabolism, biosynthesis, carbohydrate metabolism, translation, digestion and sensation pathways. (e–f) Volcano plots of differential metabolites between Fx-S versus Fa-S and Fx-SG versus Fa-SG. (g) Saliva-to-gland ratios of selected metabolites in Fx and Fa morphs. (h) Z-score heatmap of B-vitamin abundance across sample groups. (i) Z-score heatmap of free amino acid abundance across all sample groups. (j) Abundance of purine nucleosides (adenosine, cytidine, uridine, inosine, guanosine) in Fx versus Fa saliva. (k) Z-score heatmap of N-acetyl amino acids across all sample groups, showing enrichment of N-acetyl-Tyr, N-acetyl-Ser and N-acetyl-Gln in Fa-S.

The most functionally informative contrast came from gall-associated saliva. KEGG pathway enrichment across all sample groups showed that amino acid metabolism, encompassing multiple sub-pathways including tyrosine, phenylalanine, lysine, and alanine/aspartate/glutamate metabolism, and amino acid biosynthesis were the most prominently enriched functional categories (Fig. 4d; Figs. S57–S63). The comparison between Fx-S and Fa-S likewise showed significant enrichment in amino acid metabolic pathways among differential metabolites (Fig. 4e; Fig. S61). A parallel comparison of the salivary glands (Fx-SG versus Fa-SG) examined overall metabolite intensities (Fig. 4c) and the differentially abundant features distinguishing the two glands (Fig. 4f).

More specifically, Fa-S was enriched for free amino acids and N-acetyl amino acids. Z-score profiling of free amino acids showed that Fa-S had elevated abundances of most amino acids relative to other sample groups, particularly L-Ala, DL-Val, DL-Ile, L-Pro, DL-Phe and DL-Tyr (Fig. 4i). Several N-acetyl amino acids were also markedly enriched in Fa-S, including NAc-Tyr, NAc-Ser, NAc-Asp, NAc-Phe, NAc-Gln and NAc-Glu (Fig. 4k). Saliva-to-gland metabolite ratios indicated that tryptophan, serotonin, N-acetyl-glutamate, DL-proline, adenosine and related compounds were preferentially secreted into Fa-S rather than retained in the gland (Fig. 4g). B-vitamin cofactors, including nicotinamide, pyridoxal, pyridoxamine, thiamine, riboflavin and pantothenic acid, were also more abundant in gall-inducing morph saliva (Fig. 4h).

Nucleoside metabolites were strikingly upregulated in Fa-S, with adenosine approximately 8-fold, cytidine 27-fold, uridine 14-fold and inosine 3-fold higher than in Fx-S (Fig. 4j).

These metabolomic data converge with the proteomic findings on the same functional axis. Gall-associated saliva is enriched not only in M20-family peptidases and amidohydrolase candidates, but also in a chemical milieu rich in free amino acids, N-acetyl amino acids and other aminoacyl compounds. This substrate landscape is consistent with the enzymatic specificity of M20 metallopeptidases, which act on aminoacyl substrates.

### 3.5 ACY1 knockdown reduces galling success and shifts local auxin homeostasis towards conjugated forms

We next tested *ACY1* directly by RNAi. Feeding fundatrix nymphs dsRNA targeting *ACY1* for 24 h reduced transcript abundance to 0.238 ± 0.093 relative to controls (Fig. 5a, b; Table S33). Galling rate dropped from ∼90% in both control groups to 52.33 ± 6.02%, while mortality rose correspondingly to 47.67 ± 6.73% (Fig. 5a, c, d; Table S33). Targeted LC–MS/MS profiling at the aphid settlement zone showed that this phenotype was accompanied by a substantial shift in local auxin balance (Fig. 5e; Table S34). Free IAA declined significantly, whereas all seven measured conjugated forms increased, including IAA-Leu, IAA-Phe, IAA-Tyr, IAA-Ala, IAA-Asp, IAA-Glu and IAA-Glc. Correlation analysis reinforced this pattern: *ACY1* expression was strongly positively associated with galling rate and free IAA, and negatively associated with mortality and several IAA conjugates (Fig. 5f–i; Fig. S64 and S65, Table S35). Histological analysis further showed that cell expansion in the settlement zone was reduced after *ACY1* knockdown, affecting epidermal, palisade and spongy mesophyll tissues (Fig. 5j–l; Table S36).

**Fig. 5.**
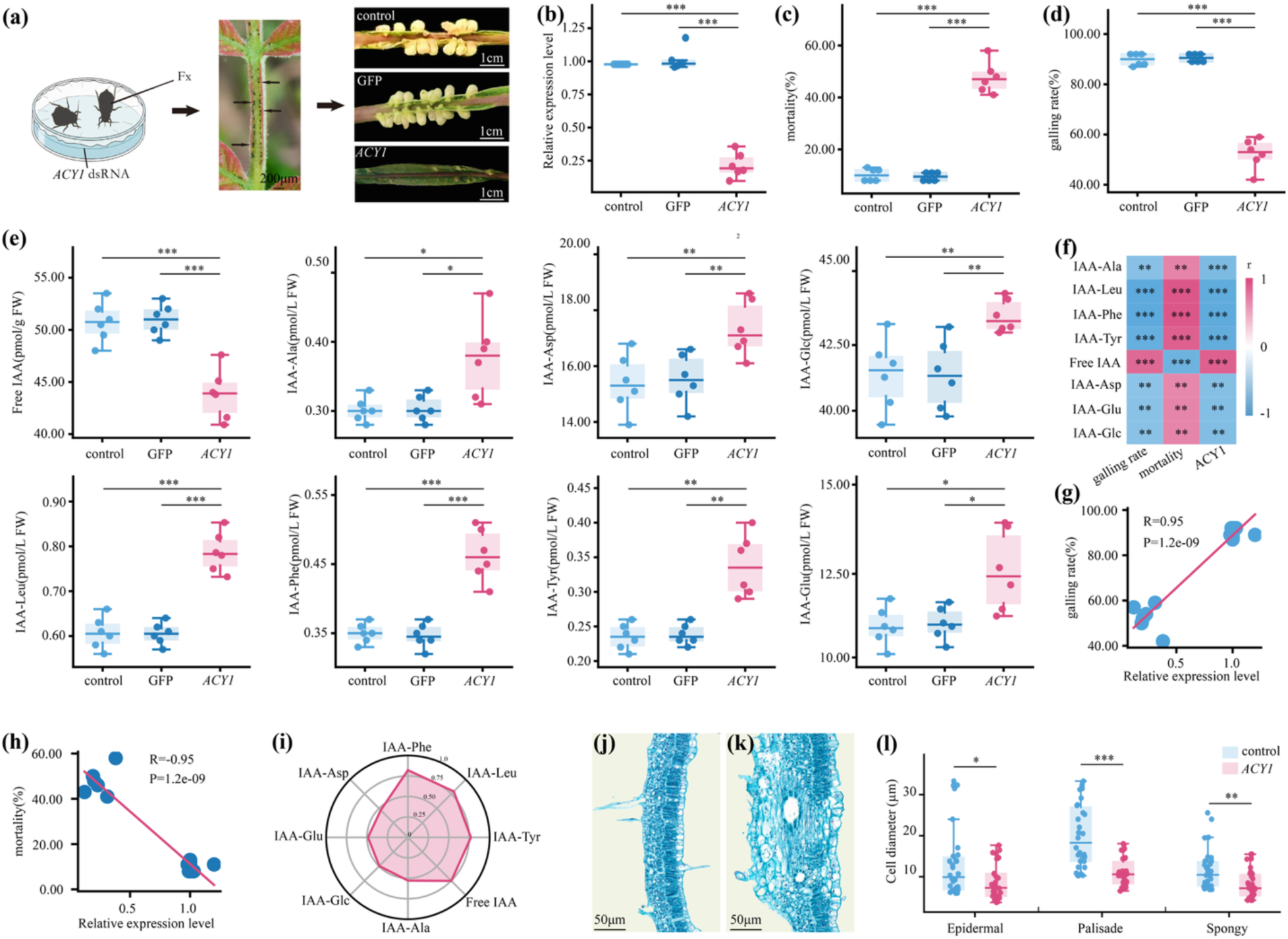
*ACY1* RNAi phenotype and auxin profiling at the galling site. (a) Experimental design: fundatrix nymphs were fed dsRNA targeting *ACY1* for 24 h, then transferred to detached *R. chinensis* leaves. (b) RT–qPCR confirmation of *ACY1* knockdown efficiency (relative expression reduced to 0.238 ± 0.093). (c) Mortality rose from 10.17 ± 2.29% and 9.50 ± 1.64% in two control groups to 47.67 ± 6.73% after *ACY1* knockdown. (d) Galling rate decreased from 89.83 ± 2.40% and 90.50 ± 1.64% in controls to 52.33 ± 6.02% after knockdown. (e) Targeted LC–MS/MS quantification of free IAA and seven IAA conjugates (IAA-Ala, IAA-Asp, IAA-Leu, IAA-Phe, IAA-Tyr, IAA-Glu, IAA-Glc) at the settlement zone; free IAA declined and all conjugates increased after *ACY1* knockdown. (f) Pearson correlation matrix among IAA metabolites, mortality, galling rate and *ACY1* expression. (g) Scatter plot showing strong positive correlation between *ACY1* expression and galling rate (R = 0.95). (h) Scatter plot showing strong negative correlation between *ACY1* expression and mortality (R = −0.95). (i) Radar chart of normalised IAA conjugate levels after *ACY1*-RNAi treatments. (j–k) Representative histological cross-sections of settlement-zone tissue stained with Safranin O–Fast Green in *ACY1*-RNAi treatments (j) and control (k). (l) Cell diameter measurements in epidermal, palisade mesophyll and spongy mesophyll tissues, showing significant reduction after *ACY1* knockdown.

### 3.6 Cell-free ACY1 treatment shifts host auxin pools in a dose-dependent manner

To test whether ACY1 activity is sufficient to alter host auxin balance *in planta*, we expressed ACY1 in cell-free systems and delivered crude enzyme preparations to young *R. chinensis* leaves by vacuum infiltration following fine-needle puncture to mimic aphid stylet penetration. Active ACY1 induced clear dose-dependent changes in auxin metabolite profiles. Free IAA levels rose progressively with increasing ACY1 concentration and peaked at 4 μM, reaching 59.40 ± 2.30 pmol g⁻¹ FW compared with 48.14 ± 2.15 pmol g⁻¹ FW in the HI-ACY1 control (Fig. 6a, Table S37). Correspondingly, the ratio of free IAA to total IAA–amino acid conjugates increased in a dose-dependent manner, with 2, 4 and 8 μM treatments all significantly elevated relative to controls (Fig. 6c, Table S37). The proportion of free IAA as a percentage of total measured IAA showed the same trend, rising to 53.06 ± 2.02% at 4 μM (Fig. 6d, Table S37).

**Fig. 6.**
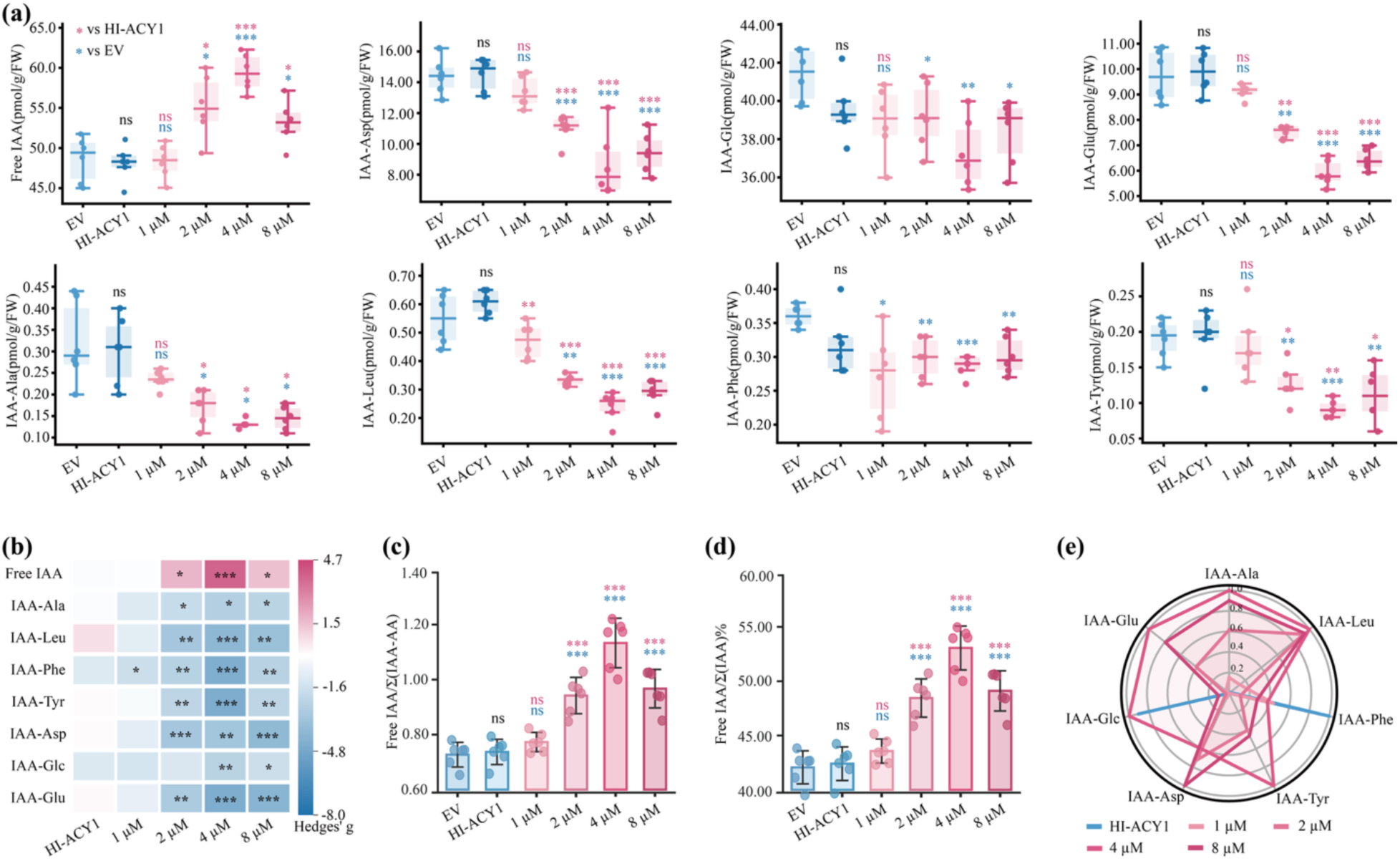
Cell-free ACY1 enzyme treatment shifts host auxin pools in a dose-dependent manner. (a) Individual auxin metabolite concentrations (free IAA, IAA-Asp, IAA-Glc, IAA-Glu, IAA-Ala, IAA-Leu, IAA-Phe, IAA-Tyr) across dose groups (EV, HI-ACY1, 1, 2, 4 and 8 μM); free IAA rises and conjugates decline with increasing ACY1 concentration. (b) Heatmap of effect sizes (Hedges’ *g*) for each auxin trait relative to HI-ACY1 control; the 4 μM concentration produces the strongest effects. (c) Ratio of free IAA to total conjugated IAA (IAA-AA), showing dose-dependent increase. (d) Percentage of free IAA relative to total measured IAA, peaking at 53.06% at 4 μM. (e) Radar chart showing the effects of different ACY1 concentrations on the levels of individual IAA conjugates.

At the level of individual conjugated species, several IAA–amino acid conjugates declined sharply under active ACY1 treatment. IAA-Leu, IAA-Phe, IAA-Tyr, IAA-Asp and IAA-Glu all showed significant reductions at 4 μM relative to both EV and HI-ACY1 controls, with IAA-Leu and IAA-Phe exhibiting the largest effect sizes. IAA-Ala declined at 2 μM and above, though with greater variability at higher concentrations. IAA-Glc also decreased at 4 μM and 8 μM. By contrast, the 1 μM treatment produced only modest and largely non-significant shifts across most conjugate species, indicating a threshold below which the enzyme preparation does not measurably alter the auxin pool (Fig. 6a, Table S37).

Effect-size analysis using Hedges’ g quantified the magnitude and direction of each treatment relative to the HI-ACY1 control (Fig. 6b, Table S37). The 4 μM concentration produced the largest positive effect on free IAA and the strongest negative effects on IAA-Leu, IAA-Phe, IAA-Tyr, IAA-Asp and IAA-Glu, with several effect sizes exceeding |g| > 4. The 8 μM treatment retained large effects on most conjugate species but did not further increase free IAA beyond 4 μM, suggesting a plateau at the highest dose. Dose–response curves for individual auxin species confirmed these patterns, with most conjugates showing monotonic declines from 2 μM onward (Fig. 6a).

Correlation analysis for the 4 μM treatment revealed strong positive associations between ACY1 concentration and free IAA levels, and strong negative associations between ACY1 concentration and the abundance of individual IAA conjugates (Fig. S66). Radar profiling of conjugate levels showed that the 4 μM treatment produced the most pronounced and uniform reduction across the measured IAA conjugates, particularly IAA-Leu, IAA-Phe and IAA-Tyr (Fig. 6e). Consistent with this, within-group R² values were highest at 4 μM across most indicators, including free IAA (Fig. S67), indicating that this concentration exerted the most consistent and predictable effect on auxin redistribution. A composite impact score (S_d), calculated as the sum of absolute standardized deviations from the control across all measured indicators, confirmed that 4 μM yielded the highest overall departure from control conditions, followed by 8 μM and 2 μM, while 1 μM showed the weakest aggregate effect (Fig. S68).

## 4 Discussion

This study resolves the *S. chinensis* and *R. chinensis* galling system at a level that connects genomic architecture with effector function. The paired chromosome-level assemblies establish the structural context in which the aphid operates, the morph-resolved secretome and metabolome identify candidate effectors and their chemical environment, and functional assays link one of those candidates, ACY1, to a measurable shift in host auxin homeostasis and to galling success. The most striking feature of the aphid genome is the extreme expansion of chromosome 1. At 93.61 Mb, chr1 carries 33.08% of all anchored genes and concentrates 70.47% of genome-wide SDs, along with 58.87% of all genome-wide TE-related genes residing on this single chromosome. Syntenic comparisons with *V. vitifoliae* and *A. pisum* show that chr1 contains material corresponding to several ancestral chromosomal elements, a pattern compatible with fusion, translocation, or progressive accretion, though the present data do not discriminate among these alternatives. What is directly supported is the current mode of expansion. The concentration of SDs, tandem arrays and repeat-associated genes on chr1 indicates that this chromosome has grown and continues to grow primarily through ongoing duplication and repeat accumulation rather than through a single historical rearrangement [31–33].

The functional organisation of chr1 reinforces this view. Stress-responsive chaperones (all nine crystallin/sHSP20 genes genome-wide), a dense opsin cluster spanning roughly 240 kb with four spectral classes, PIF1 helicases and SETMAR-associated loci are not randomly scattered across the genome but concentrated on this single oversized chromosome. The co-localisation of environmental sensing, proteostasis and genome-maintenance functions within a structural domain that has itself been heavily shaped by duplication is consistent with an amplification-driven mode of adaptive innovation[34, 35]. At the same time, chr1 carries a disproportionately low share of positively selected genes (13.31% versus 33.08% of anchored genes), suggesting that sequence-level amino-acid diversification has not been the dominant route to innovation on this chromosome. The data are therefore more consistent with a dosage- and architecture-based model, in which tandem amplification and repeat-associated restructuring generate adaptive potential, than with a conventional positive-selection-centred view of chr1 evolution.

A related finding that deserves emphasis is the reinterpretation of the lysine degradation signal on chr1. This pathway appeared as the strongest KEGG enrichment for chr1-resident genes, but manual re-annotation revealed that most of the genes contributing to this signal are Mariner-derived sequences or SETMAR fusions rather than bona fide catabolic enzymes. This result has two implications. First, it illustrates how automated pathway annotation can be misleading when applied to a chromosome with heavy transposon load, and it underscores the need for careful manual curation in repeat-rich genomic regions. Second, it points to the coupled amplification of transposon-derived sequences and chromatin-associated regulators as a distinctive feature of chr1 biology. Chr1 is therefore less a generic metabolic chromosome than one on which repeat load and adaptive innovation have become unusually entangled[36].

An important contextual factor is the holocentric nature of aphid chromosomes[37–39]. In holocentric organisms, kinetochore activity is distributed along the entire length of the chromosome rather than localised to a single centromere. This property allows chromosomal fragments to segregate normally during cell division, which in turn means that structural rearrangements, fissions, fusions and large-scale insertions can be tolerated without the mitotic instability that would result in monocentric systems [40–43]. The extreme size asymmetry of chr1 should be understood against this background. A holocentric chromosome can, in principle, accumulate duplications and repeat insertions without a proportional increase in segregation errors, and this permissiveness may help explain why one chromosome has become so disproportionately enlarged in *S. chinensis*. It remains an open question whether similar expansion biases exist in other holocentric galling lineages, but the prediction follows naturally from the mechanics of holocentric mitosis.

The chromosomal architecture recovered here differs from the two previously published *S. chinensis* genome assemblies, both of which resolved 13 chromosomes[44, 45], whereas our assembly resolves 16, a count independently confirmed by DAPI-FISH and Giemsa staining (2n = 32). Earlier cytological compilations listed this species at approximately 2n = 36, indicating that the karyotype has been under-resolved in prior genome resources. Given the holocentric nature of aphid chromosomes and the well-documented tendency of aphid autosomes to undergo extensive rearrangement [40, 43], the most plausible explanation is that repeat-rich or structurally complex regions were merged into fewer pseudochromosomes in earlier assemblies. This issue is especially relevant here because a genome dominated by a duplication-rich chr1 would be particularly susceptible to collapsed chromosome boundaries if long repetitive segments were incorrectly joined. By contrast, the host *R. chinensis* karyotype is concordant across studies, with 15 chromosome pairs (2n = 30) confirmed independently by our cytology and by a recent assembly [46].

The host genome complements the aphid picture in a different way. It lacks a structurally dominant chromosome, but it is enriched for gene families that are highly relevant to the aphid-plant interface, including pectin methylesterases and other cell-wall remodelling enzymes, CYP450s, GSTs, UGTs and ABC transporters [47, 48]. Positive selection was detected in plant hormone signal transduction and plant–pathogen interaction pathways, indicating that the host genome has been shaped by sustained selective pressure at the interaction interface. These features do not by themselves reveal how individual host cell types respond during galling, but they define a genomic background that is well equipped for both tissue restructuring and chemical defence. In that sense, the host genome provides the response landscape on which aphid effectors must operate, rather than constituting a separate story within this study.

ACY1 provides the clearest functional entry point into that landscape. Several independent lines of evidence converge on it. The proteomic data place ACY1 in gall-associated saliva, with elevated abundance in fundatrigenia and fundatrix relative to non-galling sexuparae. The metabolomic data show that gall-associated saliva is enriched in free amino acids, N-acetyl amino acids and other aminoacyl-related compounds that are consistent with M20 metallopeptidase substrates. RNAi-mediated knockdown of *ACY1* reduced galling success from roughly 90% to 52.33%, increased aphid mortality, lowered free IAA and caused all seven measured IAA conjugates to accumulate at the settlement site. Cell-free delivery of active ACY1 produced the reciprocal pattern in a dose-dependent manner, with free IAA increasing and several conjugates declining, and the 4 μM concentration yielding the strongest and most consistent effects across all measured indicators. The direction and internal consistency of these reciprocal results, loss-of-function and gain-of-function, support a model in which ACY1 mobilises host auxin from conjugated storage forms into the free pool. Among the seven species measured, the ester-linked IAA-Glc is not a substrate for an amide hydrolase, and its change is therefore interpreted as an indirect consequence of the shift in free-IAA balance rather than direct cleavage.

The conceptual significance of this mechanism lies in its distinction from other proposed routes to elevated auxin in gall tissues. In many galling systems, altered IAA levels have been documented at the gall site, but the source of that auxin has remained ambiguous [49, 48, 50].

One possibility is *de novo* synthesis by the insect, and indeed some arthropods carry endogenous auxin biosynthetic capacity [51]. Another is transcriptional upregulation of host biosynthesis genes, as has been observed in cynipid wasp galls [52]. A third possibility, supported by the present data, is that the insect directly liberates active auxin from the host plant’s own conjugated reservoir. In plants, reversible conjugation of IAA to amino acids serves as a buffering mechanism that controls the pool of active hormone, with GH3 family enzymes catalysing conjugation and amidohydrolases catalysing release [20, 53, 54]. The evidence presented here suggests that *S. chinensis* has co-opted this buffering system by secreting a salivary amidohydrolase that tips the balance towards free IAA at the feeding site.

This strategy is attractive for a long-lived galling interaction for at least two reasons. First, it is metabolically economical. Rather than synthesising and continuously injecting exogenous hormone, the aphid uses a catalytic enzyme to unlock a pre-existing and renewable substrate pool within the host. A small amount of secreted enzyme can, in principle, liberate a much larger amount of active auxin from the conjugated reservoir. Second, the mechanism is inherently spatially restricted. Because the enzyme is delivered through the stylet at the feeding site, auxin release is confined to the immediate settlement zone, consistent with the localised cell expansion and tissue remodelling observed histologically. How this initial local auxin pulse is translated into the sustained, three-dimensional growth of a mature horned gall remains an open question. Auxin is subject to polar transport and feedback regulation in plant tissues [20], and additional effectors are almost certainly involved in shaping the full gall organ. The proteomic data identified several other M20-family members and amidohydrolase candidates in gall-associated saliva [55, 56], including FAAH2, CNDP2-1 and CNDP2-2, and their possible contributions to gall maintenance have not yet been tested.

The auxin deconjugation mechanism described here can be placed alongside other recently characterised modes of insect-mediated host manipulation. In the Hessian fly *Mayetiola destructor*, a massively expanded family of secreted effector genes has been implicated in gall formation, though the biochemical targets of most of these effectors remain unknown [57]. In *Drosophila*-associated gall midges, a novel family of secreted proteins has been linked to gall development, again without a defined enzymatic mechanism [58]. More recently, secreted CAP peptides from gall-inducing sawflies have been shown to induce gall-like growth when applied exogenously to plants[9]. The present study adds a different dimension to this picture by identifying not just a secreted protein associated with galling, but a specific biochemical activity, auxin deconjugation, that can account for a measurable shift in host hormone pools. The parallels with pathogen systems are also worth noting. Root-knot nematodes manipulate host auxin transport through PIN protein redistribution [59], and several bacterial and fungal pathogens produce or manipulate auxin to facilitate infection [60]. The strategy used by *S. chinensis* differs from all of these in that it targets the host’s conjugated auxin reservoir directly, rather than modulating biosynthesis, transport or signalling.

A broader question raised by this work is how the genomic architecture of chr1 relates to the evolution of effectors like ACY1. The data show that chr1 has been a major site of duplication-driven gene-family expansion and that ACY1 is among the effector candidates identified in the salivary secretome. However, the present data do not establish a direct mechanistic link between chr1 expansion and the functional diversification of any individual effector. The apparent decoupling between copy-number expansion and positive selection on chr1 argues against a simple model in which chr1 is the primary locus of sequence-level adaptive evolution, but it does not by itself reveal the roles of gene conversion, local recombination architecture, or epigenetic regulation in shaping the effector landscape. Tracing the evolutionary origin and duplication history of individual effector loci relative to the broader pattern of segmental duplication on chr1 is a focused follow-up question that will require population-genomic sampling across related galling and non-galling aphid species.

Taken together, this study supports a model in which a gall-inducing aphid manipulates host development through a salivary enzyme that shifts auxin from conjugated storage forms into the active pool, and in which this capacity has evolved within a genome marked by an anomalously expanded chromosome that concentrates duplication and repeat-associated innovation. These two features, a specific biochemical mechanism of host developmental control and a distinctive mode of genome evolution, can now be investigated in the same system. Whether auxin deconjugation by salivary amidohydrolases is a general strategy among galling aphids, and whether the chromosomal gigantism observed here reflects a broader pattern in holocentric galling lineages, are testable predictions that follow directly from the present data.

## Acknowledgments

This work was supported by grants from the Academician Expert Workstation in Yunnan Province (202405AF140107) and the Science and Technology Department of Yunnan Province (202449CE340005), Xingdian Talent Support Program (yfgrc202525).

## Author contributions

Hang Chen conceived and designed the study and supervised the project. Qin Lu performed the genome sequencing, assembly and annotation of *S. chinensis* and *c*. Weiwei Wang and Juan Liu conducted the morph-resolved salivary proteomics and metabolomics. Xumei Chen and Yiyuan Pan performed the RNA interference and cell-free enzyme assays and the in planta auxin profiling. Qin Lu, Xin Zhang and Chenjing Ma analysed the data and prepared the figures. Hang Chen and Qin Lu wrote the manuscript with input from all authors. All authors read and approved the final manuscript.

## Data availability

The *Rhus chinensis* chromosome-level genome assemblies have been deposited in the Genome Dataset Center of NCBI under BioProject PRJNA1443190. The assembled genomes of *S. chinensis* are available under accession PRJNA1443548. The mass spectrometry proteomics data have been deposited to the ProteomeXchange Consortium via the PRIDE partner repository with the dataset identifier PXD079224. The metabolomics dataset generated in this study has been deposited in the MetaboLights database (https://www.ebi.ac.uk/metabolights/) under accession number MTBLS14665. All other data supporting the findings of this study are available within the article and its Supplementary materials, or from the corresponding author upon reasonable request.

## References

[1] Desnitskiy AG, Chetverikov PE, Ivanova LA, et al. Molecular aspects of gall formation induced by mites and insects. Life 2023;13:1347.

[2] Markel K, Novak V, Bowen BP, et al. Cynipid wasps systematically reprogram host metabolism and restructure cell walls in developing galls. Plant Physiol. 2024;195:698–712.

[3] Bellows E, Heatley M, Shah N, et al. Comparative transcriptome reprogramming in oak galls containing asexual or sexual generations of gall wasps. Plant Biol. 2024;26:798–810.

[4] Kutsukake M, Uematsu K, Fukatsu T. Plant manipulation by gall-forming social aphids for waste management. Front. Plant Sci. 2019;10:933.

[5] Chen X, Yang Z, Chen H, et al. A complex nutrient exchange between a gall-forming aphid and its plant host. Front. Plant Sci. 2020;11:811.

[6] Hirano T, Kimura S, Sakamoto T, et al. Reprogramming of the developmental program of Rhus javanica during initial stage of gall induction by Schlechtendalia chinensis. Front. Plant Sci. 2020;11:471.

[7] Huang C, Ji B, Shi Z, et al. A comparative genomic analysis at the chromosomal-level reveals evolutionary patterns of aphid chromosomes. Commun. Biol. 2025;8:427.

[8] Giron D, Huguet E, Stone GN, et al. Insect-induced effects on plants and possible effectors used by galling and leaf-mining insects to manipulate their host-plant. J. Insect Physiol. 2016;84:70–89.

[9] Hirano T, Sakamoto T, Kimura S, et al. CAP peptides artificially induce insect-gall-like growth in different plant species. Plant Cell Physiol. 2025;66:1155–68.

[10] Stern DL, Han C. Gene structure-based homology search identifies highly divergent putative effector gene family. Genome Biol. Evol. 2022;14:evac069.

[11] Tokuda M, Suzuki Y, Fujita S, et al. Terrestrial arthropods broadly possess endogenous phytohormones auxin and cytokinins. Sci. Rep. 2022;12:4750.

[12] Yamaguchi H, Tanaka H, Hasegawa M, et al. Phytohormones and willow gall induction by a gall-inducing sawfly. New Phytol. 2012;196:586–95.

[13] Mapes CC, Davies PJ. Indole-3-acetic acid and ball gall development on Solidago altissima. New Phytol. 2001;151:195–202.

[14] Body MJA, Schultz JC, Appel HM, et al. A gall-forming insect manipulates hostplant phytohormone synthesis, concentrations, and signaling. New Phytol. 2019;224:836–52.

[15] Nabity PD, Haus MJ, Berenbaum MR, et al. Leaf-galling phylloxera on grapes reprograms host metabolism and morphology. Proc. Natl. Acad. Sci. U. S. A. 2013;110:16663–8.

[16] Grunewald W, Cannoot B, Friml J, et al. Parasitic nematodes modulate PIN-mediated auxin transport to facilitate infection. PLoS Pathog. 2009;5:e1000266.

[17] Casanova-Sáez R, Mateo-Bonmatí E, Ljung K. Auxin Metabolism in Plants. Cold Spring Harb. Perspect. Biol. 2021;13:a039867.

[18] Sanchez Carranza AP, Singh A, Steinberger K, et al. Hydrolases of the ILR1-like family of Arabidopsis thaliana modulate auxin response by regulating auxin homeostasis in the endoplasmic reticulum. Sci. Rep. 2016;6:24212.

[19] Casanova-Sáez R, Voß U. Auxin Metabolism Controls Developmental Decisions in Land Plants. Trends Plant Sci. 2019;24:741–54.

[20] Prinsen E. Auxin homeostasis: new roads in a tight network. J. Exp. Bot. 2025;76:3260–2.

[21] Cheng H, Asri M, Lucas J, et al. Scalable telomere-to-telomere assembly for diploid and polyploid genomes with double graph. Nat. Methods 2024;21:967–70.

[22] Cheng H, Concepcion GT, Feng X, et al. Haplotype-resolved de novo assembly using phased assembly graphs with hifiasm. Nat. Methods 2021;18:170–5.

[23] Durand NC, Shamim MS, Machol I, et al. Juicer provides a one-click system for analyzing loop-resolution Hi-C experiments. Cell Syst. 2016;3:95–8.

[24] Dudchenko O, Batra SS, Omer AD, et al. De novo assembly of the Aedes aegypti genome using Hi-C yields chromosome-length scaffolds. Science 2017;356:92–5.

[25] Manni M, Berkeley MR, Seppey M, et al. BUSCO update: novel and streamlined workflows along with broader and deeper phylogenetic coverage for scoring of eukaryotic, prokaryotic, and viral genomes. Mol. Biol. Evol. 2021;38:4647–54.

[26] Emms DM, Kelly S. OrthoFinder: phylogenetic orthology inference for comparative genomics. Genome Biol. 2019;20:238.

[27] Mendes FK, Vanderpool D, Fulton B, et al. CAFE 5 models variation in evolutionary rates among gene families. Bioinformatics 2020;36:5516–8.

[28] Yang Z. PAML 4: phylogenetic analysis by maximum likelihood. Mol. Biol. Evol. 2007;24:1586–91.

[29] Wang Y, Tang H, DeBarry JD, et al. MCScanX: a toolkit for detection and evolutionary analysis of gene synteny and collinearity. Nucleic Acids Res. 2012;40:e49.

[30] Cox J, Mann M. MaxQuant enables high peptide identification rates, individualized p.p.b.-range mass accuracies and proteome-wide protein quantification. Nat. Biotechnol. 2008;26:1367–72.

[31] Ruckman SN, Jonika MM, Casola C, et al. Chromosome number evolves at equal rates in holocentric and monocentric clades. PLoS Genet. 2020;16:e1009076.

[32] Mandrioli M, Manicardi GC. Holocentric chromosomes. PLoS Genet. 2020;16:e1008918.

[33] Mathers TC, Wouters RHM, Mugford ST, et al. Chromosome-scale genome assemblies of aphids reveal extensively rearranged autosomes and long-range gene regulation. Mol. Biol. Evol. 2021;38:856–75.

[34] Hahn MW. Distinguishing among evolutionary models for the maintenance of gene duplicates. J. Hered. 2009;100:605–17.

[35] Kondrashov FA. Gene duplication as a mechanism of genomic adaptation to a changing environment. Proc. R. Soc. B Biol. Sci. 2012;279:5048–57.

[36] Ahmad A, Ren Z. Mobilome of the Rhus gall aphid Schlechtendalia chinensis provides insight into TE insertion-related inactivation of functional genes. Int. J. Mol. Sci. 2022;23:15967.

[37] Mandrioli M, Zambonini G, Manicardi GC. Comparative gene mapping as a tool to understand the evolution of pest crop insect chromosomes. Int. J. Mol. Sci. 2017;18:1919.

[38] Hofstatter PG, Thangavel G, Lux T, et al. Repeat-based holocentromeres influence genome architecture and karyotype evolution. Cell 2022;185:3153–3168.e18.

[39] Escudero M, Marques A, Lucek K, et al. Genomic hotspots of chromosome rearrangements explain conserved synteny despite high rates of chromosome evolution in a holocentric lineage. Mol. Ecol. 2024;33:e17086.

[40] Jing T, Yang J, Pan J, et al. A near-complete genome reveals the population evolution of the cotton-melon aphid Aphis gossypii. Insect Biochem. Mol. Biol. 2025;176:104215.

[41] Mata-Sucre Y, Krátká M, Oliveira L, et al. Repeat-based holocentromeres of the woodrush Luzula sylvatica reveal insights into the evolutionary transition to holocentricity. Nat. Commun. 2024;15:9565.

[42] Márquez-Corro JI, Martín-Bravo S, Blanco-Pastor JL, et al. The holocentric chromosome microevolution: From phylogeographic patterns to genomic associations with environmental gradients. Mol. Ecol. 2024;33:e17156.

[43] Albuquerque L, Milani D, Martí E, et al. Exploring the Satellitome of the Pest Aphid Acyrthosiphon pisum (Hemiptera, Aphididae): Insights Into Genome Organization and Intraspecies Evolution. Genome Biol. Evol. 2025;17:evaf104.

[44] Ahmad A, von Dohlen C, Ren Z. A chromosome-level genome assembly of the Rhus gall aphid Schlechtendalia chinensis provides insight into the endogenization of Parvovirus-like DNA sequences. BMC Genomics 2024;25:16.

[45] Wei H-Y, Ye Y-X, Huang H-J, et al. Chromosome-level genome assembly for the horned-gall aphid provides insights into interactions between gall-making insect and its host plant. Ecol. Evol. 2022;12:e8815.

[46] Lu Z, Zou H, Cui J, et al. The Rhus chinensis Genome Provides Insights Into Tannin, Flavonoid Biosynthesis, and Glandular Trichome Development. Plant Biotechnol. J. 2026;24:988–1013.

[47] Fan B-L, Chen L-H, Chen L-L. Analysis of gene expansion and defense-related genes in Anacardiaceae family from an evolutionary aspect. Front. Plant Sci. 2025;16:1638044.

[48] Ma Y, Şen ZB, Ng HP. Plant galls induced by insects: Coordinated developmental reprogramming and defence manipulation. Curr. Opin. Plant Biol. 2025;86:102757.

[49] Connor EF. Insect Production and Secretion of Phytohormones and Impacts on Host Plants. Annu. Rev. Entomol. 2026;71:427–46.

[50] Boo K-H, Oh YK, Møller C, et al. Dasineura asteriae Reprograms the Flower Gene Expressions of Vegetative Organs to Create Flower-Like Gall in Aster scaber. Plant Cell Environ. 2025;48:8217–31.

[51] Natahusada J, Roy SW, Connor EF. A bioinformatic examination of indole-3-acetic acid biosynthesis in insecta and hexapoda. Arthropod-Plant Interact. 2025;19:4.

[52] Wang Y, Xue C, Wu S, et al. Transcriptome and phytohormone analysis reveal mechanism of gall formation by Trichagalma acutissimae larvae on oak leaves. Front. Plant Sci. 2025;16:1646230.

[53] Luo P, Li T-T, Shi W-M, et al. The Roles of GRETCHEN HAGEN3 (GH3)-Dependent Auxin Conjugation in the Regulation of Plant Development and Stress Adaptation. Plants 2023;12:4111.

[54] Zheng H, Zhang Q, Liu Q, et al. Auxin Biosynthesis, Transport, Signaling, and Its Roles in Plant Leaf Morphogenesis. Plants 2025;15:72.

[55] Bleau JR, Gaur N, Fu Y, et al. Unveiling the slippery secrets of saliva: effector proteins of phloem-feeding insects. Mol. Plant. Microbe Interact. 2024;37:211–9.

[56] Portillo Lemus L, Tricard J, Duclercq J, et al. Salivary proteins of Phloeomyzus passerinii, a plant-manipulating aphid, and their impact on early gene responses of susceptible and resistant poplar genotypes. Plant Sci. 2020;294:110468.

[57] Zhao C, Escalante LN, Chen H, et al. A massive expansion of effector genes underlies gall-formation in the wheat pest Mayetiola destructor. Curr. Biol. 2015;25:613–20.

[58] Korgaonkar A, Han C, Lemire AL, et al. A novel family of secreted insect proteins linked to plant gall development. Curr. Biol. 2021;31:1836–49.

[59] Yadav H, Roberts PA, Lopez-Arredondo D. Combating Root-Knot Nematodes (Meloidogyne spp.): From Molecular Mechanisms to Resistant Crops. Plants 2025;14:1321.

[60] Duhan L, Pasrija R. Unveiling exogenous potential of phytohormones as sustainable arsenals against plant pathogens: molecular signaling and crosstalk insights. Mol. Biol. Rep. 2025;52:98.

