## Supplementary figures and tables for "Chromosome gigantism and auxin deconjugation underpin gall induction in a horned gall aphid"

#### Supplementary tables

**Table S1.** Primers used for RT-qPCR

| Target gene | Primer name | Sequence (5′–3′) | Amplicon size (bp) |
|---|---|---|---|
| *FAAH2* | Forward | TTGCGGTAGTCCTTTGGGTC | 166 |
| | Reverse | AGGTCCAACGCTAACCATGG | |
| *ACY1* | Forward | ATGGCTCGGACAGAAACCAG | 150 |
| | Reverse | ATGTCTTGTGCGCCTCTAGC | |
| *CNDP2-1* | Forward | CTGTTCCAGCTCCGCTAGTC | 119 |
| | Reverse | TCTCAAATCTGGGTCAGCCG | |
| *CNDP2-2* | Forward | CTGTGTTCCATTGTTCCCGC | 154 |
| | Reverse | TCTCAAATCTGGATCAGCCGA | |
| *ADL* | Forward | AGGAGAACCATCTAGTGTTGCG | 176 |
| | Reverse | GTCCATCAGACGTACCTCCG | |
| *GatA* | Forward | TCGTCGGGAAGACCAACTTG | 133 |
| | Reverse | CTTCCACCCGAACTACCACC | |
| *$\beta$-actin* | Forward | CTCTGTCTGGATTGGAGGGTC | 125 |
| | Reverse | GGCTTGAGAAATGGTCGGA | |

**Table S2.** *S. chinensis* genome assembly statistics at the contig level

| Parameter | This study | Reported genome (Wei et al., 2021) |
|---|---|---|
| Number of contigs | 1,710 | 1,409 |
| Total length (bp) | 337,848,735 | 304,774,269 |
| Largest contig (bp) | 16,419,613 | – |
| Contig N50 (bp) | 5,496,496 | 2,961,835 |
| Contig N75 (bp) | 2,126,621 | – |
| Contig N90 (bp) | 80,426 | 115,058 |
| Contigs ≥ 5,000 bp | 1,578 | – |

| | | |
|---|---|---|
| Contigs ≥ 10,000 bp | 1,224 | – |
| Contigs ≥ 25,000 bp | 703 | – |
| Contigs ≥ 50,000 bp | 339 | – |
| Chromosome-anchored length (bp) | 305,098,848 | – |
| Anchoring rate (%) | 90.31 | – |
| Genome size (Mb) | 297.95 | 304.77 |
| GC content (%) | 34.64 | – |
| Repeat content (%) | 42.91 | – |
| Protein-coding genes | 19,109 | – |
| Chromosome number (2n) | 32 | – |

**Table S3.** Size of each *S. chinensis* chromosome

| Chromosome | Size (bp) | Gene count | % of total genes |
|---|---|---|---|
| chr1 | 95,852,465 | 5,986 | 33.08 |
| chr2 | 21,522,562 | 1,007 | 5.56 |
| chr3 | 21,223,707 | 1,073 | 5.93 |
| chr4 | 21,089,406 | 1,314 | 7.26 |
| chr5 | 20,837,838 | 962 | 5.32 |
| chr6 | 20,650,788 | 972 | 5.37 |
| chr7 | 14,817,943 | 858 | 4.74 |
| chr8 | 13,420,551 | 665 | 3.67 |
| chr9 | 12,597,102 | 685 | 3.78 |
| chr10 | 12,068,285 | 705 | 3.90 |
| chr11 | 11,695,868 | 750 | 4.14 |
| chr12 | 11,206,492 | 626 | 3.46 |
| chr13 | 10,121,268 | 693 | 3.83 |
| chr14 | 7,470,798 | 518 | 2.86 |
| chr15 | 6,050,636 | 659 | 3.64 |
| chr16 | 4,473,139 | 623 | 3.44 |

**Table S4.** Chromosome quantity and genome size of seven aphid species

| Species | Genome size (Mb) | GC (%) | Chromosome number |
|---|---|---|---|
| *Acyrthosiphon pisum* | 465.15 | 30.50 | 2n = 8 |
| *Aphis gossypii* | 299.57 | 27.53 | 2n = 8 |
| *Myzus persicae* | 347.31 | 30.01 | – |
| *Schizaphis graminum* | 494.64 | 28.40 | 2n = 12 |
| *Viteus vitifoliae* | 282.59 | 26.60 | – |
| *Eriosoma lanigerum* | 334.87 | 26.37 | 2n = 12 |
| *S. chinensis* | 305.10 | 34.63 | 2n = 32 |

**Table S5.** Repeat content in the *S. chinensis* genome

| Method | Repeat size (bp) | % of genome |
|---|---|---|
| TRF (tandem repeats) | 13,386,947 | 3.96 |
| RepeatMasker | 8,292,167 | 2.45 |
| RepeatProteinMask | 14,399,500 | 4.26 |
| De novo (RepeatModeler + LTR-FINDER) | 136,254,511 | 40.31 |
| Total (non-redundant) | 145,046,207 | 42.91 |

**Table S6.** Types of repetitive elements in the *S. chinensis* genome

| Method | DNA transposons (bp) | LINE (bp) | LTR (bp) | SINE (bp) | Total (bp) | Unknown (bp) |
|---|---|---|---|---|---|---|
| RepeatMasker | 4,926,678 | 1,359,291 | 2,058,223 | 1,714 | 8,228,248 | 418,740 |
| RepeatProteinMask | 4,213,556 | 4,975,580 | 5,214,534 | 0 | 14,399,500 | 576 |
| De novo | 41,629,335 | 27,203,571 | 45,061,632 | 15,791 | 134,699,806 | 37,131,871 |
| Combined (non-redundant) | 45,849,613 | 29,186,535 | 46,174,853 | 17,505 | 140,566,308 | 37,541,860 |

**Table S7.** Gene prediction of *S. chinensis* compared with related species

| Gene set | Number | Avg gene length (bp) | Avg CDS length (bp) | Avg exon number | Avg exon length (bp) | Avg intron length (bp) |
|---|---|---|---|---|---|---|
| AUGUSTUS | 27,733 | 5,301.72 | 1,267.46 | 4.76 | 266.39 | 1,073.52 |
| GlimmerHMM | 26,883 | 10,702.52 | 776.86 | 4.69 | 165.63 | 2,689.56 |
| Genscan | 22,495 | 8,757.44 | 1,261.97 | 5.52 | 228.69 | 1,658.95 |
| RNAseq | 6,426 | 10,825.29 | 1,295.52 | 7.63 | 330.35 | 1,253.06 |
| *A. pisum* | 19,968 | 5,164.39 | 1,078.25 | 4.51 | 239.23 | 1,165.07 |
| *A. gossypii* | 19,245 | 5,663.65 | 1,140.92 | 4.77 | 239.18 | 1,199.59 |
| *M. persicae* | 19,464 | 5,336.60 | 1,094.05 | 4.63 | 236.04 | 1,167.15 |
| *S. graminum* | 21,281 | 4,310.56 | 953.06 | 4.15 | 229.68 | 1,066.02 |
| *V. vitifoliae* | 16,615 | 5,925.48 | 1,158.58 | 5.04 | 229.93 | 1,180.29 |
| *E. lanigerum* | 24,098 | 3,170.28 | 909.39 | 3.79 | 239.76 | 809.49 |
| *S. chinensis* | 19,109 | 6,691.85 | 1,346.68 | 5.62 | 286.31 | 1,098.80 |

**Table S8.** BUSCO assessment of *S. chinensis* and *R. chinensis* genome assemblies

| Parameter | *S. chinensis* | *R. chinensis* |
|---|---|---|
| Complete BUSCOs | 2,458 (98.0%) | 1,572 (97.39%) |
| Complete single-copy | 2,369 (94.4%) | 1,444 (89.47%) |
| Complete duplicated | 34 (1.4%) | 38 (2.35%) |
| Fragmented | 12 (0.5%) | 16 (0.99%) |
| Missing | 40 (1.5%) | 26 (1.61%) |
| Total BUSCO groups | 2,510 (100%) | 1,614 (100%) |

**Table S9.** GC content of each *S. chinensis* chromosome

| Chromosome | GC (%) |
|---|---|
| chr1 | 31.48 |
| chr2 | 34.08 |
| chr3 | 34.04 |
| chr4 | 34.34 |
| chr5 | 34.57 |
| chr6 | 34.59 |
| chr7 | 31.91 |
| chr8 | 36.35 |
| chr9 | 37.11 |

| | |
|---|---|
| chr10 | 36.64 |
| chr11 | 36.88 |
| chr12 | 34.76 |
| chr13 | 38.02 |
| chr14 | 32.80 |
| chr15 | 33.62 |
| chr16 | 33.02 |

**Table S10.** Functional annotation of the *S. chinensis* genome by database

| Database | Number | Percent (%) |
|---|---|---|
| Total | 19,109 | 100.00 |
| InterPro | 12,033 | 62.97 |
| GO | 5,996 | 31.38 |
| KEGG | 17,172 | 89.86 |
| KEGG_KO | 7,061 | 36.95 |
| Swissprot | 10,921 | 57.15 |
| TrEMBL | 17,260 | 90.32 |
| NR | 17,499 | 91.57 |
| Annotated (any database) | 17,586 | 92.03 |
| Non-annotated | 1,523 | 7.97 |

**Table S11.** Annotation of non-coding RNA in *S. chinensis*

| Type | Subtype | Copy | Average length (bp) | Total length (bp) | % of genome |
|---|---|---|---|---|---|
| miRNA | | 27 | 86.74 | 2,342 | 0.0007 |
| tRNA | | 145 | 75.60 | 10,962 | 0.0032 |
| rRNA | rRNA | 101 | 3,175.56 | 320,732 | 0.0949 |
| | 18S | 35 | 2,511.97 | 87,919 | 0.0260 |
| | 28S | 42 | 5,477.02 | 230,035 | 0.0680 |
| | 5S | 24 | 115.75 | 2,778 | 0.0008 |
| snRNA | snRNA | 77 | 152.69 | 11,757 | 0.0035 |
| | CD-box | 6 | 124.83 | 749 | 0.0002 |
| | HACA-box | 0 | 0 | 0 | 0 |
| | splicing | 71 | 155.04 | 11,008 | 0.0033 |
| | scaRNA | 0 | 0 | 0 | 0 |

**Table S12.** BUSCO assessment of *S. chinensis* genome assembly and gene annotation

| Category | Assembly Proteins | Assembly % | Annotation Proteins | Annotation % |
|---|---|---|---|---|
| Complete BUSCOs | 2,458 | 98.0 | 2,369 | 94.4 |
| Complete single- | 2,424 | 96.6 | 2,308 | 92.0 |

| | | | | |
|---|---|---|---|---|
| copy BUSCOs | | | | |
| Complete duplicated BUSCOs | 34 | 1.4 | 61 | 2.4 |
| Fragmented BUSCOs | 12 | 0.5 | 21 | 0.8 |
| Missing BUSCOs | 40 | 1.5 | 120 | 4.8 |
| Total BUSCO groups | 2,510 | 100 | 2,510 | 100 |

**Table S13.** Number of duplicated genes on each *S. chinensis* chromosome

| Chr | Dispersed | Proximal | Tandem | SDs | % SDs | Total duplicated | % Total |
|---|---|---|---|---|---|---|---|
| chr1 | 1,213 | 237 | 197 | 3,353 | 70.47 | 5,000 | 39.15 |
| chr2 | 440 | 30 | 90 | 6 | 0.13 | 566 | 4.43 |
| chr3 | 508 | 86 | 121 | 87 | 1.83 | 802 | 6.28 |
| chr4 | 541 | 87 | 97 | 20 | 0.42 | 745 | 5.83 |
| chr5 | 467 | 57 | 75 | 74 | 1.56 | 673 | 5.27 |
| chr6 | 486 | 62 | 85 | 46 | 0.97 | 679 | 5.32 |
| chr7 | 415 | 82 | 72 | 6 | 0.13 | 575 | 4.50 |
| chr8 | 348 | 32 | 45 | 13 | 0.27 | 438 | 3.43 |
| chr9 | 303 | 45 | 62 | 0 | 0 | 410 | 3.21 |
| chr10 | 302 | 47 | 62 | 32 | 0.67 | 443 | 3.47 |
| chr11 | 259 | 53 | 79 | 22 | 0.46 | 413 | 3.23 |
| chr12 | 285 | 34 | 46 | 7 | 0.15 | 372 | 2.91 |
| chr13 | 296 | 27 | 52 | 17 | 0.36 | 392 | 3.07 |
| chr14 | 34 | 17 | 11 | 366 | 7.69 | 428 | 3.35 |
| chr15 | 47 | 16 | 16 | 560 | 11.77 | 639 | 5.00 |
| chr16 | 25 | 11 | 12 | 149 | 3.13 | 197 | 1.54 |

**Table S14.** Number of TE-related genes on each *S. chinensis* chromosome

| Chromosome | TE related genes | Percentage |
|---|---|---|
| chr1 | 1,682 | 58.87% |
| chr2 | 50 | 1.75% |
| chr3 | 101 | 3.54% |
| chr4 | 54 | 1.89% |
| chr5 | 77 | 2.70% |
| chr6 | 61 | 2.14% |
| chr7 | 77 | 2.70% |
| chr8 | 35 | 1.23% |
| chr9 | 51 | 1.79% |
| chr10 | 72 | 2.52% |
| chr11 | 49 | 1.72% |
| chr12 | 21 | 0.74% |
| chr13 | 28 | 0.98% |
| chr14 | 184 | 6.44% |
| chr15 | 221 | 7.74% |
| chr16 | 94 | 3.29% |

**Table S15.** Gene family statistics for aphid species

| Species | Family number | Unique families | Average genes per family |
|---|---|---|---|
| *S. chinensis* | 9,668 | 295 | 1.817 |
| *E. lanigerum* | 11,346 | 734 | 1.966 |
| *V. vitifoliae* | 9,091 | 265 | 1.566 |
| *A. pisum* | 10,395 | 108 | 1.668 |
| *A. craccivora* | 12,556 | 461 | 2.225 |
| *A. gossypii* | 10,115 | 57 | 1.455 |
| *M. persicae* | 10,207 | 34 | 1.454 |
| *S. graminum* | 12,245 | 603 | 1.781 |
| *B. tabaci* | 8,730 | 417 | 1.439 |
| *E. pela* | 5,240 | 297 | 1.502 |
| *N. lugens* | 9,434 | 589 | 1.743 |

**Table S16.** Number of expanded, contracted, and positively selected genes on each *S. chinensis* chromosome

| Chromosome | Expanded genes | % expanded | Contracted genes | % contracted | PSG genes | % PSG |
|---|---|---|---|---|---|---|
| chr1 | 2,806 | 51.86 | 102 | 45.13 | 41 | 13.31 |
| chr2 | 92 | 1.70 | 27 | 11.95 | 23 | 7.47 |
| chr3 | 291 | 5.38 | 11 | 4.87 | 28 | 9.09 |
| chr4 | 128 | 2.37 | 4 | 1.77 | 39 | 12.66 |
| chr5 | 196 | 3.62 | 7 | 3.10 | 27 | 8.77 |
| chr6 | 131 | 2.42 | 7 | 3.10 | 34 | 11.04 |
| chr7 | 233 | 4.31 | 8 | 3.54 | 10 | 3.25 |
| chr8 | 75 | 1.39 | 3 | 1.33 | 21 | 6.82 |
| chr9 | 101 | 1.87 | 8 | 3.54 | 20 | 6.49 |
| chr10 | 127 | 2.35 | 11 | 4.87 | 14 | 4.55 |
| chr11 | 126 | 2.33 | 9 | 3.98 | 17 | 5.52 |
| chr12 | 96 | 1.77 | 5 | 2.21 | 10 | 3.25 |
| chr13 | 65 | 1.20 | 9 | 3.98 | 24 | 7.79 |
| chr14 | 317 | 5.86 | 4 | 1.77 | 0 | 0.00 |
| chr15 | 474 | 8.76 | 11 | 4.87 | 0 | 0.00 |
| chr16 | 153 | 2.83 | 0 | 0.00 | 0 | 0.00 |

**Table S17.** *R. chinensis* genome assembly statistics at the contig level

| Parameter | Value |
|---|---|
| Total length (bp) | 337,528,122 |
| Largest contig (bp) | 25,512,145 |
| Smallest contig (bp) | 15,291 |
| Number of contigs | 105 |

| | |
|---|---|
| Contig N50 (bp) | 17,116,798 |
| Contig N70 (bp) | 8,266,836 |
| Contig N90 (bp) | 3,065,070 |
| Number of N | 1,536 |
| Number of gaps | 73 |
| GC (%) | 35.45 |
| Chromosome-anchored length (bp) | 308,253,885 |
| Anchoring rate (%) | 91.33% |

**Table S18.** Size of each *R. chinensis* chromosome

| **Chromosome** | **Size (bp)** |
|---|---|
| chr1 | 29,700,863 |
| chr2 | 24,035,999 |
| chr3 | 23,159,415 |
| chr4 | 22,918,500 |
| chr5 | 22,334,945 |
| chr6 | 22,319,349 |
| chr7 | 22,239,939 |
| chr8 | 21,022,737 |
| chr9 | 20,351,598 |
| chr10 | 19,547,469 |
| chr11 | 19,114,662 |
| chr12 | 16,915,179 |
| chr13 | 15,842,101 |
| chr14 | 15,411,246 |
| chr15 | 13,339,883 |
| Total | 308,253,885 |

**Table S19.** Repeat content in the *R. chinensis* genome

| **Method** | **Repeat size (bp)** | **% of genome** |
|---|---|---|
| TRF | 16,338,129 | 4.84 |
| RepeatMasker | 5,450,123 | 1.61 |
| RepeatProteinMask | 58,816,776 | 17.41 |
| De novo | 200,482,102 | 59.36 |
| Total (non-redundant) | 209,046,499 | 61.89 |

**Table S20.** Gene prediction of *R. chinensis* compared with related species

| Gene set | Number | Avg gene length (bp) | Avg CDS length (bp) | Avg exon number | Avg exon length (bp) | Avg intron length (bp) |
|---|---|---|---|---|---|---|
| Augustus | 22,585 | 2,742.92 | 1,207.45 | 5.50 | 219.55 | 341.25 |
| GlimmerHMM | 27,595 | 10,235.12 | 998.89 | 4.48 | 222.81 | 2,651.62 |
| Genscan | 14,077 | 11,814.48 | 1,473.76 | 5.09 | 289.55 | 2,528.42 |
| RNAseq | 10,241 | 40,517.42 | 379.22 | 3.46 | 133.64 | 16,280.15 |
| *M. indica* | 34,383 | 3,746.09 | 1,018.38 | 3.84 | 265.37 | 961.29 |
| *P. vera* | 36,110 | 3,365.95 | 979.03 | 3.64 | 268.99 | 904.26 |
| *P. trichocarpa* | 32,189 | 3,733.51 | 976.23 | 3.82 | 255.61 | 978.02 |
| *A. thaliana* | 28,601 | 3,145.04 | 916.70 | 3.91 | 234.25 | 764.89 |
| *R. chinensis* | 23,451 | 3,866.85 | 1,218.93 | 5.14 | 237.91 | 639.68 |

**Table S21.** Gene counts on each *R. chinensis* chromosome

| Chromosome | Gene count | % of total genes |
|---|---|---|
| chr1 | 1,892 | 8.22 |
| chr2 | 1,244 | 5.41 |
| chr3 | 1,056 | 4.59 |
| chr4 | 1,202 | 5.22 |
| chr5 | 1,326 | 5.76 |
| chr6 | 1,608 | 6.99 |
| chr7 | 1,404 | 6.10 |
| chr8 | 1,332 | 5.79 |
| chr9 | 1,418 | 6.16 |
| chr10 | 1,181 | 5.13 |
| chr11 | 1,168 | 5.08 |
| chr12 | 2,105 | 9.15 |
| chr13 | 918 | 3.99 |
| chr14 | 1,083 | 4.71 |
| chr15 | 1,044 | 4.54 |
| Unanchored | 3,024 | 13.14 |

**Table S22.** Functional annotation of the *R. chinensis* genome by database

| Database | Number | Percent (%) |
|---|---|---|
| Total | 23,451 | |
| InterPro | 18,366 | 78.32 |
| GO | 5,884 | 25.09 |
| KEGG | 12,886 | 54.95 |
| KEGG_KO | 8,551 | 36.46 |
| Swissprot | 10,964 | 46.75 |
| TrEMBL | 14,924 | 63.64 |
| NR | 14,242 | 60.73 |
| Annotated (any database) | 19,323 | 82.40 |
| Non-annotated | 4,128 | 17.60 |

**Table S23.** Annotation of non-coding RNA in *R. chinensis*

| Type | Subtype | Copy | Average length (bp) | Total length (bp) | % of genome |
|---|---|---|---|---|---|
| miRNA | | 100 | 122.70 | 12,270 | 0.0036 |
| tRNA | | 568 | 75.35 | 42,801 | 0.0127 |
| rRNA | rRNA | 433 | 563.56 | 244,021 | 0.0722 |
| | 18S | 27 | 1,799.33 | 48,582 | 0.0144 |
| | 28S | 31 | 4,915.68 | 152,386 | 0.0451 |
| | 5S | 375 | 114.81 | 43,053 | 0.0127 |
| snRNA | snRNA | 202 | 123.58 | 24,964 | 0.0074 |
| | CD-box | 103 | 103.71 | 10,682 | 0.0032 |
| | HACA-box | 18 | 128.39 | 2,311 | 0.0007 |
| | splicing | 81 | 147.79 | 11,971 | 0.0035 |
| | scaRNA | 0 | 0 | 0 | 0 |

**Table S24.** BUSCO assessment of *R. chinensis* genome assembly and gene annotation

| Category | Assembly Proteins | Assembly % | Annotation Proteins | Annotation % |
|---|---|---|---|---|
| Complete BUSCOs | 1,572 | 97.39 | 1,482 | 91.82 |
| Complete single-copy BUSCOs | 1,539 | 95.35 | 1,444 | 89.47 |
| Complete duplicated BUSCOs | 33 | 2.04 | 38 | 2.35 |
| Fragmented BUSCOs | 16 | 0.99 | 66 | 4.14 |
| Missing BUSCOs | 26 | 1.61 | 66 | 4.09 |
| Total BUSCO groups | 1,614 | 100 | 1,614 | 100 |

**Table S25.** Number of duplicated genes on each *R. chinensis* chromosome

| Chr | Dispersed | Proximal | Tandem | SDs | % SDs | Total duplicated | % Total |
|---|---|---|---|---|---|---|---|
| chr1 | 785 | 84 | 325 | 356 | 9.97 | 1,550 | 9.11 |
| chr2 | 567 | 127 | 281 | 77 | 2.16 | 1,052 | 6.18 |
| chr3 | 679 | 93 | 207 | 290 | 8.12 | 1,269 | 7.46 |
| chr4 | 486 | 115 | 241 | 174 | 4.87 | 1,016 | 5.97 |
| chr5 | 604 | 86 | 209 | 221 | 6.19 | 1,120 | 6.58 |
| chr6 | 665 | 111 | 255 | 300 | 8.40 | 1,331 | 7.82 |
| chr7 | 618 | 58 | 198 | 276 | 7.73 | 1,150 | 6.76 |
| chr8 | 525 | 101 | 315 | 159 | 4.45 | 1,100 | 6.47 |
| chr9 | 651 | 47 | 173 | 315 | 8.82 | 1,186 | 6.97 |
| chr10 | 491 | 104 | 203 | 219 | 6.13 | 1,017 | 5.98 |
| chr11 | 432 | 89 | 200 | 254 | 7.11 | 975 | 5.73 |
| chr12 | 979 | 61 | 235 | 412 | 11.54 | 1,687 | 9.92 |
| chr13 | 371 | 108 | 172 | 131 | 3.67 | 782 | 4.60 |
| chr14 | 424 | 48 | 176 | 241 | 6.75 | 889 | 5.23 |
| chr15 | 418 | 76 | 248 | 146 | 4.09 | 888 | 5.22 |

**Table S26.** DNA transposable element counts on each *R. chinensis* chromosome

| Chromosome | DNA TEs | % of total TEs |
|---|---|---|
| chr1 | 294,612 | 28.84 |
| chr2 | 72,870 | 7.13 |
| chr3 | 58,873 | 5.76 |
| chr4 | 64,617 | 6.32 |
| chr5 | 53,950 | 5.28 |
| chr6 | 55,379 | 5.42 |
| chr7 | 53,736 | 5.26 |
| chr8 | 60,697 | 5.94 |
| chr9 | 47,901 | 4.69 |
| chr10 | 53,429 | 5.23 |
| chr11 | 54,558 | 5.34 |
| chr12 | 27,616 | 2.70 |
| chr13 | 46,102 | 4.51 |
| chr14 | 43,023 | 4.21 |
| chr15 | 34,286 | 3.36 |

**Table S27.** GC content of each *R. chinensis* chromosome

| Chromosome | GC (%) |
|---|---|
| chr1 | 35.72 |
| chr2 | 35.32 |
| chr3 | 35.43 |
| chr4 | 35.38 |
| chr5 | 35.49 |
| chr6 | 35.29 |
| chr7 | 35.58 |
| chr8 | 35.26 |
| chr9 | 35.91 |
| chr10 | 35.27 |
| chr11 | 35.51 |
| chr12 | 35.32 |
| chr13 | 35.32 |
| chr14 | 35.34 |
| chr15 | 35.55 |

**Table S28.** Number of expanded and contracted genes on each *R. chinensis* chromosome

| Chromosome | Expanded genes | % expanded | Contracted genes | % contracted |
|---|---|---|---|---|
| chr1 | 358 | 9.46 | 388 | 8.97 |
| chr2 | 352 | 9.30 | 188 | 4.35 |
| chr3 | 224 | 5.92 | 336 | 7.77 |
| chr4 | 266 | 7.03 | 239 | 5.52 |
| chr5 | 201 | 5.31 | 258 | 5.96 |
| chr6 | 323 | 8.54 | 344 | 7.95 |
| chr7 | 205 | 5.42 | 298 | 6.89 |
| chr8 | 326 | 8.62 | 236 | 5.46 |
| chr9 | 177 | 4.68 | 331 | 7.65 |
| chr10 | 256 | 6.77 | 271 | 6.26 |
| chr11 | 220 | 5.82 | 248 | 5.73 |
| chr12 | 221 | 5.84 | 554 | 12.81 |
| chr13 | 227 | 6.00 | 193 | 4.46 |
| chr14 | 178 | 4.71 | 262 | 6.06 |
| chr15 | 249 | 6.58 | 180 | 4.16 |

**Table S29.** Proteomics overview across aphid morphs and sample types

Summary of proteomic profiling across five sample groups. Values are derived from the main text (Fig. 3) and are provided to facilitate citation and cross-reference.

| Metric | Value | Notes/Related main figure |
|---|---|---|
| Sample groups | Fx-SG, Fa-SG, Se-SG, Fx-S, Fa-S | See Fig. 3A, G–H |
| Shared proteins across all groups | 1,375 | Fig. 3G |
| Group-specific proteins (Fx-SG) | 61 | Fig. 3G |
| Group-specific proteins (Fa-SG) | 105 | Fig. 3G |
| Group-specific proteins (Fx-S) | 60 | Fig. 3G |
| Group-specific proteins (Fa-S) | 156 | Fig. 3G |

**Table S30.** Chromosomal distribution of hydrolases identified in the *S. chinensis* proteome.

| Chromosome | Hydrolase count | Percentage (%) |
|---|---|---|
| chr1 | 193 | 20.08 |
| chr2 | 90 | 9.37 |
| chr3 | 81 | 8.43 |
| chr4 | 104 | 10.82 |
| chr5 | 73 | 7.6 |
| chr6 | 82 | 8.53 |
| chr7 | 37 | 3.85 |
| chr8 | 41 | 4.27 |
| chr9 | 57 | 5.93 |
| chr10 | 45 | 4.68 |
| chr11 | 43 | 4.47 |
| chr12 | 22 | 2.29 |
| chr13 | 70 | 7.28 |
| chr14 | 6 | 0.62 |
| chr15 | 7 | 0.73 |
| chr16 | 5 | 0.52 |
| Unanchored | 5 | 0.52 |
| **Total** | **961** | **100** |

**Table S31.** Differentially abundant salivary proteins and M20 metallopeptidase candidates

| Comparison / item | Result | Notes/Related main figure |
|---|---|---|
| Fa-S vs Fx-S (upregulated proteins) | 314 | Fig. 3L |
| Fa-S vs Fx-S (downregulated proteins) | 186 | Fig. 3L |
| M20 candidates highlighted | FAAH2; CNDP2-1; ACY1; CNDP2-2; ADL; GatA | Fig. 3L–M |
| RT-qPCR validation (Fa-SG vs Se-SG) | GatA, FAAH2, ACY1 significantly higher ($P < 0.001$ or $P < 0.0001$) | Fig. 3N |

**Table S32.** Metabolomics overview across aphid morphs and sample types

| Metric | Value | Notes/Related main figure |
|---|---|---|
| Identified metabolites (total) | 1,475 | Fig. 4A |
| Metabolites assigned to a superclass | 1,313 | Fig. 4A |
| Top superclass: organic acids and derivatives | 327 (24.90%) | Fig. 4A |
| Second superclass: lipids and lipid-like molecules | 305 (23.23%) | Fig. 4A |
| Third superclass: organoheterocyclic compounds | 196 (14.93%) | Fig. 4A |
| Differential metabolites (\|log2FC\| > 1, $P < 0.05$) | 288 (in 5/6 comparisons); 131 (in all 6) | Fig. 4B |
| Fa-S vs Fx-S differential metabolites | 928 up; 79 down | Fig. 4E |
| Hormone-related enrichment | IAA and ABA enriched in gall-inducing saliva (Fa-S) | Fig. 4D–F |

**Table S33.** RNAi knockdown efficiency and phenotypic outcomes

| Group | Relative ACY1 expression | Galling rate (%) | Mortality (%) | Notes/Related main figure |
|---|---|---|---|---|
| Blank control | 1.000 ± 0.000 | 89.83 ± 2.40 | 10.17 ± 2.40 | Fig. 5B–D |
| GFP-dsRNA | 1.035 ± 0.083 | 90.50 ± 1.64 | 9.50 ± 1.64 | Fig. 5B–D |
| ds*ACY1* | 0.238 ± 0.093 | 52.33 ± 6.02 | 47.67 ± 6.73 | Tukey-adjusted $P < 0.0001$ vs both controls |

**Table S34.** Auxin metabolite profiling at the galling/settlement zone under *ACY1* RNAi

| Metabolite | Control (approx.) | ds*ACY1* (mean ± s.d.) | Direction | Notes |
|---|---|---|---|---|
| Free IAA (pmol $g^{-1}$ FW) | ~509 (507.5–510.0) | 43.83 ± 2.43 | decrease | $P < 0.0001$ |
| IAA-Leu | ~0.606 | 0.787 ± 0.045 | increase | $P < 0.0001$ |
| IAA-Phe | ~0.349 | 0.463 ± 0.038 | increase | $P < 0.0001$ |
| IAA-Tyr | ~0.237 | 0.338 ± 0.044 | increase | $P = 0.0001$ |
| IAA-Ala | ~0.303 | 0.377 ± 0.059 | increase | $P < 0.01$ |
| IAA-Asp | ~15.46 | 17.33 ± 0.99 | increase | $P < 0.02$ |
| IAA-Glu | ~10.94 | 12.50 ± 1.16 | increase | $P < 0.02$ |
| IAA-Glc | ~41.29 | 43.39 ± 0.58 | increase | $P < 0.02$ |

**Table S35.** Correlation between *ACY1* expression, galling phenotypes and auxin metabolites

| Variable pair | Pearson r | P value | Direction | Related figure |
|---|---|---|---|---|
| *ACY1* expression vs galling rate | 0.95 | $1.2 \times 10^{-9}$ | positive | Fig. 5G |
| *ACY1* expression vs mortality | −0.95 | < 0.001 (FDR-significant) | negative | Fig. 5H |
| *ACY1* expression vs Free IAA | 0.867 | $3.28 \times 10^{-6}$ | positive | Fig. 5I |
| *ACY1* expression vs IAA-Phe | −0.906 | FDR-significant | negative | Fig. 5I |

| | | | | |
|---|---|---|---|---|
| *ACY1* expression vs IAA-Leu | −0.891 | FDR-significant | negative | Fig. 5I |
| *ACY1* expression vs IAA-Tyr | −0.878 | FDR-significant | negative | Fig. 5I |

**Table S36.** Cell diameter differences in the settlement zone under *ACY1* RNAi

| Tissue type | Control (µm) | ds*ACY1* (µm) | P value | Related figure |
|---|---|---|---|---|
| Epidermal cells | 13.21 | 8.74 | 0.022 | Fig. 5L |
| Palisade cells | 20.02 | 11.07 | $3.0 \times 10^{-6}$ | Fig. 5L |
| Spongy mesophyll cells | 11.75 | 8.24 | 0.002 | Fig. 5L |

**Table S37.** Key outcomes of ACY1 cell-free enzyme treatments. Selected quantitative outcomes highlighting dose-dependent liberation of free IAA and IAA-AA.

| Metric | Control | ACY1 (key dose) | Effect / notes |
|---|---|---|---|
| Free IAA (pmol $g^{-1}$ FW) | 481.37 ± 21.49 (HI control) | 594.01 ± 23.02 (4 µM) | +23.4%; Holm-adjusted $P < 0.001$ |
| Free-to-conjugated IAA ratio | ~7.4 (controls) | 11.33 ± 0.90 (4 µM) | Holm-adjusted $P < 0.001$ |
| Free IAA proportion of total IAA | ~88% (controls) | 91.86 ± 0.62% (4 µM) | $P < 0.001$ |
| Conjugate reduction at 4 µM | — | IAA-Leu −60.2%; IAA-Ala −56.4%; IAA-Tyr −52.6%; IAA-Asp −40.5%; IAA-Glu −40.3% | Dose-dependent decreases |

### Supplementary figures

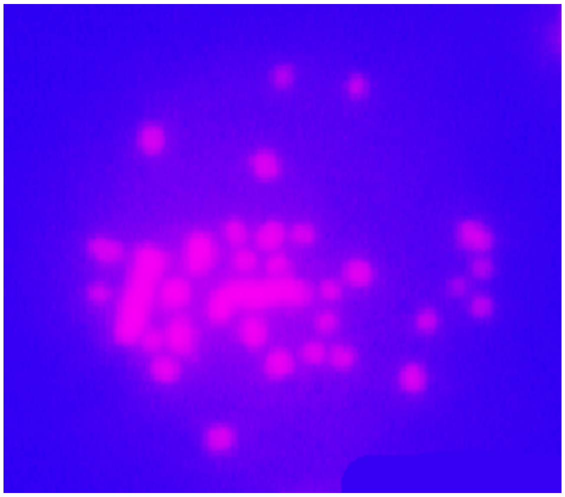

**Fig. S1. Giemsa staining of *S. chinensis* chromosomes.** Giemsa-stained karyotype spread of *S. chinensis* showing chromosome morphology. This cytogenetic preparation complements the DAPI-FISH analysis (Fig. 1b) and confirms the 2n = 32 chromosome count.

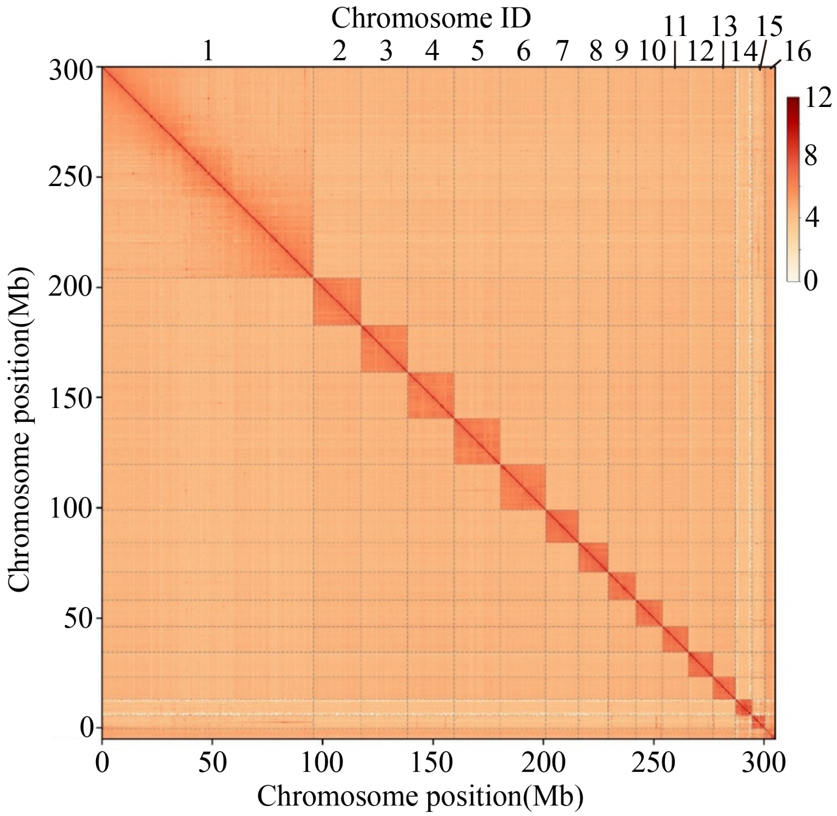


**Fig. S2. Hi-C interaction heatmap of the *S. chinensis* genome at the chromosome level.** Genome-wide Hi-C chromatin interaction frequency matrix of Schlechtendalia chinensis. The x- and y-axes represent chromosomal positions (Mb), with 16 chromosomes labeled. Color intensity indicates the log-transformed interaction frequency, with deeper red denoting higher contact probability. Prominent diagonal squares correspond to intra-chromosomal interactions, confirming successful chromosome-level scaffolding. Chr1 spans the largest genomic interval (~100 Mb), harboring >30% of all anchored genes.

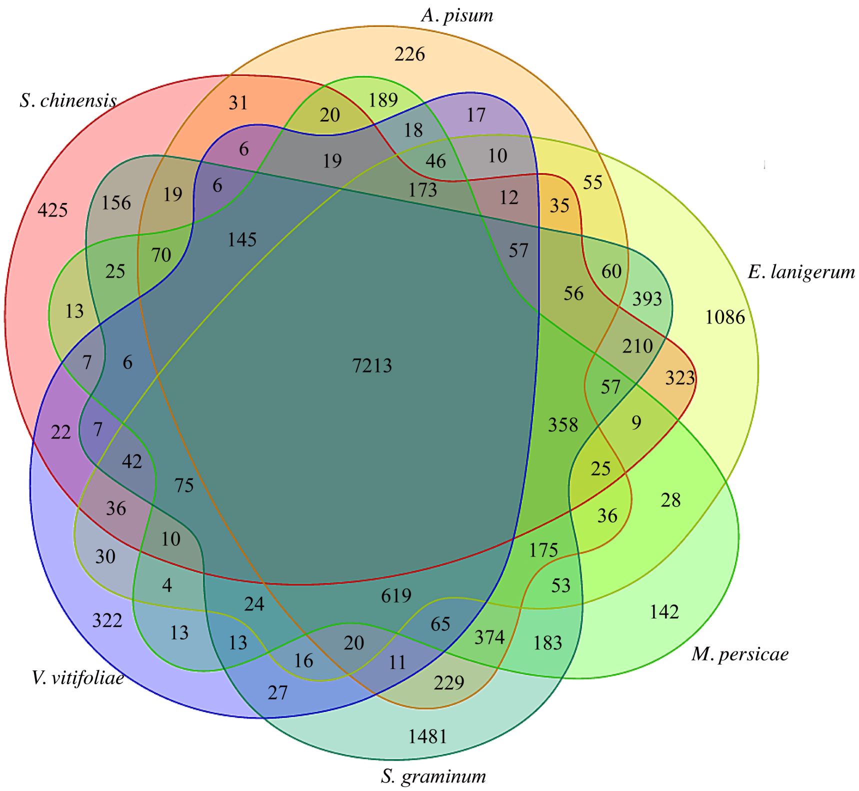


**Fig. S3. Venn diagram of gene family overlap between galling and non-galling aphids.** Six-set Venn diagram comparing orthologous gene families among galling aphids (*S. chinensis*, *E. lanigerum, V. vitifoliae*) and non-galling aphids (*A. pisum, M. persicae, S. graminum*). A core set of 7,213 families is shared. *S. chinensis* retains 425 lineage-specific families; the three galling species collectively share 145 families exclusive to the galling clade.

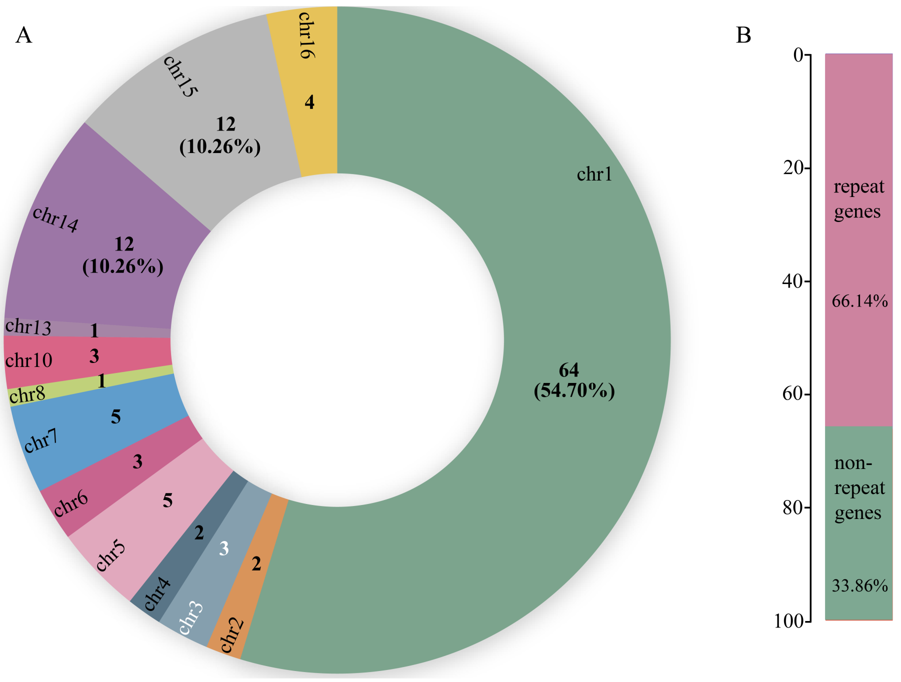


**Fig. S4. Overview of galling aphid-specific gene families: chromosomal distribution and repeat composition.** (A) Donut chart showing the chromosomal distribution of galling aphid-specific gene families in *S. chinensis*. Chr1 harbors the largest share (64 genes, 54.70%). (B) Repeat-associated genes (66.14%) versus non-repeat genes (33.86%), suggesting transposon-mediated duplication drove their expansion.

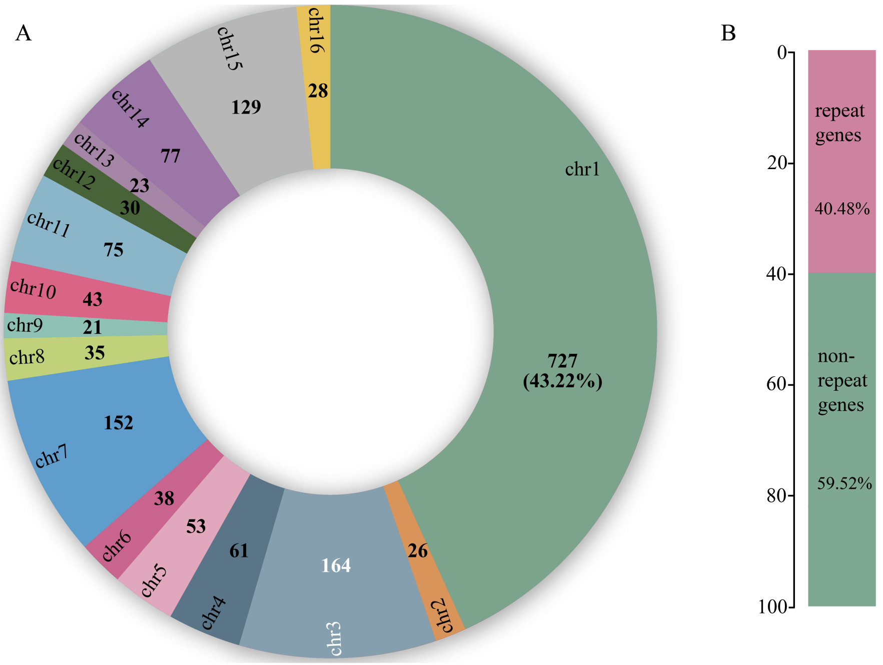


**Fig. S5. Overview of *S. chinensis* species-specific genes: chromosomal distribution and repeat composition.** (A) Donut chart of 727 species-specific genes; chr1 accounts for 43.22%, followed by chr3 (164) and chr7 (152). (B) Repeat genes (40.48%) versus non-repeat genes (59.52%). Compared with galling-specific genes (Fig. S4), lower repeat content suggests mixed duplication and de novo origination mechanisms.

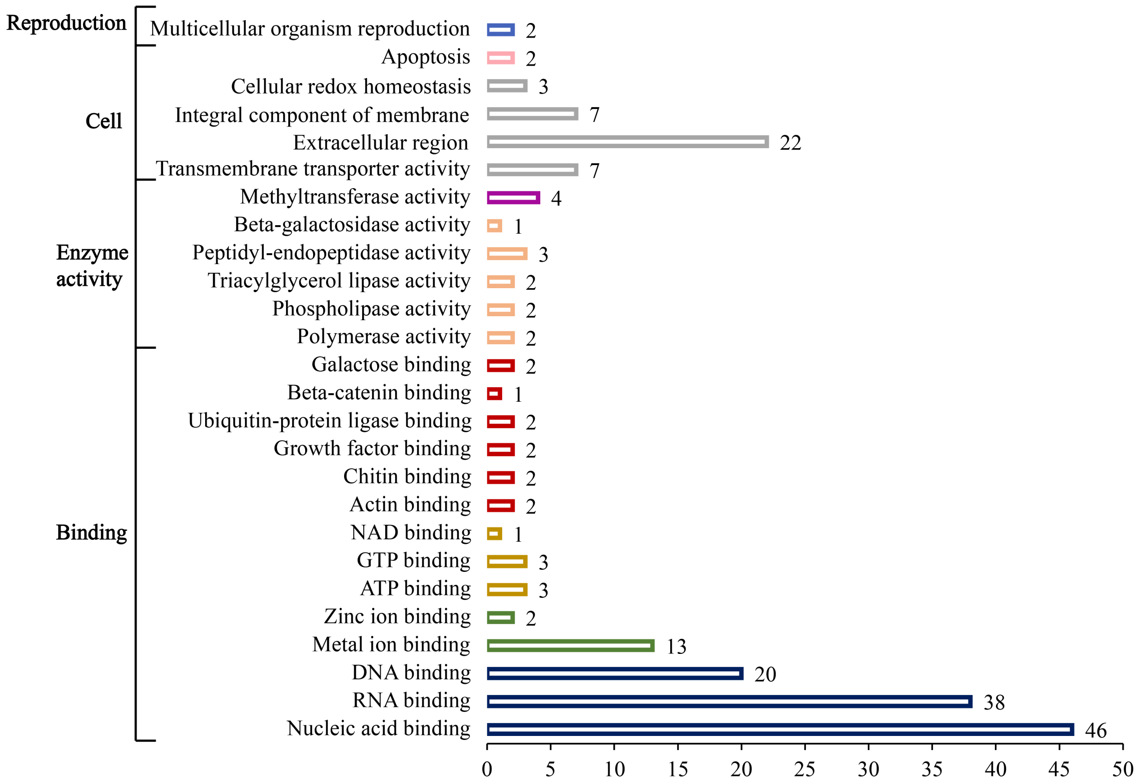


**Fig. S6. GO classification of *S. chinensis* species-specific genes.** Horizontal bar chart of GO functional classification of 727 species-specific genes, organized by biological process, cellular component, and molecular function. Nucleic acid binding (46 genes), RNA binding (38), and DNA binding (20) are the most represented molecular functions; extracellular region (22 genes) is prominent among cellular components.

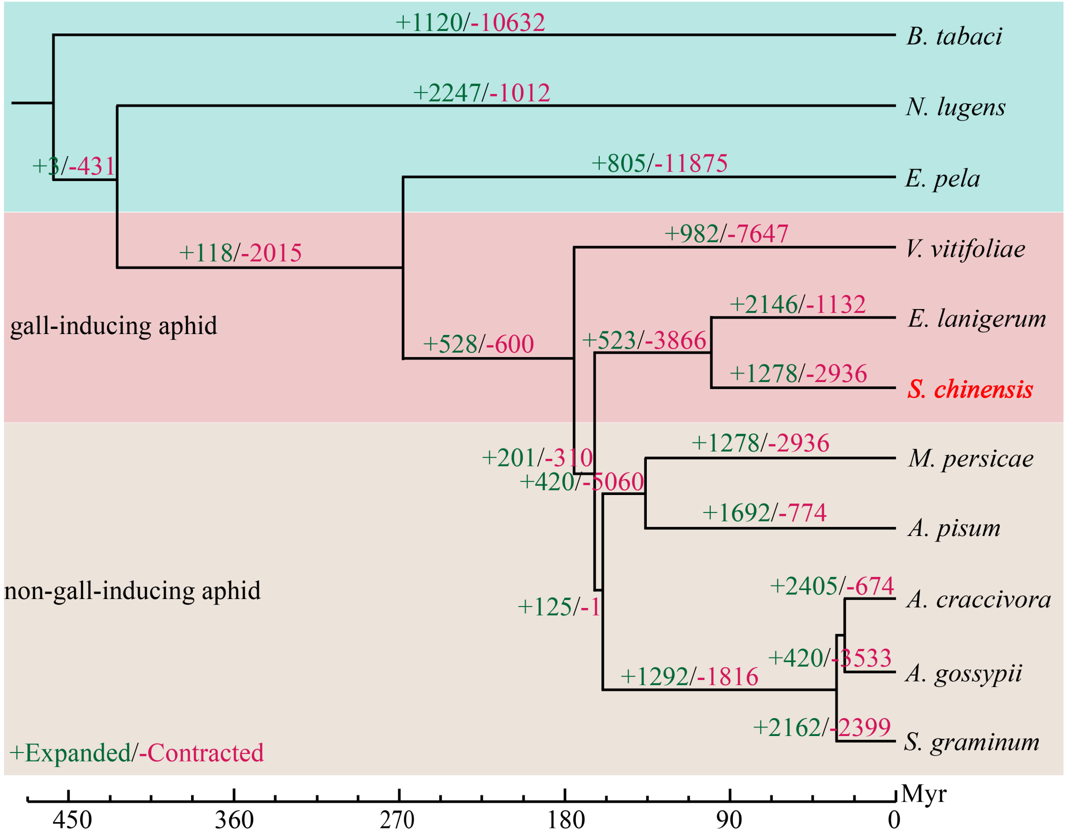


**Fig. S7. Numbers of expanded and contracted gene families across the Aphididae phylogeny.** Time-calibrated phylogenetic tree of Aphididae showing expanded (+, green) and contracted (−, red) gene family numbers per branch. Galling aphids (pink) versus non-galling (green) are distinguished. *S. chinensis*: +1,278/−2,936 families. Time scale in million years.

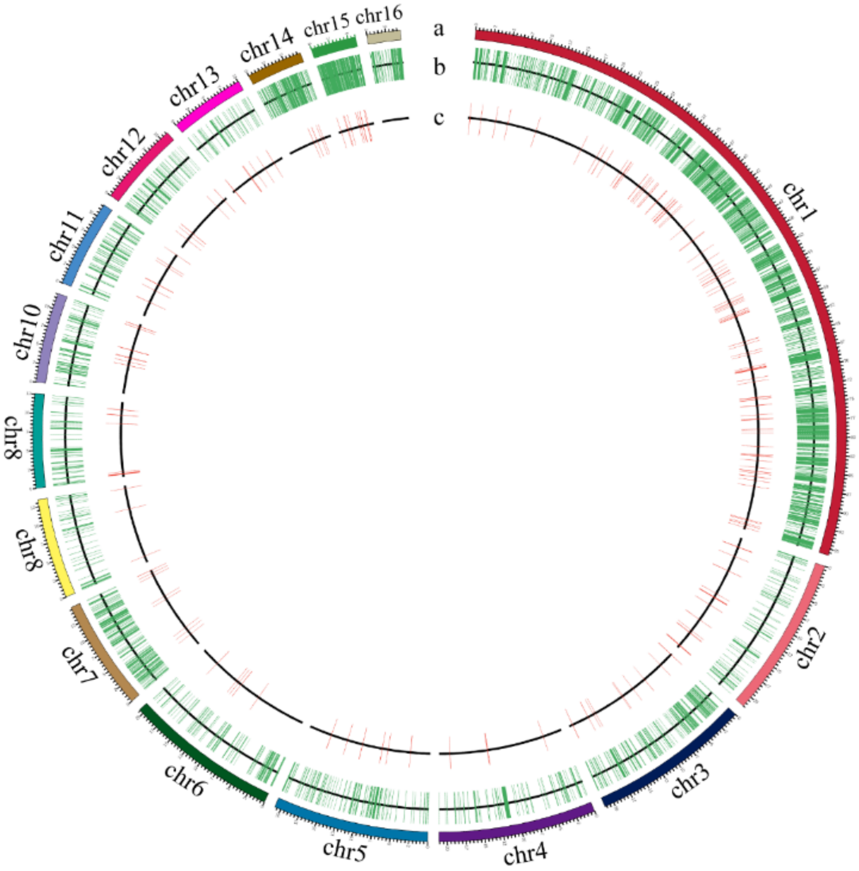


**Fig. S8. Chromosomal distribution of expanded and contracted genes in *S. chinensis*.** Circos plot of expanded/contracted gene families across 16 chromosomes. Tracks: (a) chromosome ideogram; (b) expanded (green) and contracted (red) gene density; (c) reference ring. Chr1 carries the highest expanded gene density.

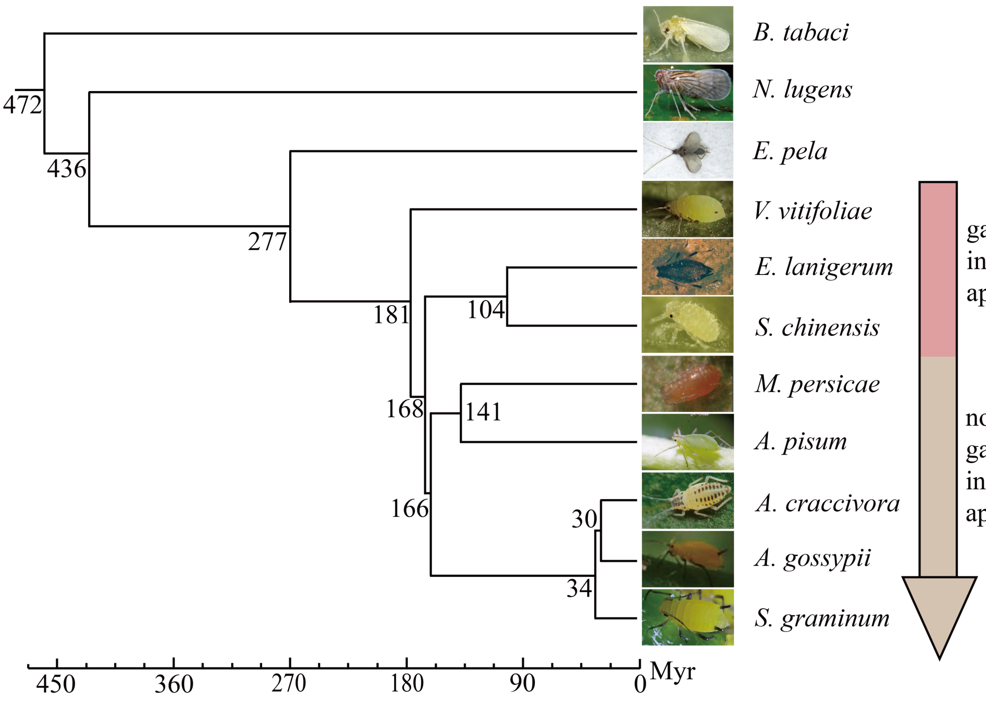


**Fig. S9. Genome-based phylogenetic tree of Aphididae species.** Maximum-likelihood phylogeny of Aphididae constructed from single-copy orthologous genes. Supports monophyly of galling aphids with *S. chinensis* sister to *E. lanigerum*. This tree serves as the framework for gene family expansion/contraction and positive selection analyses.

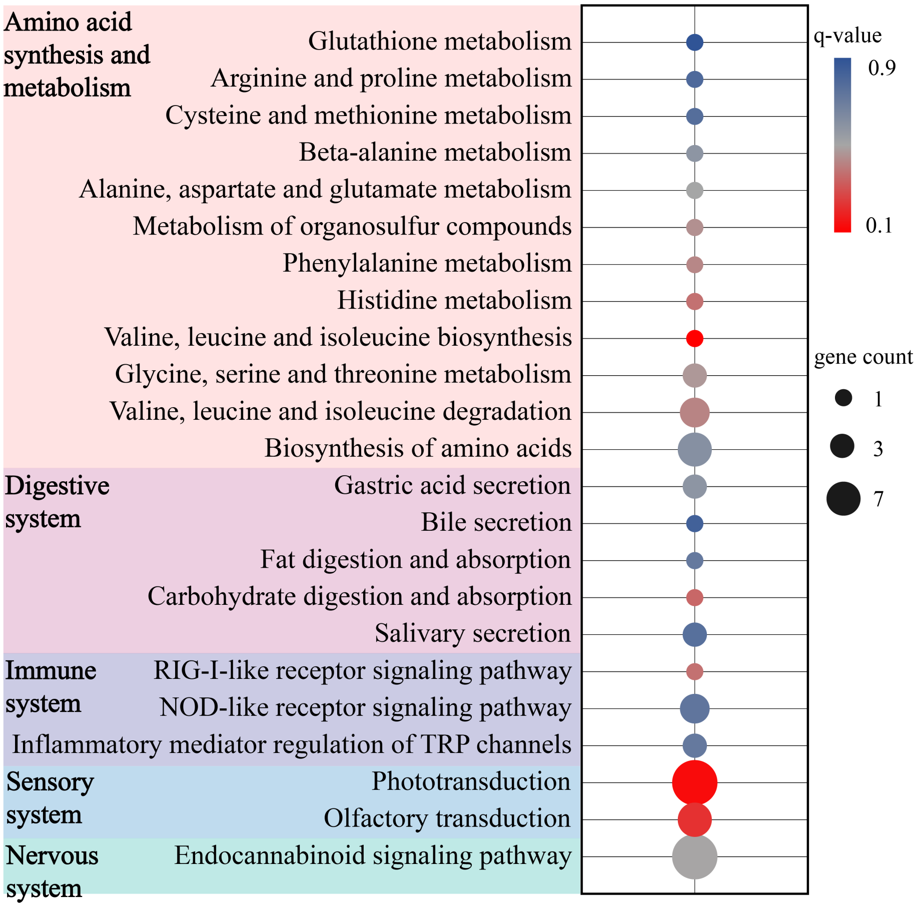


**Fig. S10. KEGG enrichment of positively selected genes in *S. chinensis*.** Bubble plot of KEGG pathway enrichment for positively selected genes (FDR < 0.05) grouped by functional category. Phototransduction and olfactory transduction are among the most enriched pathways, consistent with sensory adaptation in the gall-forming lineage. Bubble size represents gene count; color indicates P value.

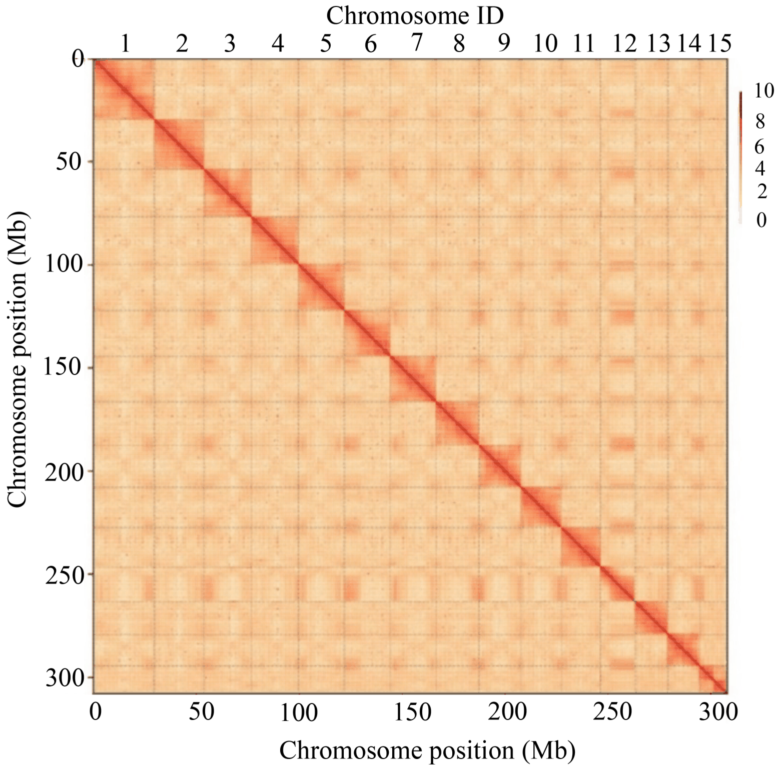


**Fig. S11. Hi-C interaction heatmap of the *R. chinensis* genome at the chromosome level.** Genome-wide Hi-C chromatin interaction frequency matrix of *R. chinensis* with 15 chromosomes labeled. Strong diagonal signals confirm successful anchoring of scaffolds to 15 pseudo-chromosomes. Compared with *S. chinensis* (Fig. S2), chromosomes are more uniform in size, consistent with the balanced gene distribution (Fig. 2b).

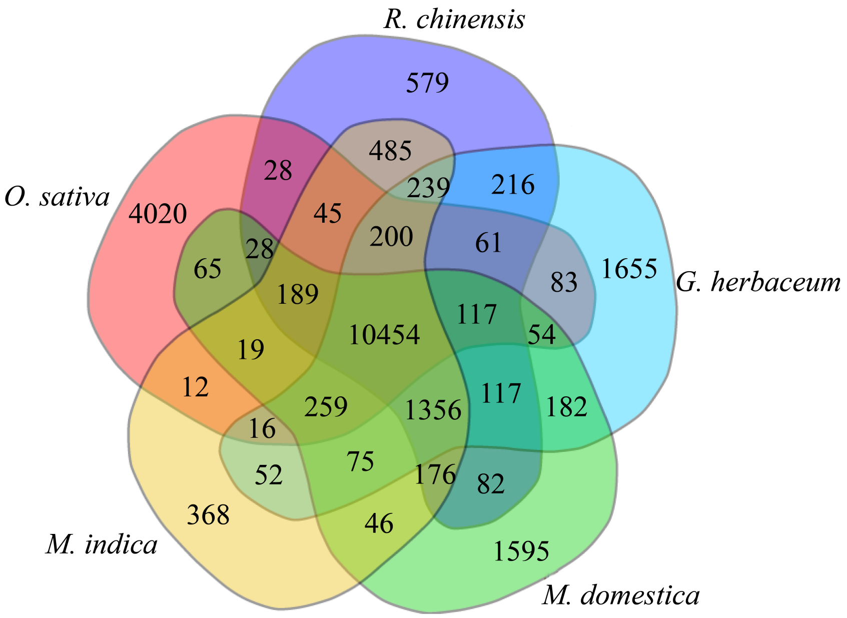


**Fig. S12. Venn diagram of gene family overlap among selected plant species including *R. chinensis*.** Five-set Venn diagram comparing gene families among *R. chinensis, O. sativa, G. herbaceum, M. indica,* and *M. domestica*. A core of 10,454 families is shared. *R. chinensis* retains 579 species-specific families, the highest among the five species.

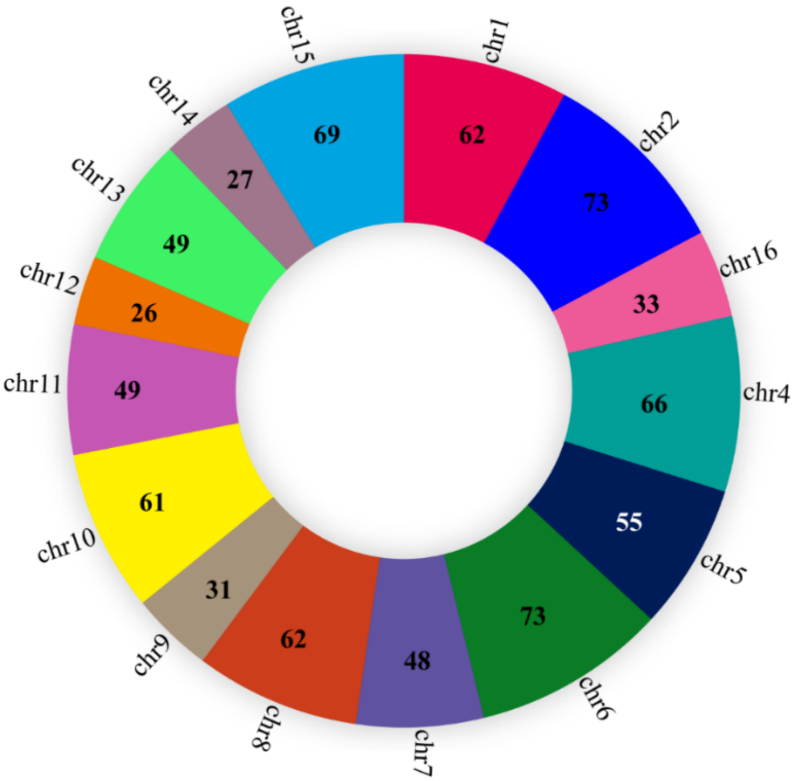


**Fig. S13. Chromosomal distribution of *R. chinensis* species-specific genes.** Donut chart showing species-specific gene distribution across 15 chromosomes. Chr6 (73 genes), chr4 (66), chr1 and chr9 (62 each) are most gene-rich, indicating a relatively even distribution compared with the chr1-dominated pattern in *S. chinensis*.

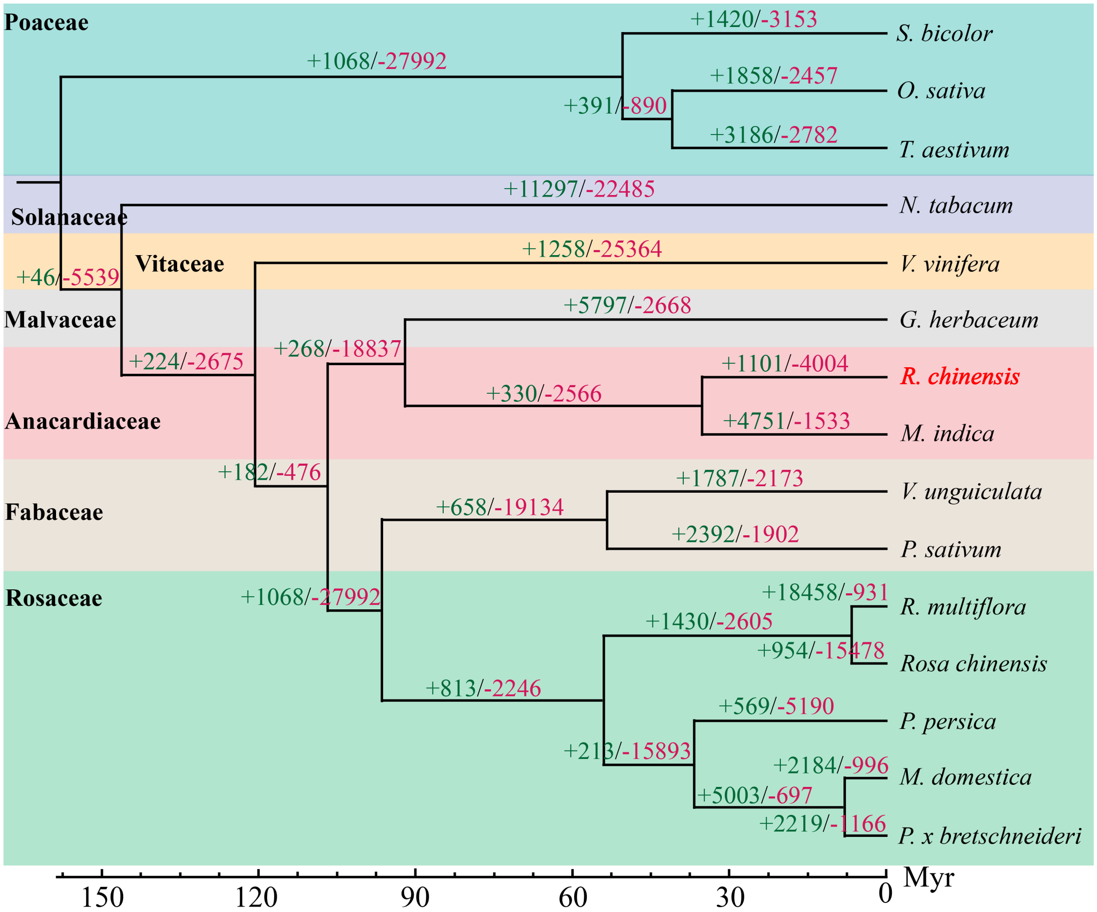


**Fig. S14. Numbers of expanded and contracted gene families across the host plant phylogeny.** Time-calibrated phylogenetic tree of angiosperms showing expanded/contracted gene families. *R. chinensis*: +1,101/−4,004 families. Species are color-coded by family: Poaceae, Solanaceae, Vitaceae, Malvaceae, Anacardiaceae, Fabaceae, Rosaceae.

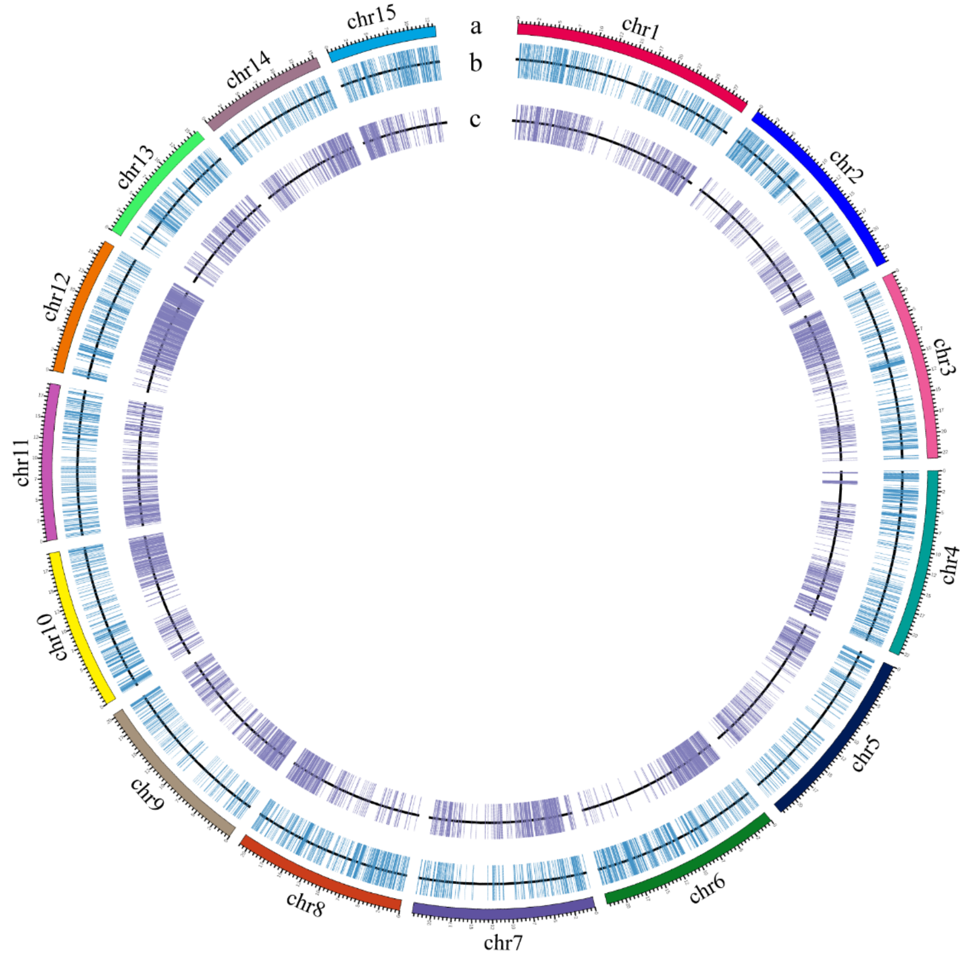


**Fig. S15. Chromosomal distribution of expanded and contracted genes in *R. chinensis*.** Circos plot of expanded/contracted genes across 15 chromosomes. Unlike *S. chinensis*, expanded genes are distributed more evenly, with chr1 and chr12 showing moderate enrichment.

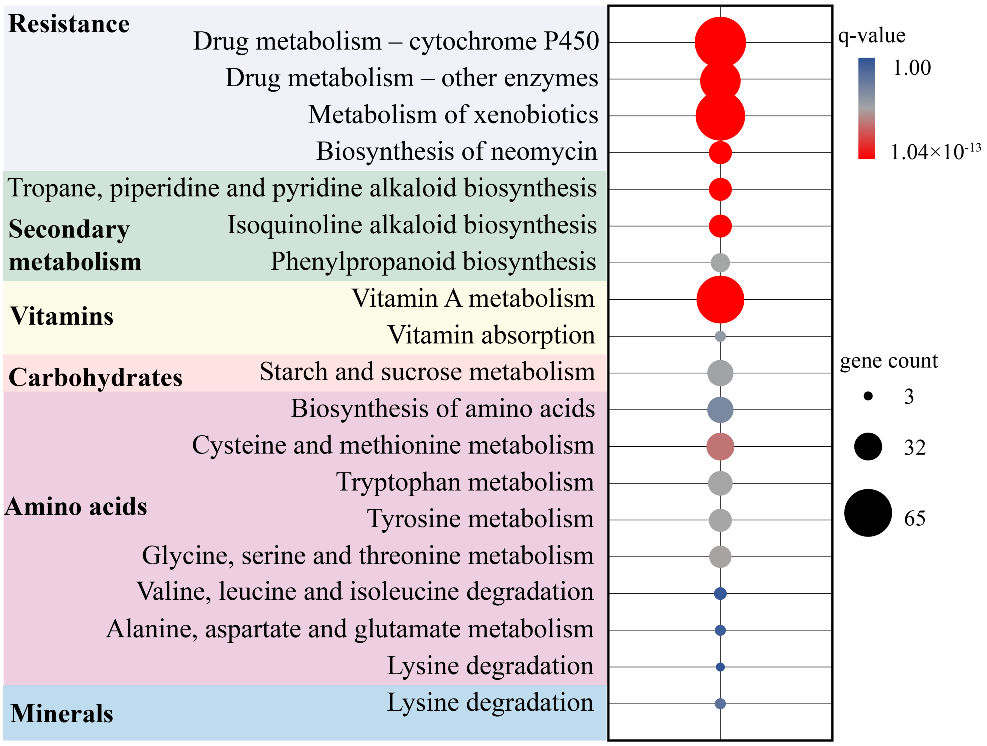


**Fig. S16. KEGG enrichment of expanded gene families in *R. chinensis*.** KEGG pathway enrichment for expanded gene families. Key pathways include pentose and glucuronate interconversions, drug metabolism–cytochrome P450, xenobiotic metabolism by cytochrome P450, and steroid hormone biosynthesis, consistent with large-scale expansion of secondary metabolism and detoxification genes (Fig. 2d, e). Bubble size represents gene count; color indicates Q value.

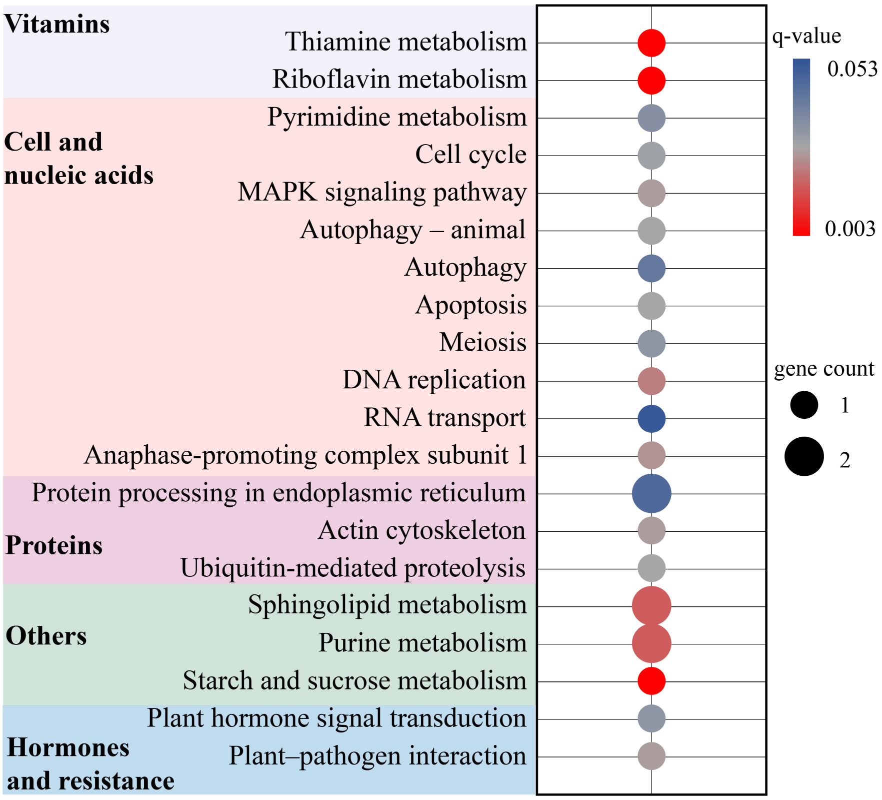


**Fig. S17. KEGG enrichment of positively selected genes in *R. chinensis*.** KEGG pathway enrichment for positively selected genes. Enriched pathways ($Q < 0.05$) include plant hormone signal transduction, plant–pathogen interaction, and MAPK signaling–plant, suggesting adaptive evolutionary pressure on hormone signaling and defense pathways in the host plant.

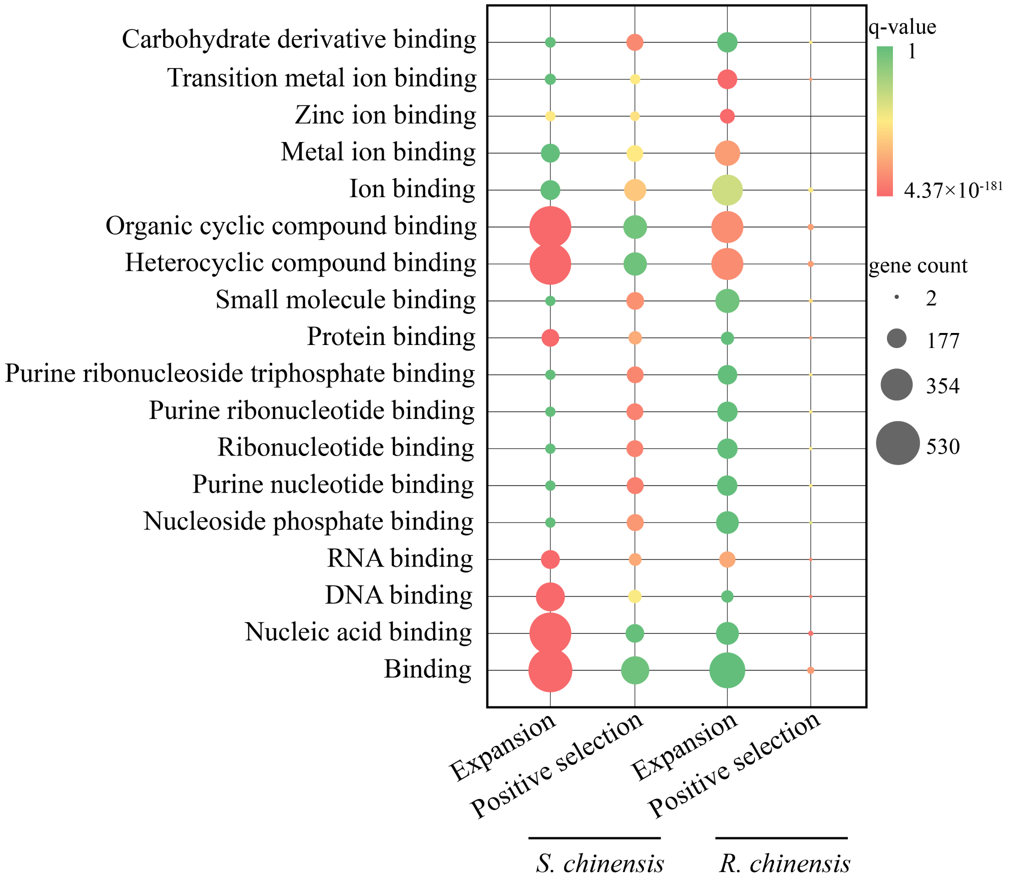


**Fig. S18. GO enrichment comparison of expanded and positively selected genes between *S. chinensis* and *R. chinensis*.** Comparative GO enrichment of expanded and positively selected genes between the aphid and its host plant, highlighting convergent and divergent functional enrichment and shared selective pressures at the insect–plant interface.

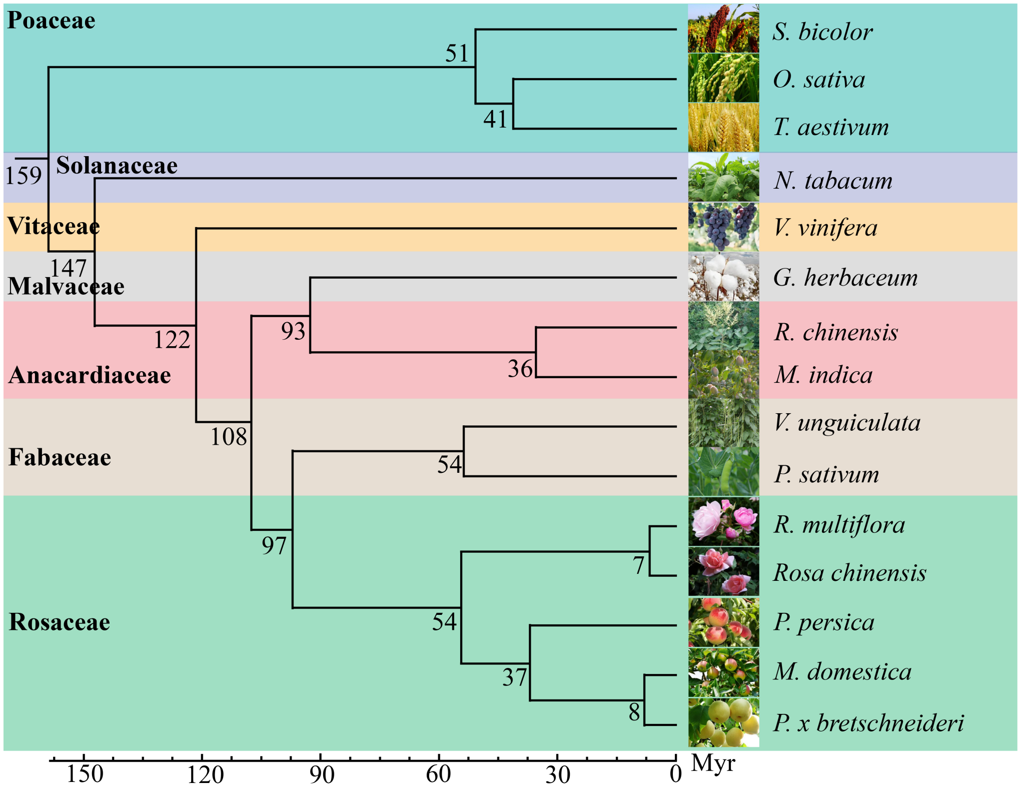


**Fig. S19. Genome-based phylogenetic tree of host plant species.** Maximum-likelihood phylogeny of angiosperms from single-copy orthologous genes. *R. chinensis* (Anacardiaceae) is placed within Sapindales. This tree serves as the framework for host plant comparative genomics analyses.

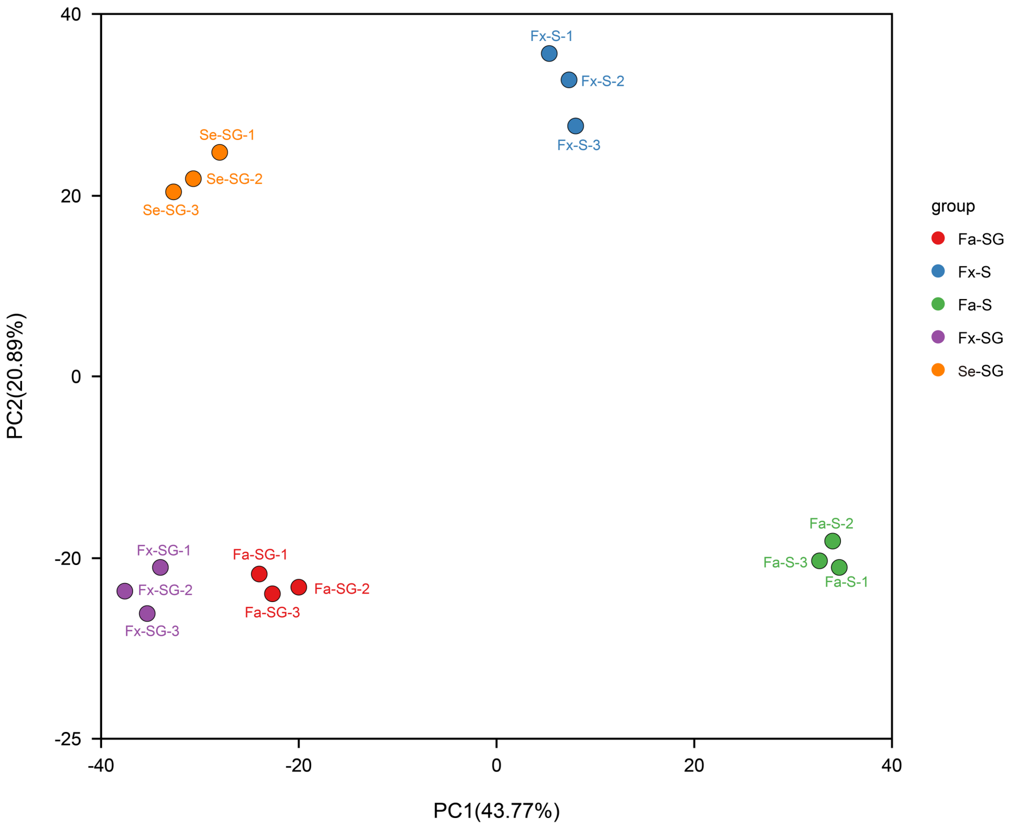


**Fig. S20. PCA of proteomic sample quality control.** Principal component analysis (PCA) score plot of all proteomic samples, used for quality assessment. Biological replicates within each group cluster tightly, indicating high reproducibility. The clear separation among sample groups confirms distinct proteomic profiles across different aphid morphs and tissue types.

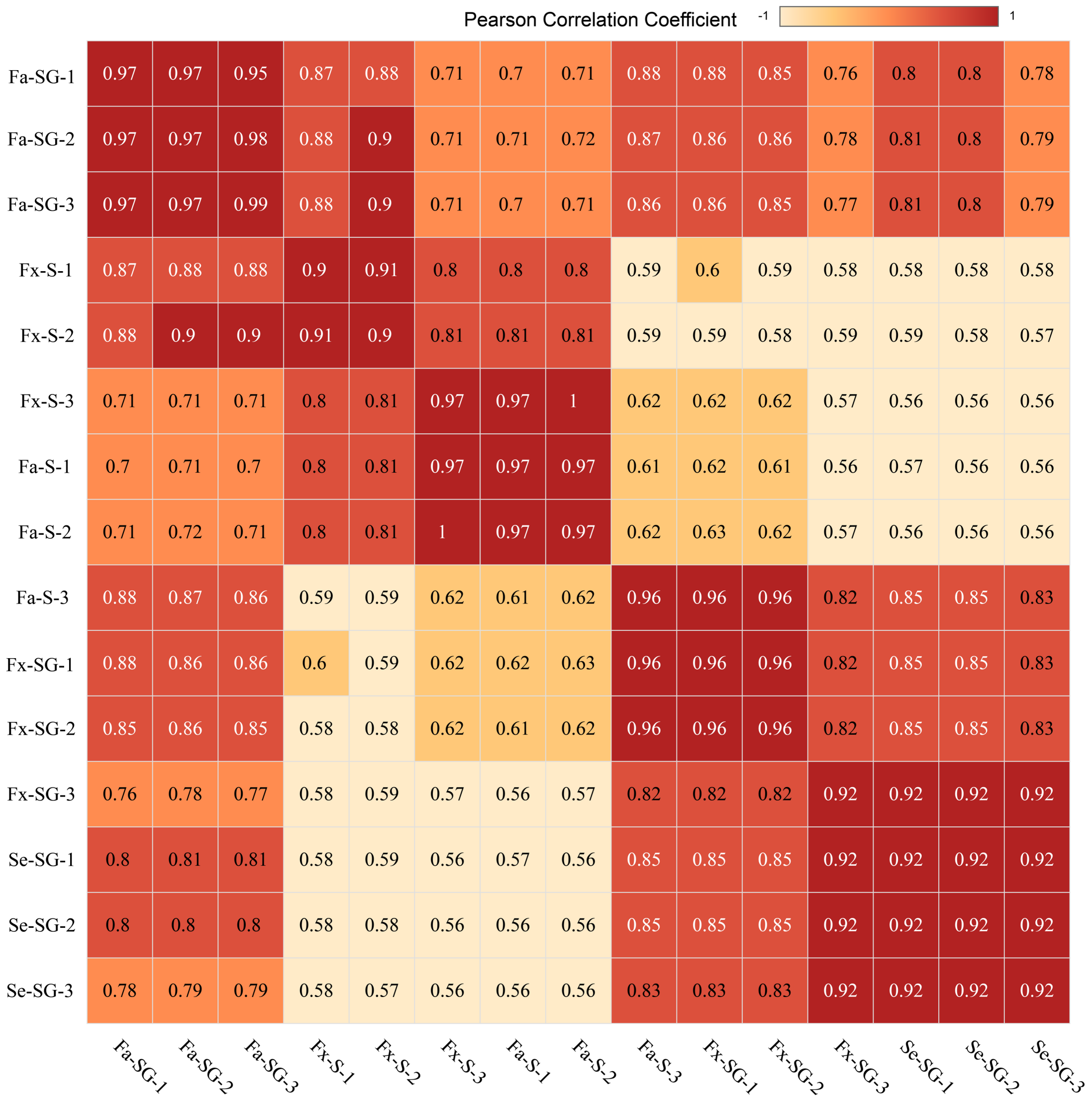


**Fig. S21. Pearson correlation coefficient (PCC) matrix of proteomic samples.** Pairwise Pearson correlation heatmap of protein abundance profiles across all 15 proteomic samples (3 replicates × 5 groups). High intra-group correlations (r > 0.9) confirm biological reproducibility. Inter-group correlations reflect functional similarity, with salivary gland samples clustering separately from saliva samples.

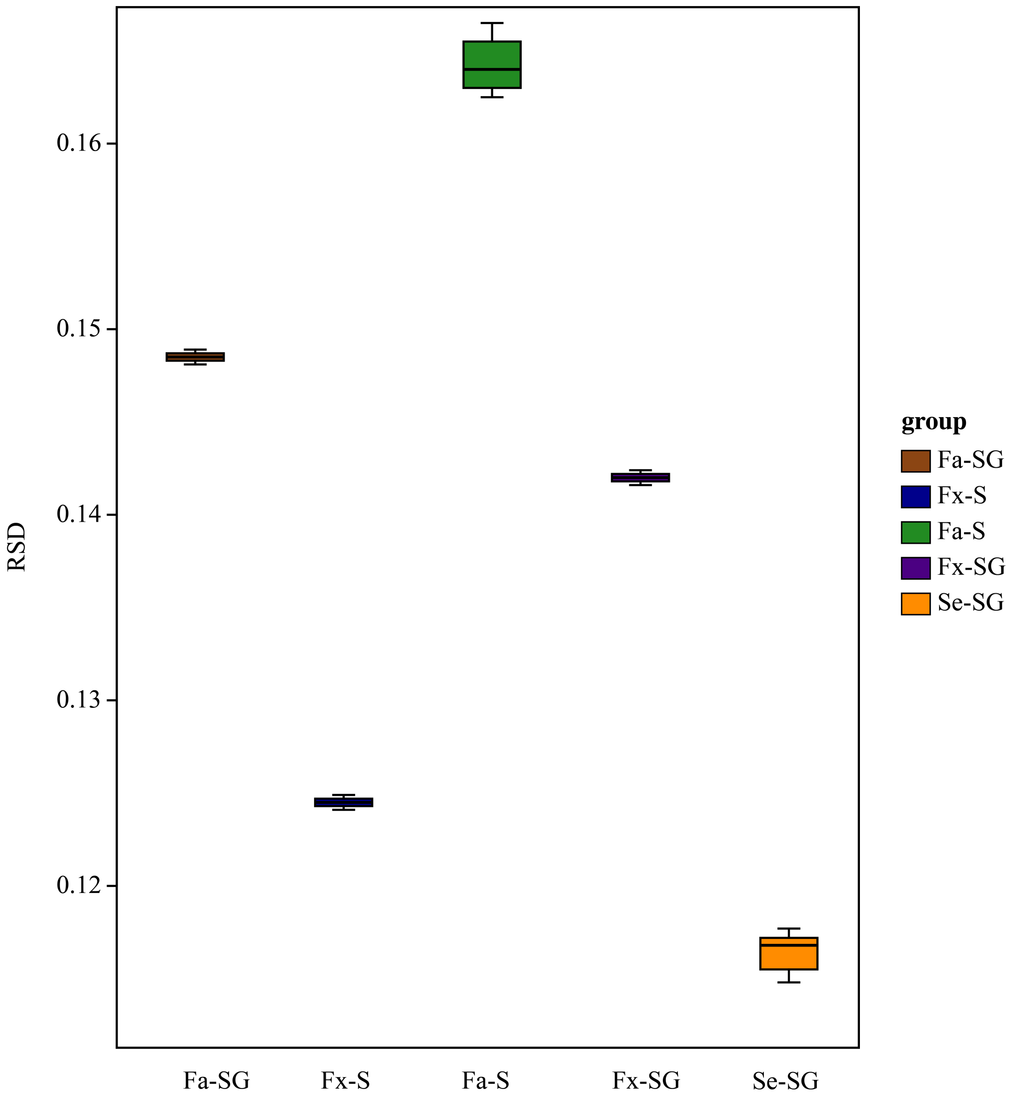


**Fig. S22. Relative standard deviation (RSD) distribution of proteomic quantification.** Distribution of relative standard deviation (RSD) values across quantified proteins in each sample group. The majority of proteins show RSD < 30%, confirming the reliability and precision of the label-free quantification approach.

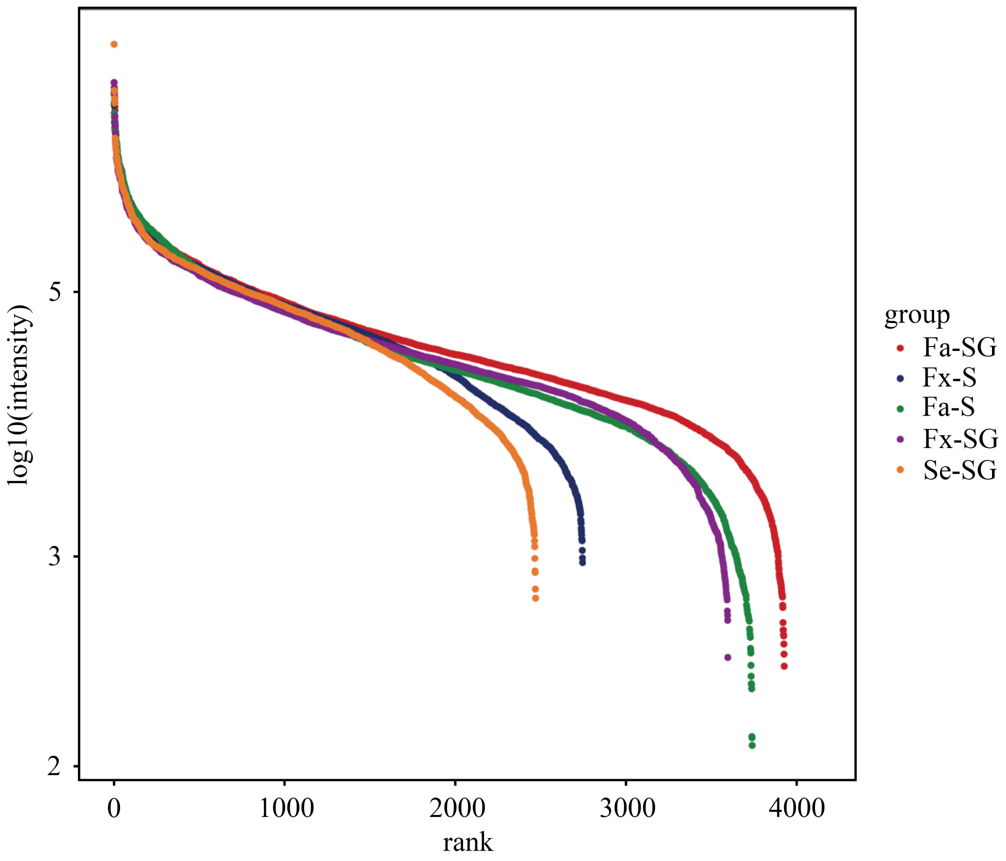


**Fig. S23. Overall protein intensity distributions across proteomic samples after normalization.** Rank-intensity plot of $log_{10}$ protein intensities across five groups after normalization. Overlapping curves confirm comparable dynamic ranges. Se-SG shows slightly fewer quantified proteins, consistent with the smaller proteome of the non-feeding sexual morph.

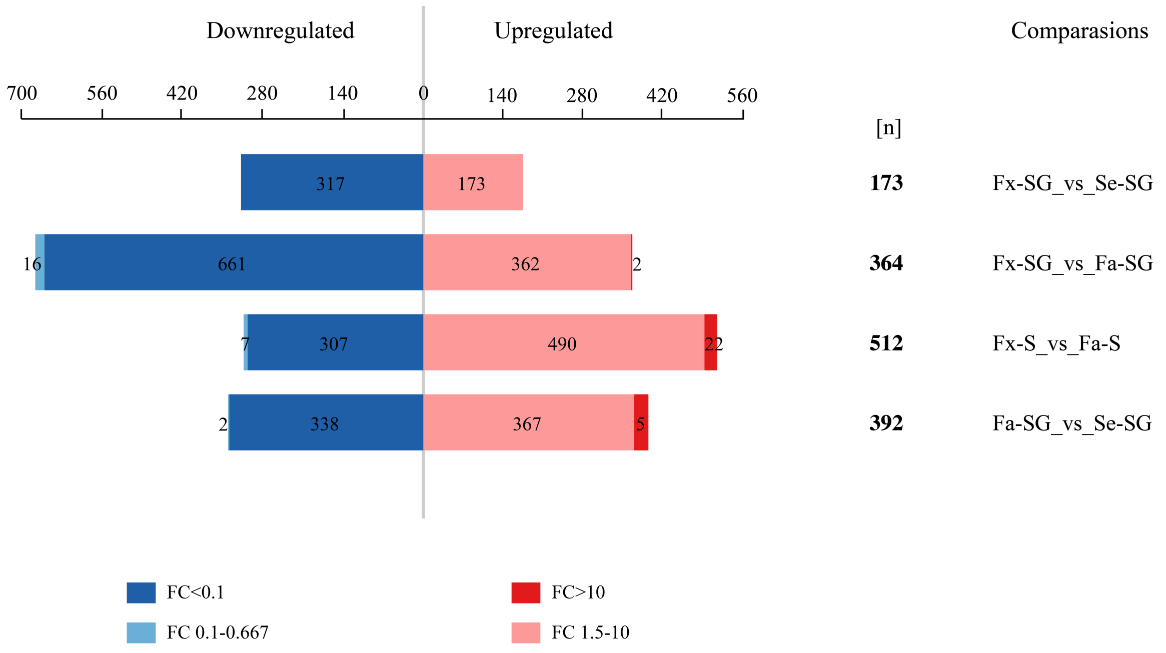


**Fig. S24. Summary of differentially abundant proteins across pairwise comparisons.** Bar chart summarizing upregulated (red) and downregulated (blue) protein numbers in four pairwise comparisons: Fx-SG vs Se-SG, Fx-SG vs Fa-SG, Fx-S vs Fa-S, Fa-SG vs Se-SG. Fold-change thresholds are color-coded; total number of differentially expressed proteins [n] is indicated per comparison.

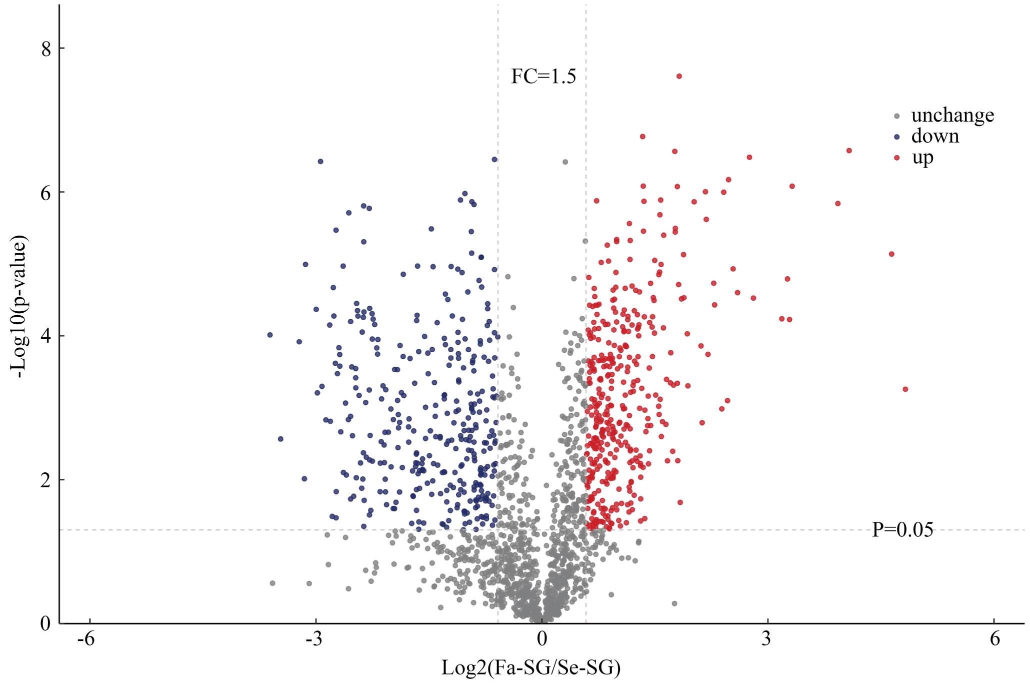


**Fig. S25. Volcano plot of differentially abundant proteins: Fa-SG vs Se-SG.** Volcano plot showing –$\log_{10}$(P-value) versus $\log_2$(fold change) for the comparison between fundatrigenia salivary gland (Fa-SG) and sexuparae salivary gland (Se-SG). Red dots: significantly upregulated in Fa-SG; blue dots: significantly downregulated. Thresholds: $|\log_2 FC| > 1$, $P < 0.05$.

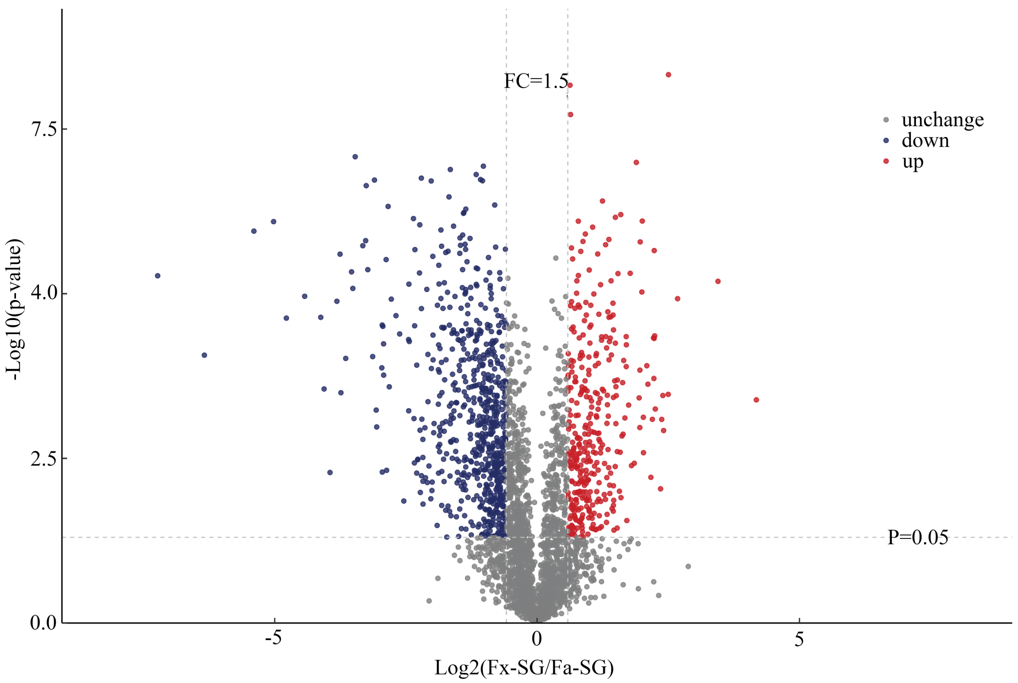


**Fig. S26. Volcano plot of differentially abundant proteins: Fx-SG vs Fa-SG.** Volcano plot for the comparison between fundatrix salivary gland (Fx-SG) and fundatrigenia salivary gland (Fa-SG). Differentially abundant proteins are colored by significance and fold-change direction.

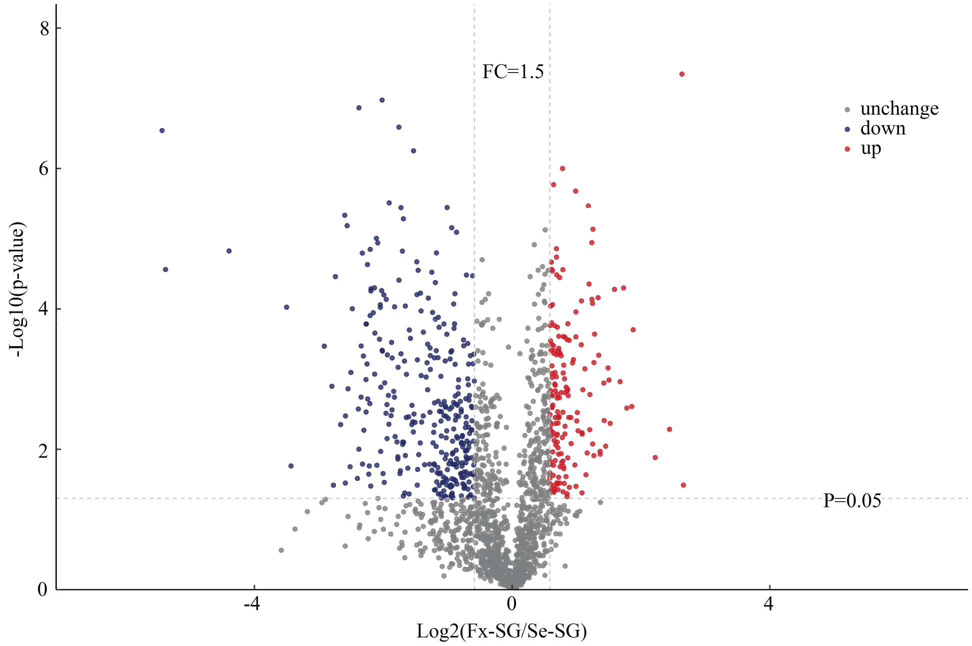


**Fig. S27. Volcano plot of differentially abundant proteins: Fx-SG vs Se-SG.** Volcano plot for the comparison between fundatrix salivary gland (Fx-SG) and sexuparae salivary gland (Se-SG). This comparison reveals proteins associated with salivary gland activity in feeding versus non-feeding morphs.

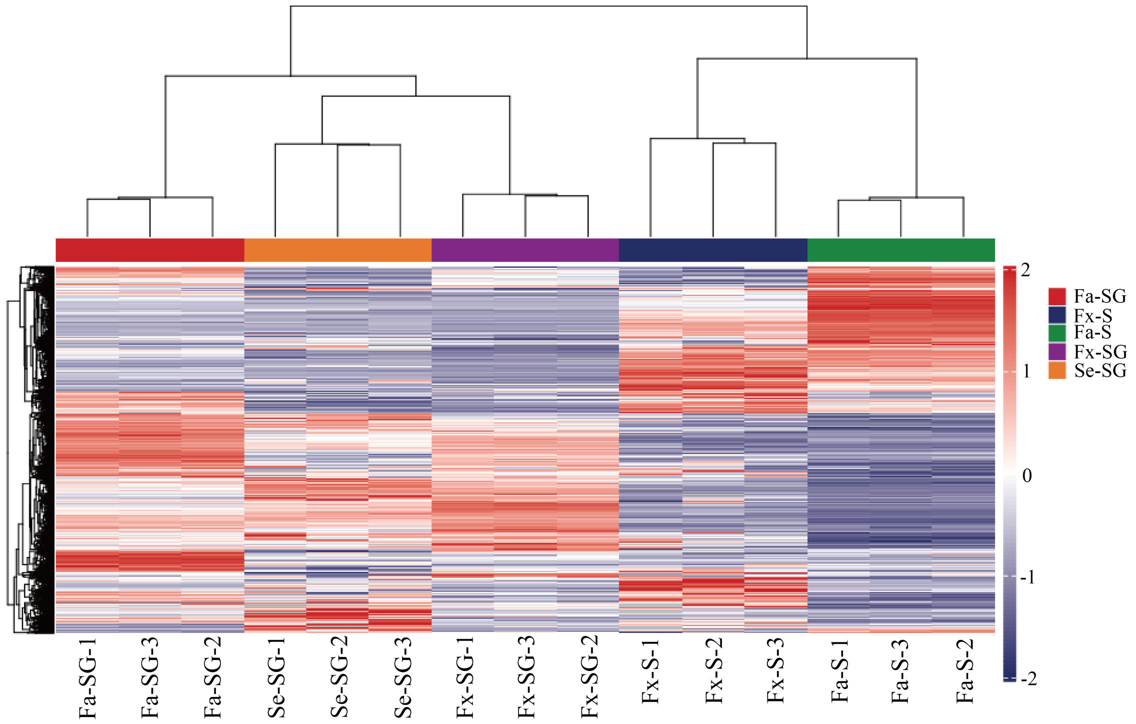


**Fig. S28. Hierarchical clustering of proteomic samples.** Unsupervised hierarchical clustering dendrogram and heatmap of quantified proteins across all 15 samples. Z-score-normalized intensity is shown (blue: low, red: high). Biological replicates cluster together, and the dendrogram reflects functional relationships among sample groups.

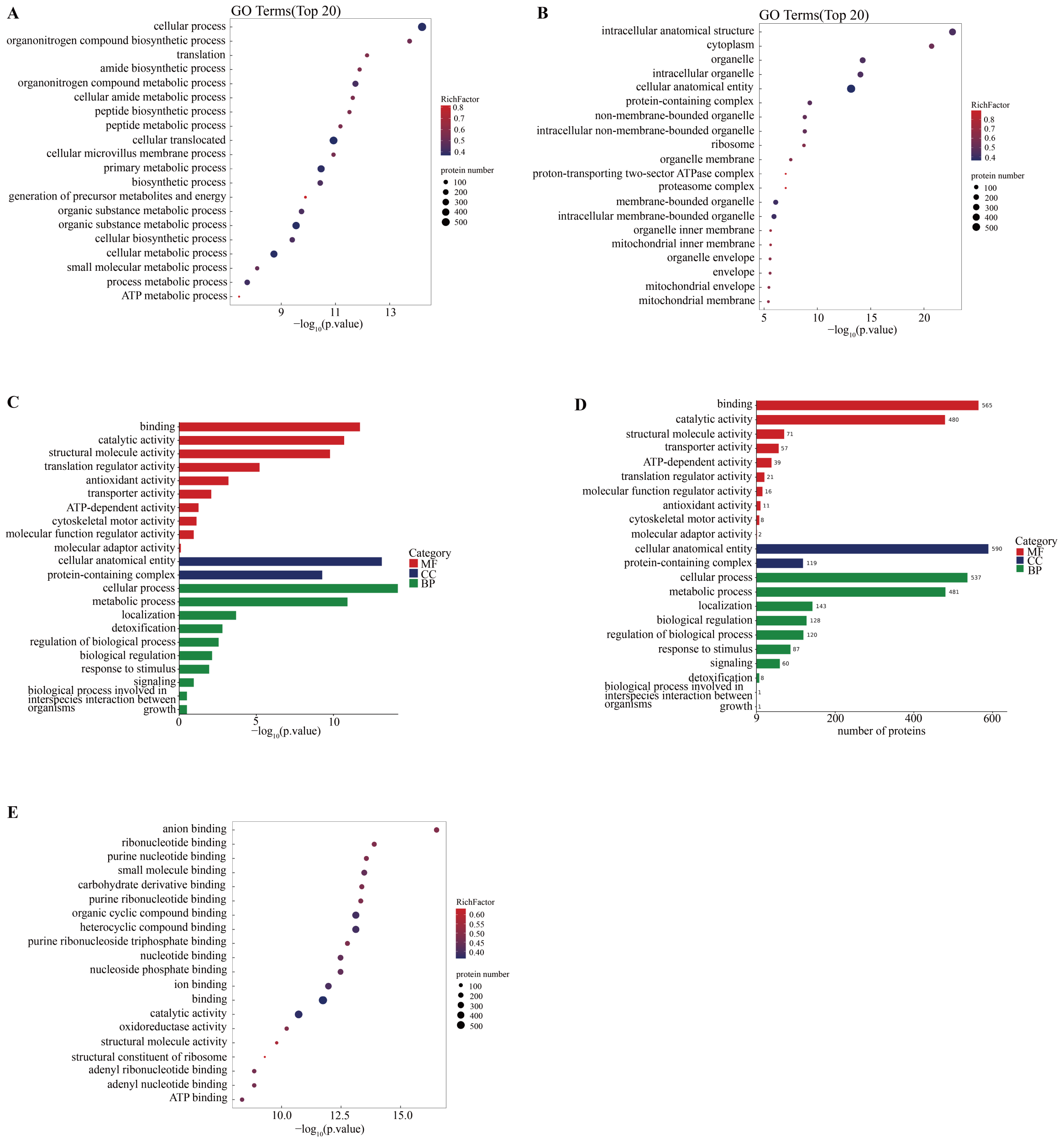


**Fig. S29. GO enrichment of differentially abundant proteins: one-way ANOVA across all five groups.** GO enrichment analysis of proteins differentially abundant across all five sample groups by one-way ANOVA ($P < 0.05$). (A) Biological process (BP); (B) Cellular component (CC); (C) Molecular function (MF); (D) GO Level 2 functional categories; (E) GO Level 2 overview.

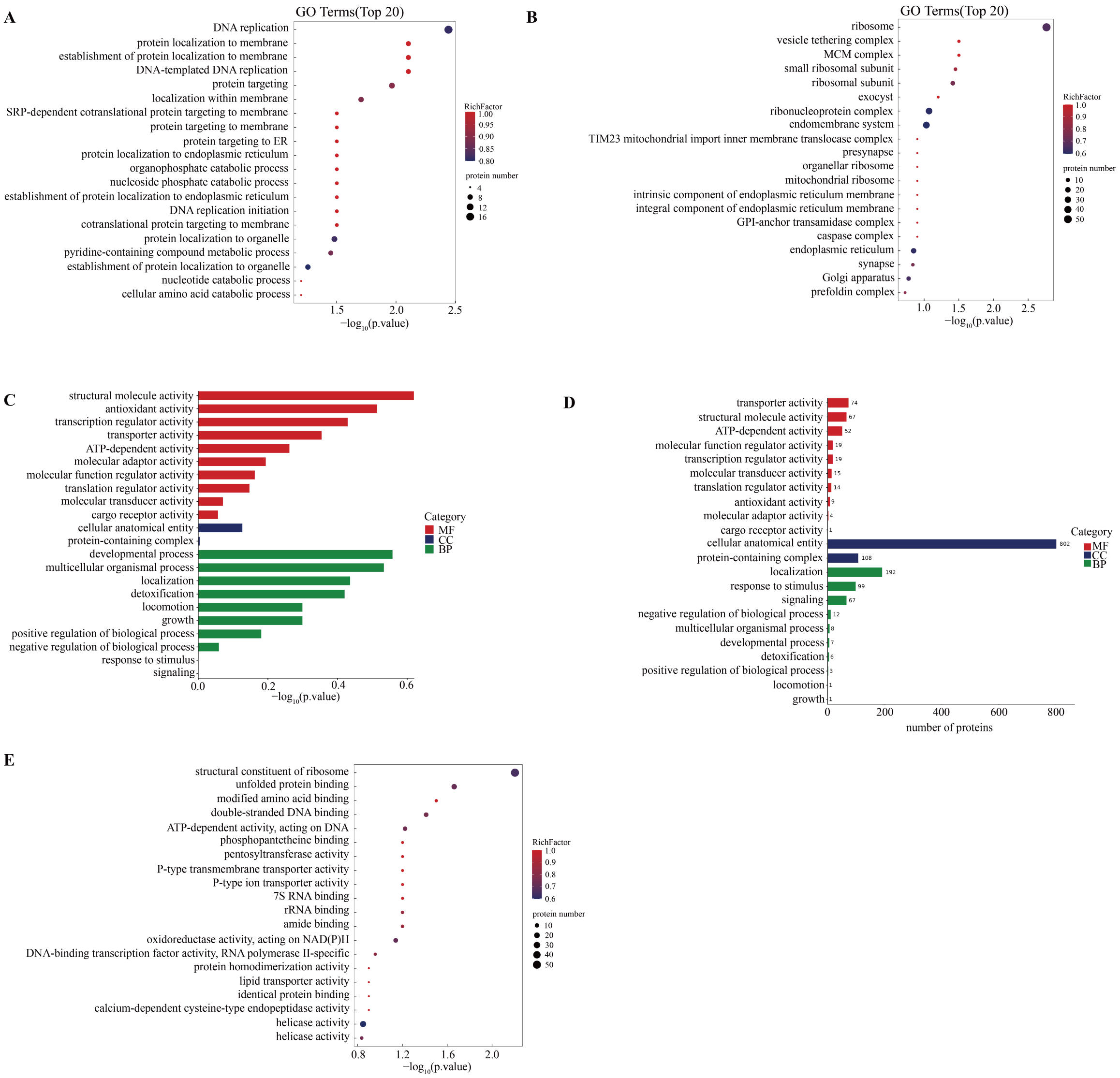


**Fig. S30. GO enrichment of differentially abundant proteins: Fa-SG vs Se-SG.** (A) Biological process (BP); (B) Cellular component (CC); (C) Molecular function (MF); (D) GO Level 2; (E) GO Level 2 overview. This comparison highlights functional differences between the salivary glands of gall-inducing fundatrigenia and non-galling sexuparae.

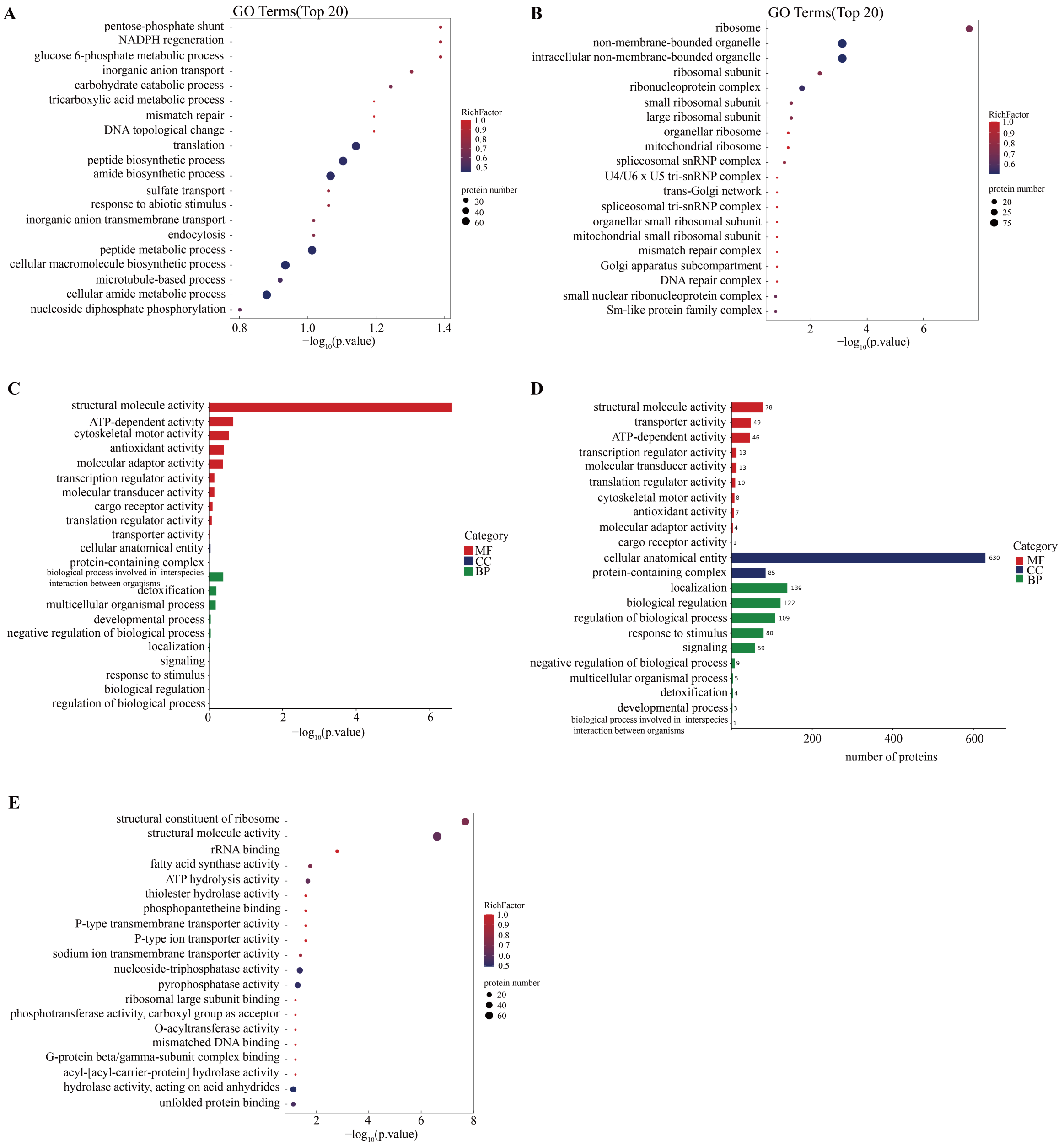


**Fig. S31. GO enrichment of differentially abundant proteins: Fx-S vs Fa-S.** (A) BP; (B) CC; (C) MF; (D) GO Level 2 with P values; (E) GO Level 2 overview. Comparison between fundatrix saliva and fundatrigenia saliva reveals functional shifts in secreted protein composition between the two gall-inducing morphs.

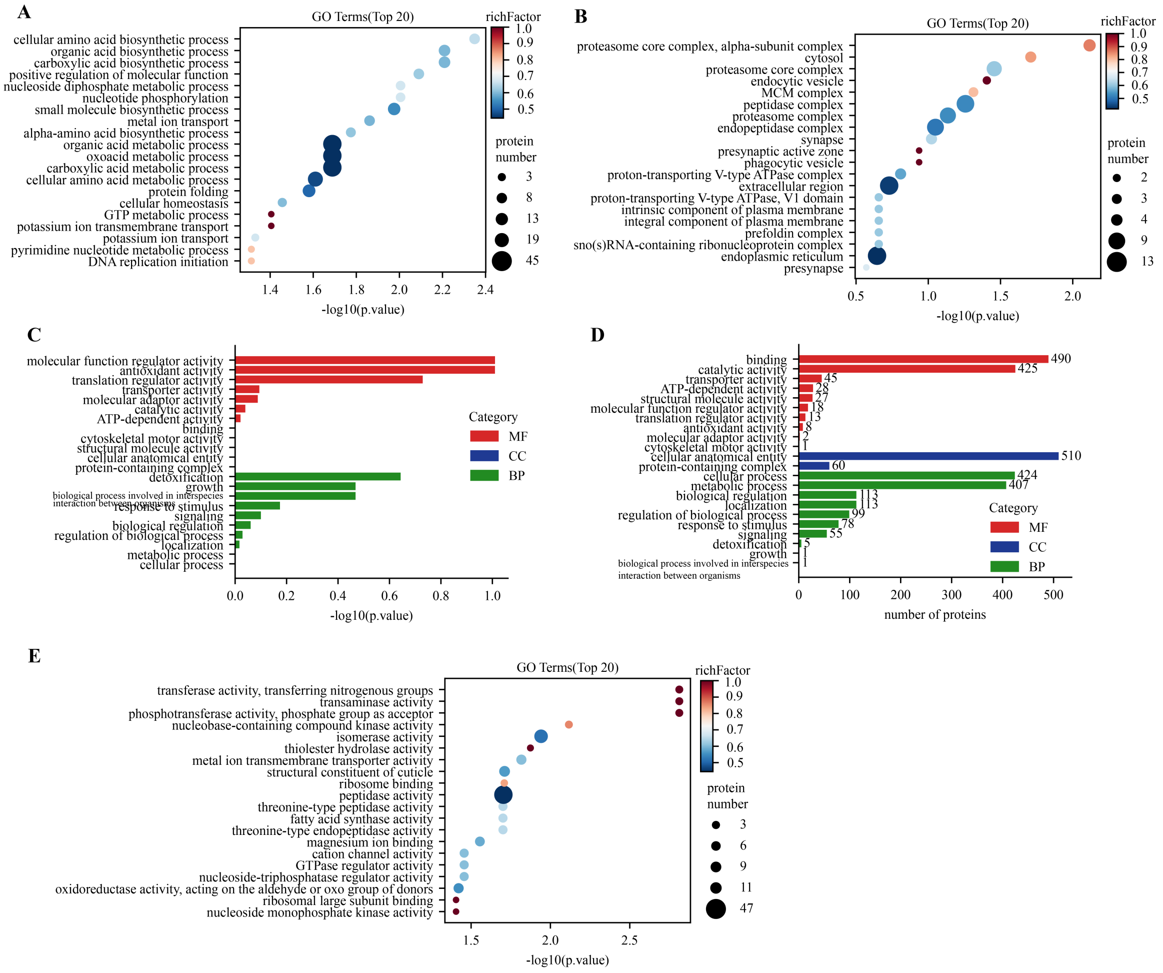


**Fig. S32. GO enrichment of differentially abundant proteins: Fx-SG vs Fa-SG.** (A) BP; (B) CC; (C) MF; (D) GO Level 2; (E) GO Level 2 overview.

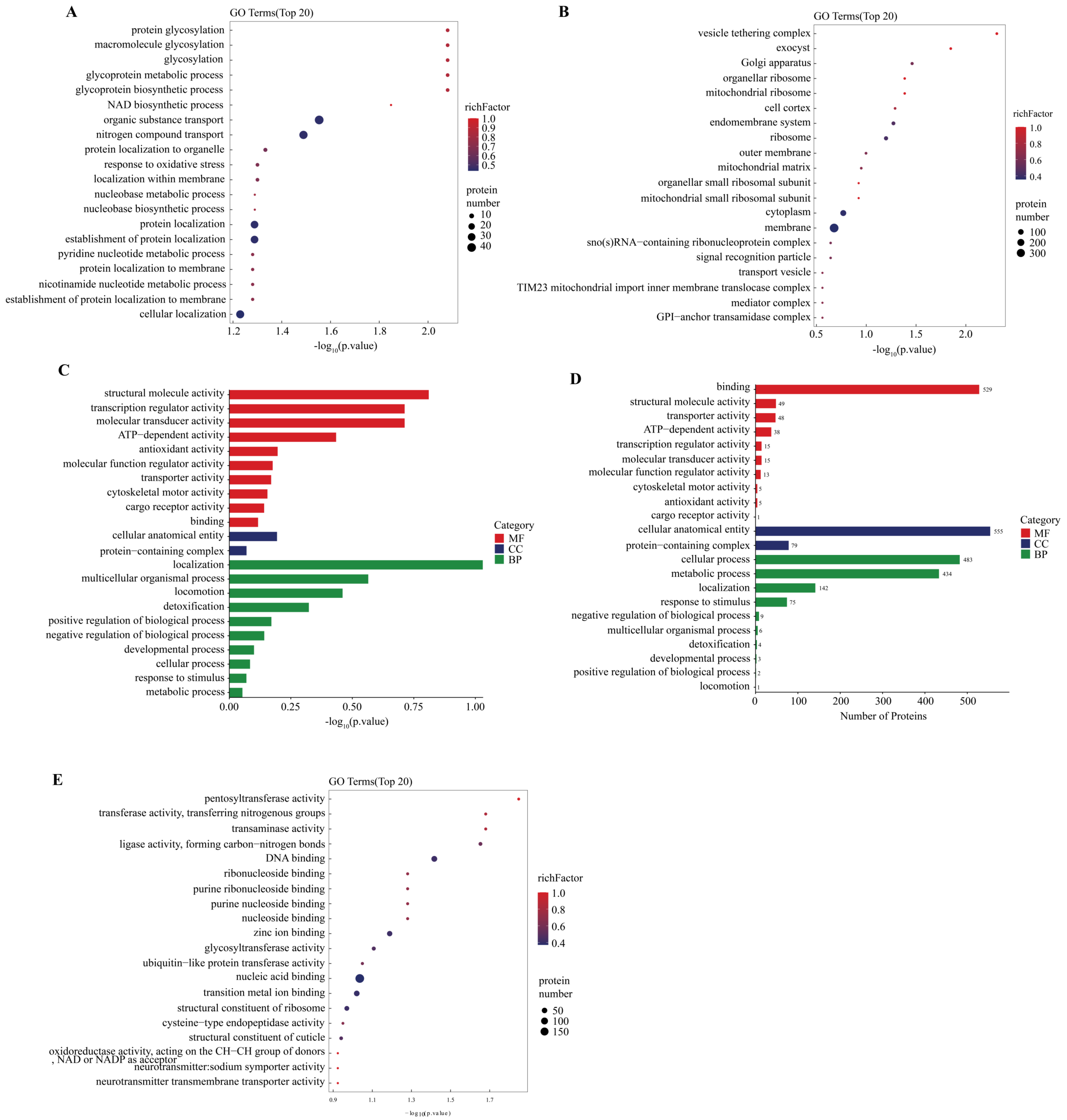


**Fig. S33. GO enrichment of differentially abundant proteins: Fx-SG vs Se-SG.** (A) BP; (B) CC; (C) MF; (D) GO Level 2; (E) GO Level 2 overview.

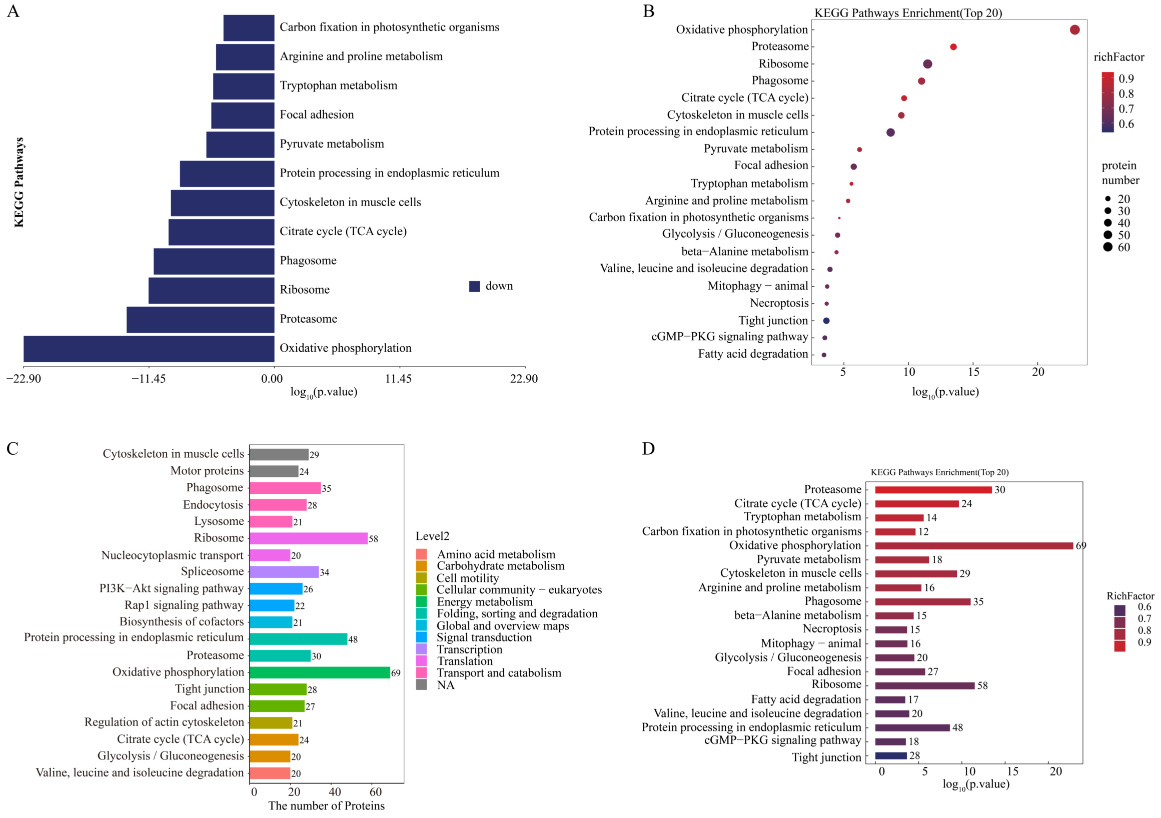


**Fig. S34. KEGG pathway enrichment of differentially abundant proteins: one-way ANOVA across all five groups.** (A) Up/down-regulated pathway classification; (B) KEGG bubble plot showing enriched pathways; (C) Level 2 statistics summarizing pathway categories; (D) Summary of enrichment results.

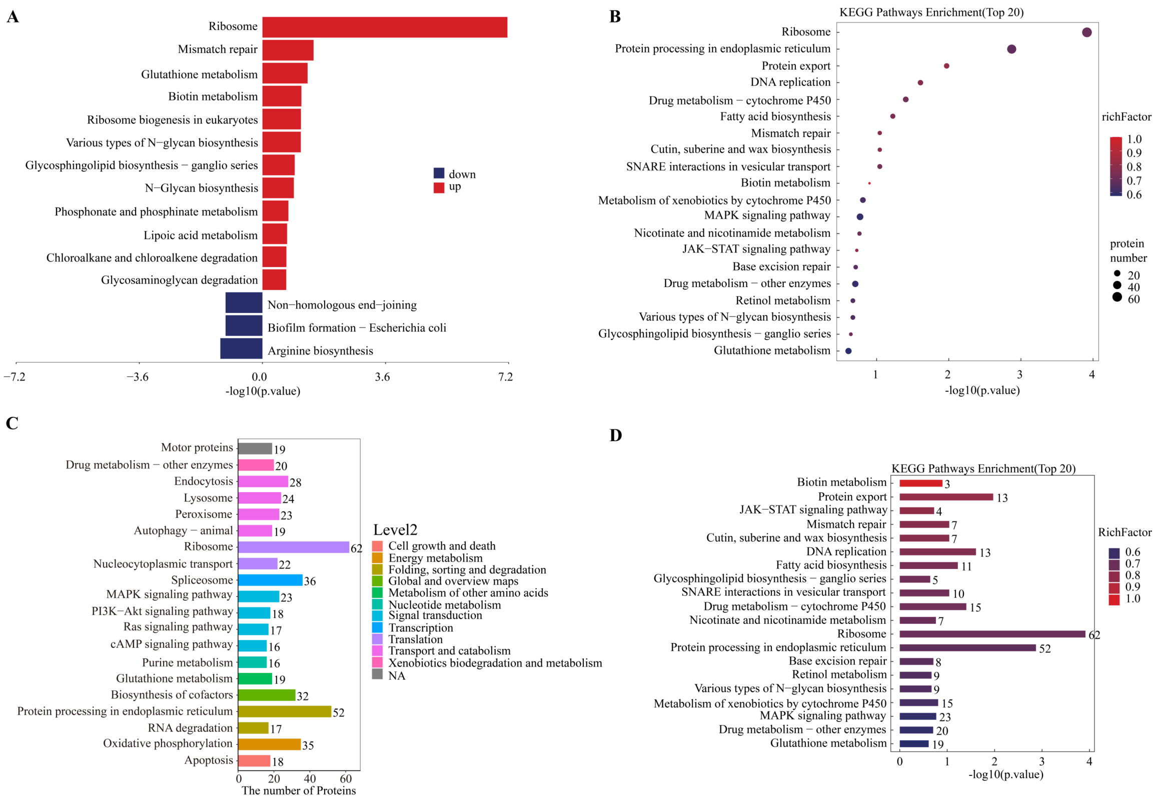


**Fig. S35. KEGG pathway enrichment of differentially abundant proteins: Fa-SG vs Se-SG.** (A) Up/down-regulated pathway classification; (B) Enrichment bubble plot; (C) Level 2 summary; (D) Overall summary. Enriched pathways reflect functional specialization of fundatrigenia salivary glands compared to the non-feeding sexuparae.

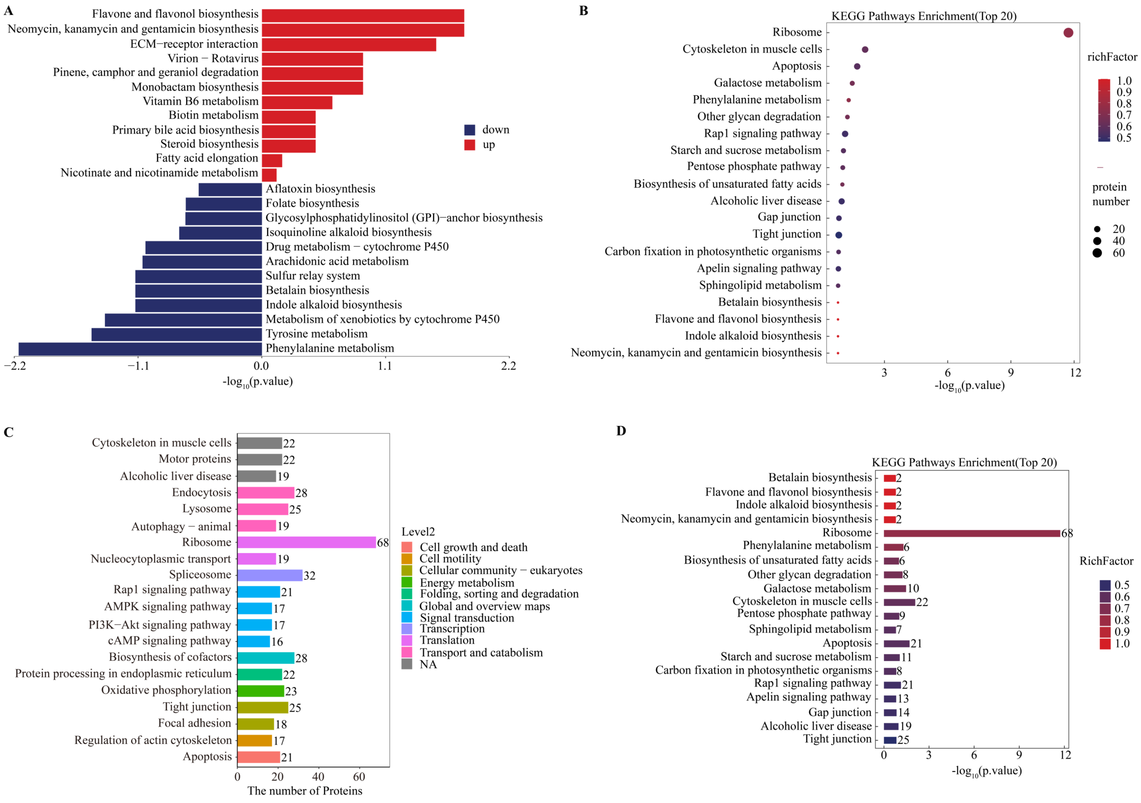


**Fig. S36. KEGG pathway enrichment of differentially abundant proteins: Fx-S vs Fa-S.** (A) Up/down-regulated pathway classification; (B) Enrichment bubble plot; (C) Level 2 summary; (D) Overall summary.

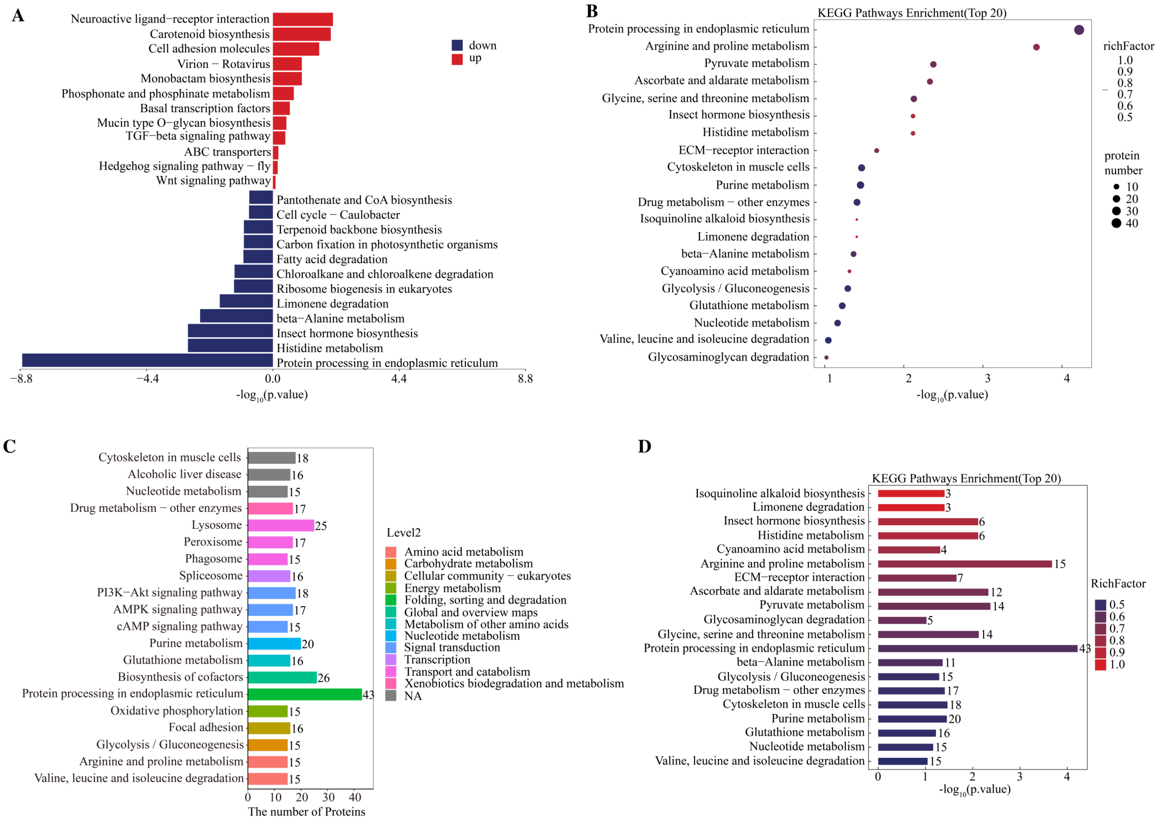


**Fig. S37. KEGG pathway enrichment of differentially abundant proteins: Fx-SG vs Fa-SG.** (A) Up/down-regulated pathway classification; (B) Enrichment bubble plot; (C) Level 2 summary; (D) Overall summary.

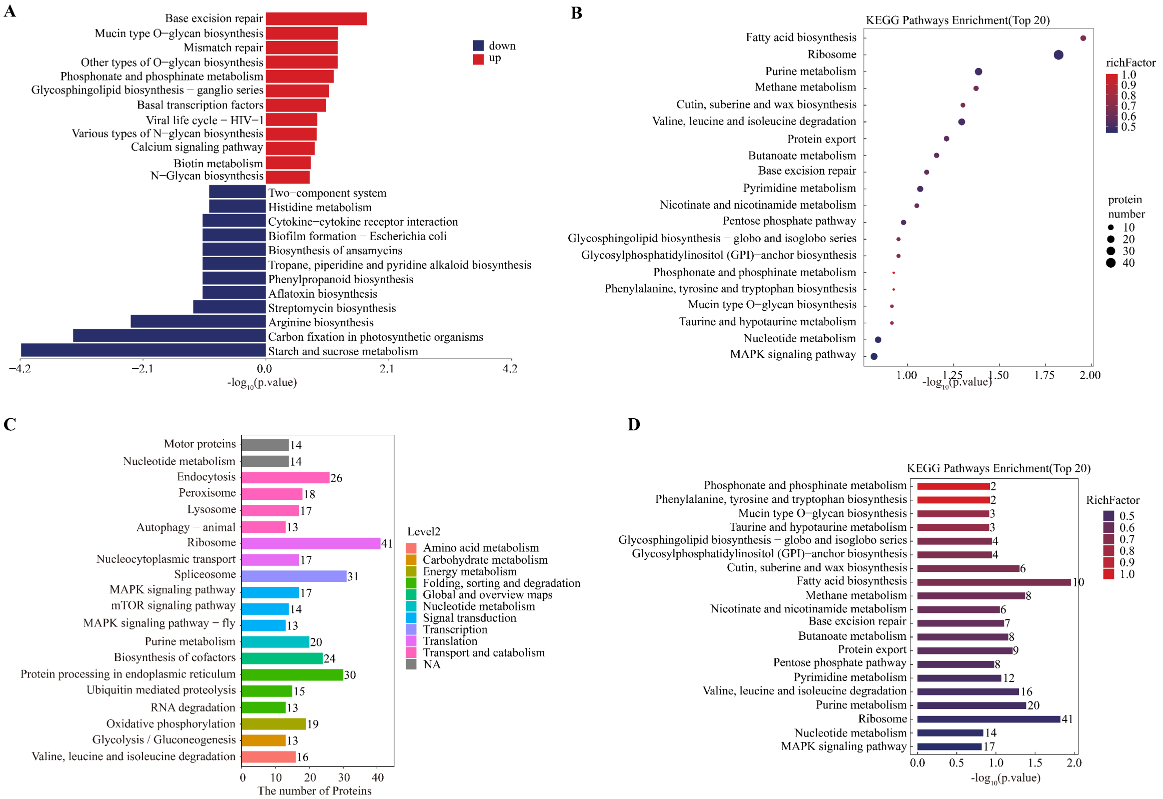


**Fig. S38. KEGG pathway enrichment of differentially abundant proteins: Fx-SG vs Se-SG.** (A) Up/down-regulated pathway classification; (B) Enrichment bubble plot; (C) Level 2 summary; (D) Overall summary.

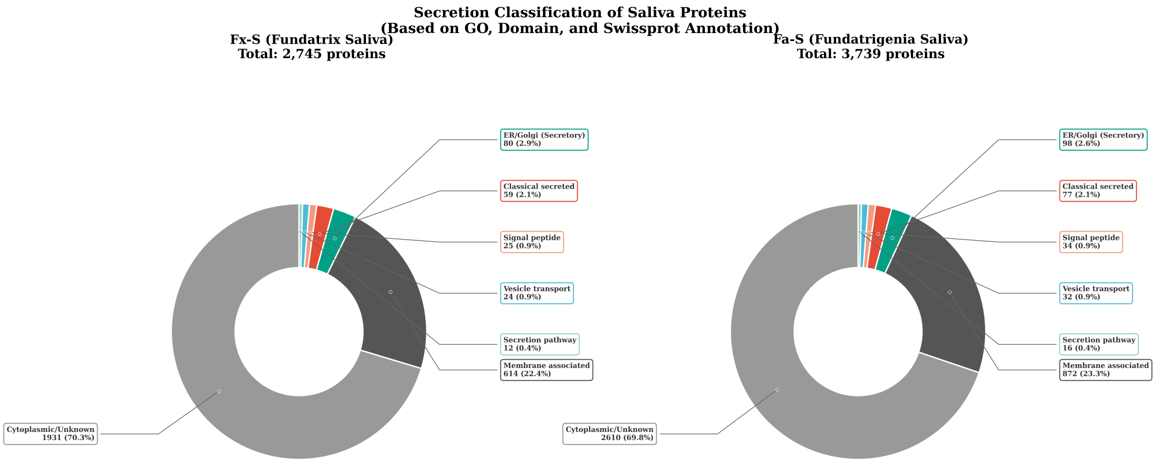


**Fig. S39. Secretion classification of saliva proteins based on GO, domain, and Swissprot annotation.** Donut charts showing the distribution of proteins in Fx-S (fundatrix saliva, left, n = 2,745) and Fa-S (fundatrigenia saliva, right, n = 3,739) samples. Proteins were classified into seven categories: Cytoplasmic/Unknown, Membrane associated, ER/Golgi localized (Secretory pathway), Classical secreted (Signal peptide + Extracellular), Signal peptide containing, Vesicle transport associated, and Secretion pathway associated. The majority of salivary proteins (~70%) were classified as cytoplasmic or unknown, while approximately 23% were membrane-associated. Secretory pathway-related proteins collectively accounted for ~7% of the total proteome in both samples.

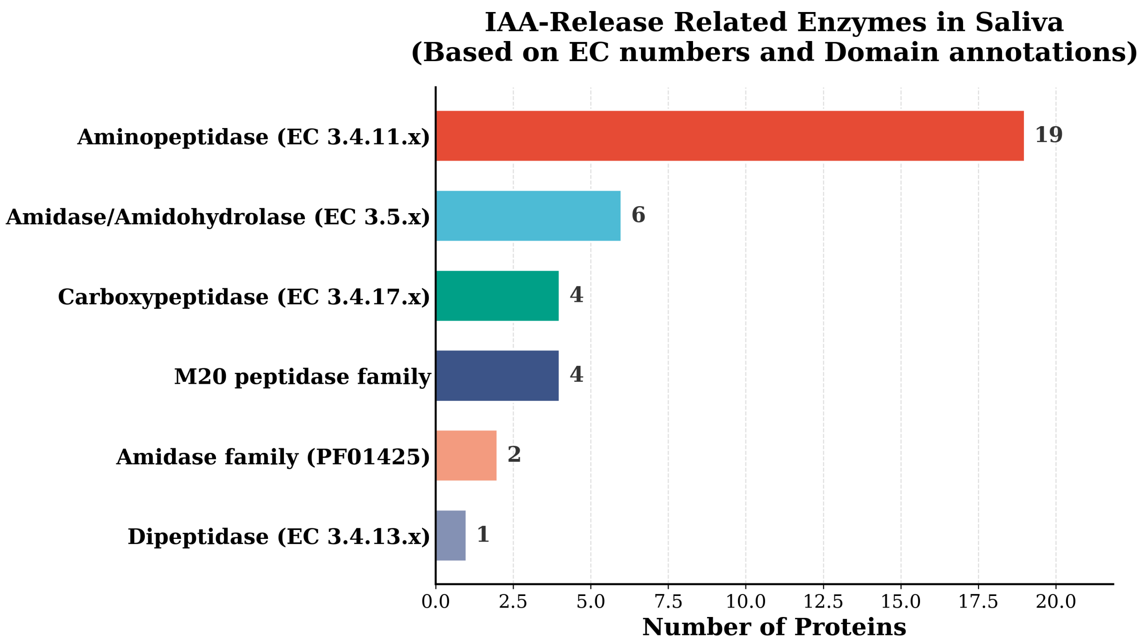


**Fig. S40. Distribution of IAA-release related enzymes identified in aphid saliva proteome.** Horizontal bar chart showing the number of proteins in each enzyme category based on EC numbers and domain annotations. Aminopeptidases (EC 3.4.11.x) represent the largest group with 19 proteins, followed by Amidase/Amidohydrolase (EC 3.5.x, 6 proteins), Carboxypeptidase (EC 3.4.17.x, 4 proteins), M20 peptidase family (4 proteins), Amidase family (PF01425, 2 proteins), and Dipeptidase (EC 3.4.13.x, 1 protein). These enzymes are potentially involved in the release of free IAA from conjugated forms.

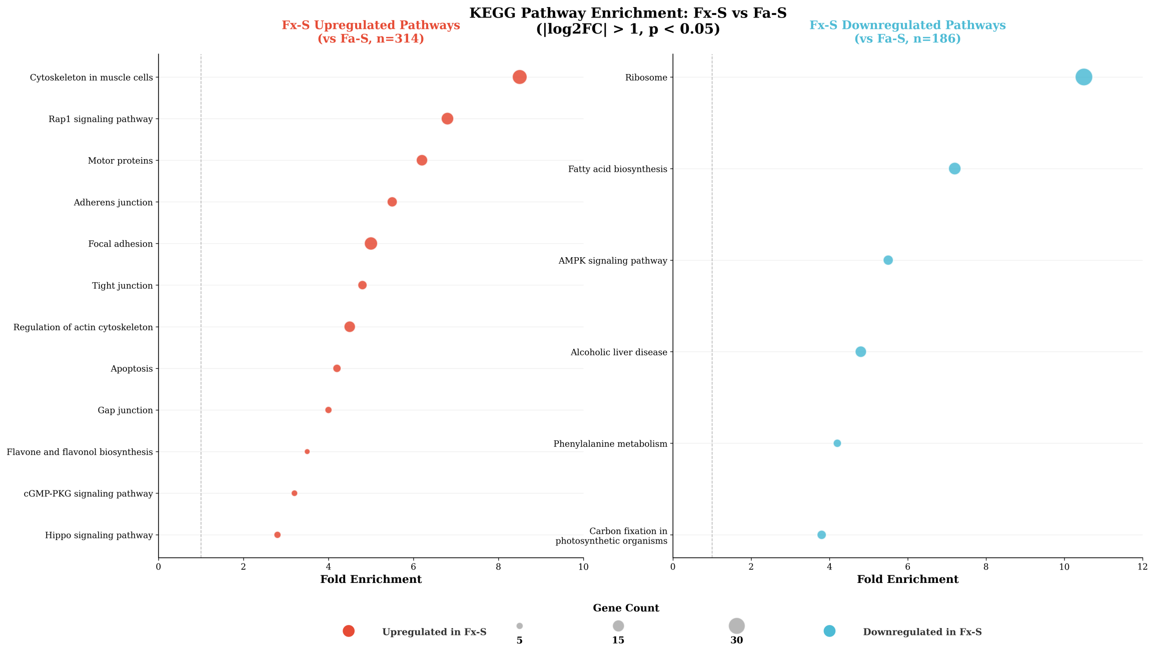


**Fig. S41. KEGG pathway enrichment analysis comparing differentially expressed proteins between Fx-S and Fa-S.** Left panel: Pathways upregulated in Fx-S (vs Fa-S, n = 314 proteins), shown in coral/red. Right panel: Pathways downregulated in Fx-S (i.e., upregulated in Fa-S, n = 186 proteins), shown in cyan/blue. Bubble size represents gene count; x-axis indicates fold enrichment. Fx-S-upregulated pathways are enriched in cytoskeleton-related processes (regulation of actin cytoskeleton, motor proteins) and cell junction pathways (adherens junction, focal adhesion). Fa-S-upregulated pathways show enrichment in translation (ribosome), lipid metabolism (fatty acid biosynthesis), and metabolic processes (AMPK signaling, phenylalanine metabolism). Thresholds: $|\log_2 FC| > 1$, $P < 0.05$.

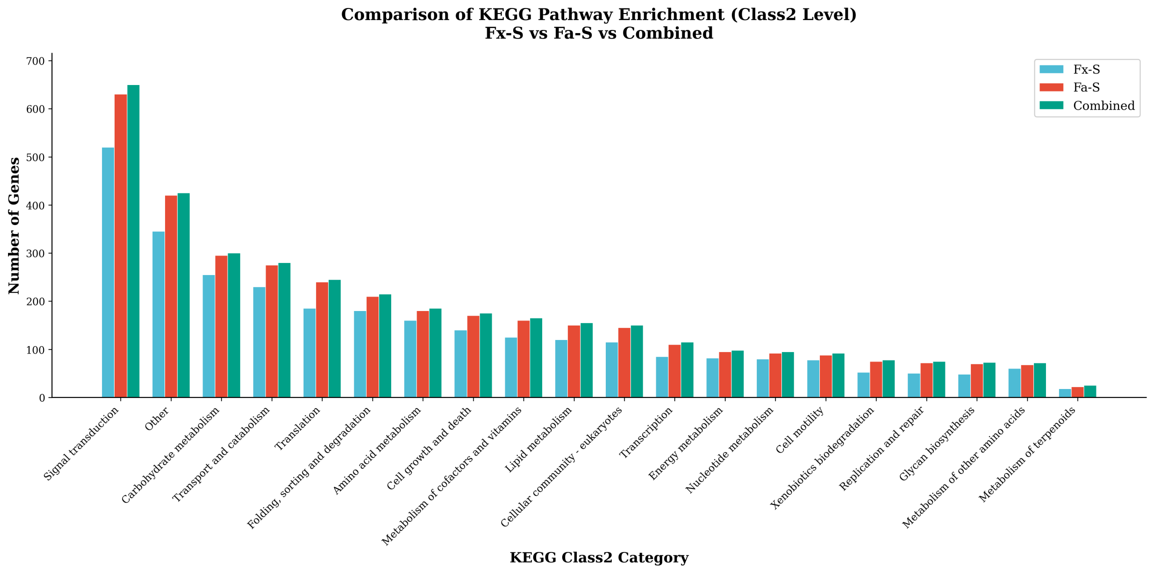


**Fig. S42. Comparison of KEGG pathway enrichment at Class 2 level among Fx-S, Fa-S, and Combined samples.** Bar chart showing the number of genes/proteins assigned to each functional category. Blue bars: Fx-S; Red/coral bars: Fa-S; Green bars: Combined. Signal transduction represents the most enriched category (>500 genes), followed by carbohydrate metabolism and transport and catabolism. Fa-S generally shows higher gene counts across most categories compared to Fx-S, consistent with its larger proteome size.

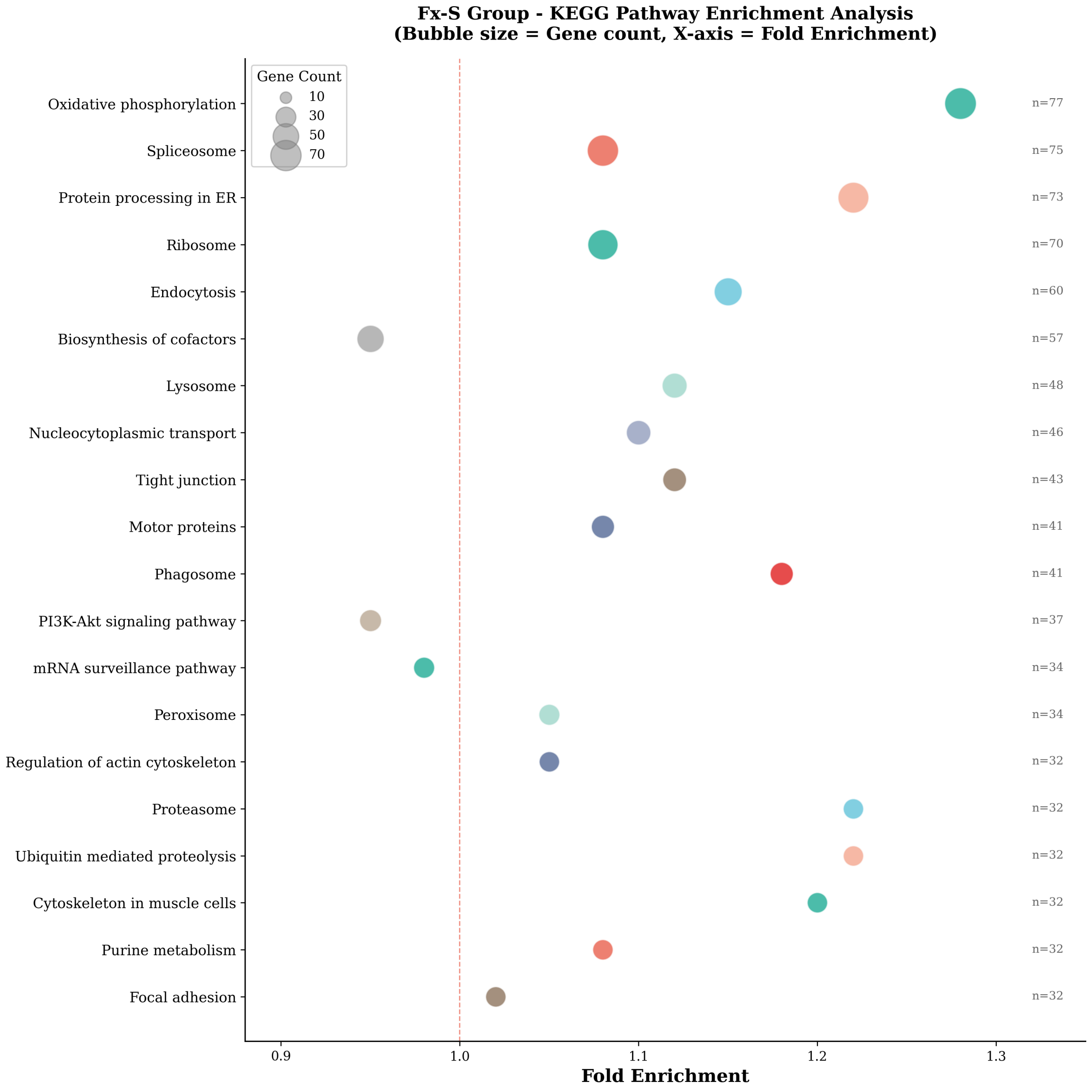


**Fig. S43. KEGG pathway enrichment analysis for Fx-S (fundatrix saliva) proteome.** Bubble plot displaying the top 20 enriched pathways ranked by gene count. The x-axis represents fold enrichment (ratio of observed to expected), with the red dashed line indicating fold enrichment = 1.0. Bubble size corresponds to gene count. Key enriched pathways include: oxidative phosphorylation (n = 77, highest enrichment), protein processing in endoplasmic reticulum (n = 73), spliceosome (n = 75), ribosome (n = 70), and endocytosis (n = 60).

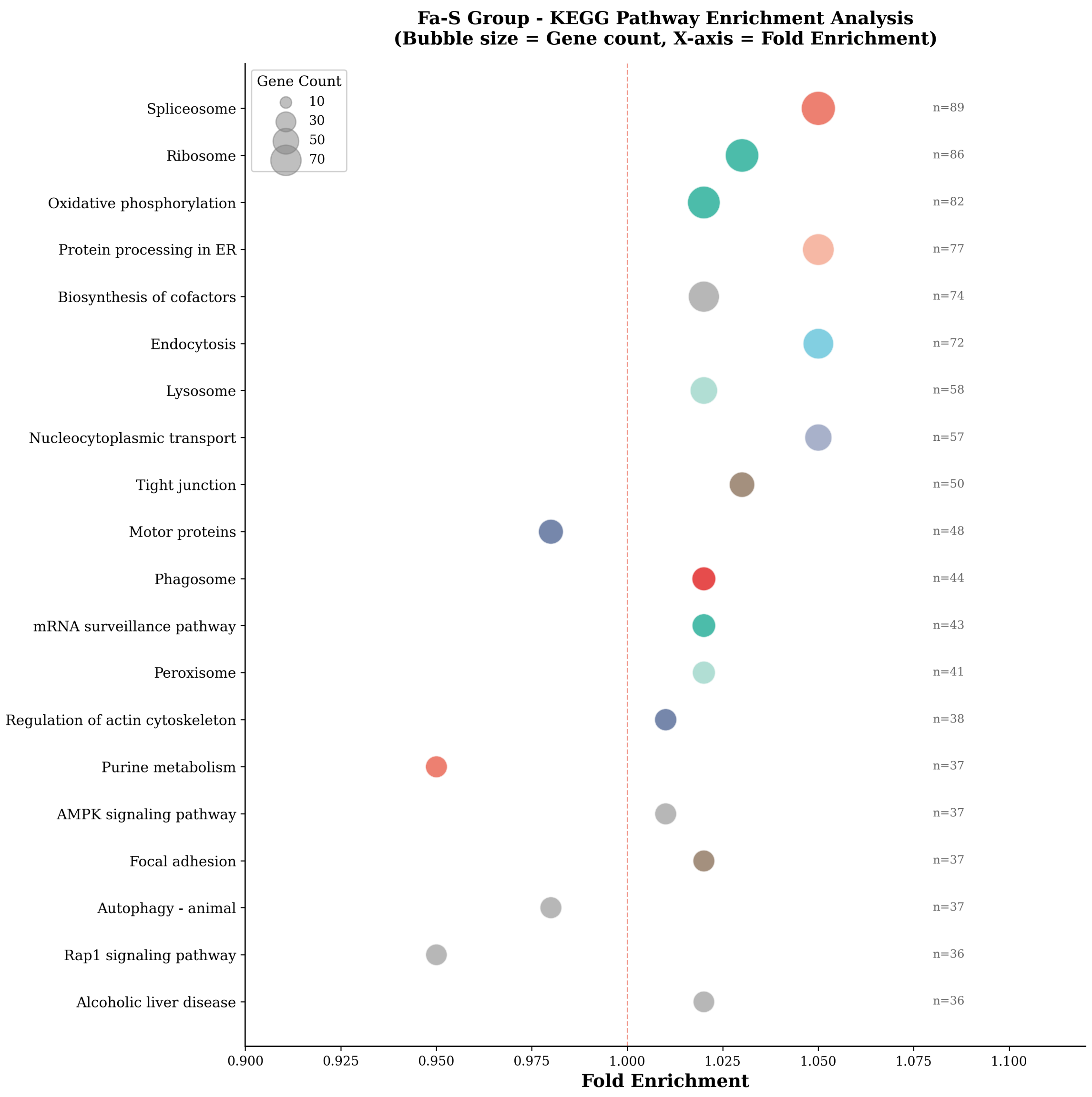


**Fig. S44. KEGG pathway enrichment analysis for Fa-S (fundatrigenia saliva) proteome.** Bubble plot displaying the top 20 enriched pathways ranked by gene count. Compared to Fx-S, Fa-S shows higher gene counts across most pathways, with spliceosome (n = 89), ribosome (n = 86), and oxidative phosphorylation (n = 82) being the top three. Notable differences from Fx-S include stronger enrichment in biosynthesis of cofactors (n = 74), endocytosis (n = 72), and nucleocytoplasmic transport (n = 57). The overall fold enrichment values are lower than Fx-S, reflecting the larger background proteome size.

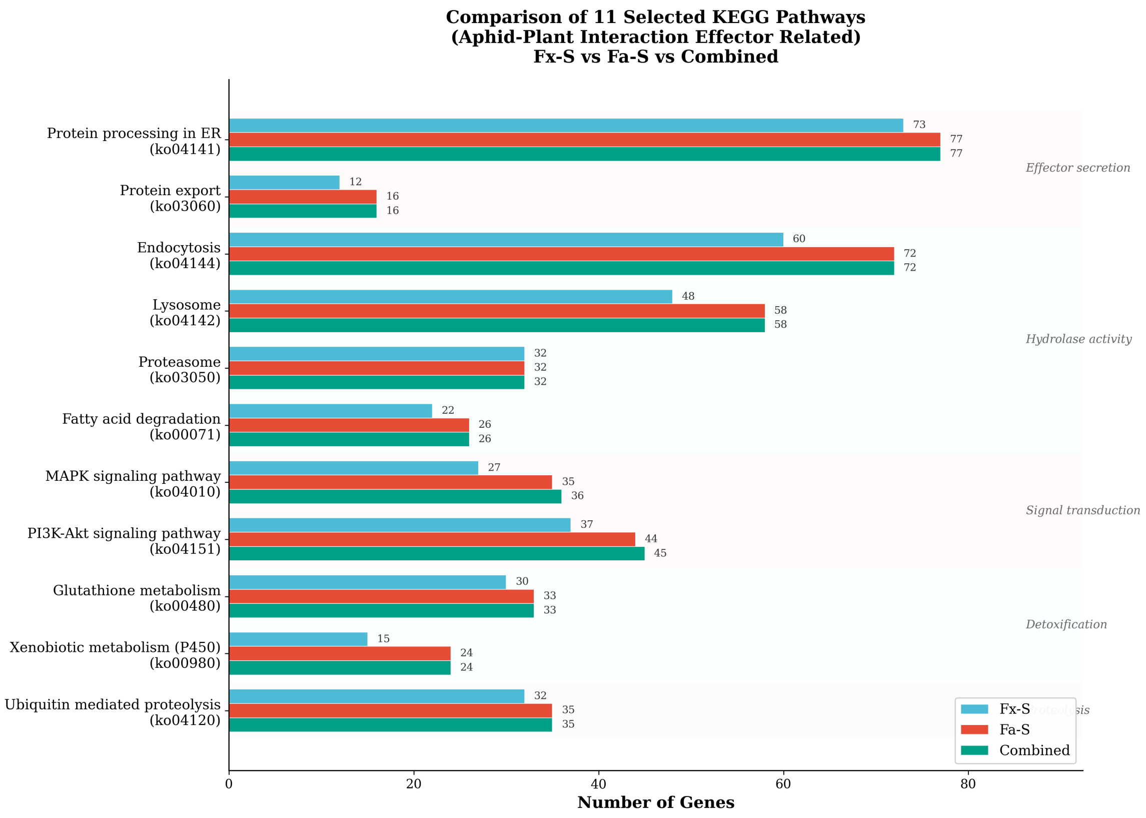


**Fig. S45. Comparison of 11 selected KEGG pathways related to aphid–plant interaction and effector function among Fx-S, Fa-S, and Combined samples.** Horizontal grouped bar chart with pathway names and KEGG IDs. Blue bars: Fx-S; Red/coral bars: Fa-S; Green bars: Combined. Numbers indicate gene counts. Pathways are grouped by functional category: Effector secretion (Protein processing in ER, Protein export), Hydrolase activity (Endocytosis, Lysosome, Proteasome, Fatty acid degradation), Signal transduction (MAPK signaling pathway, PI3K-Akt signaling pathway), Detoxification (Glutathione metabolism, Xenobiotic metabolism), and Proteolysis (Ubiquitin mediated proteolysis). Protein processing in ER shows the highest gene counts (73–77), followed by Endocytosis (60–72) and Lysosome (48–58).

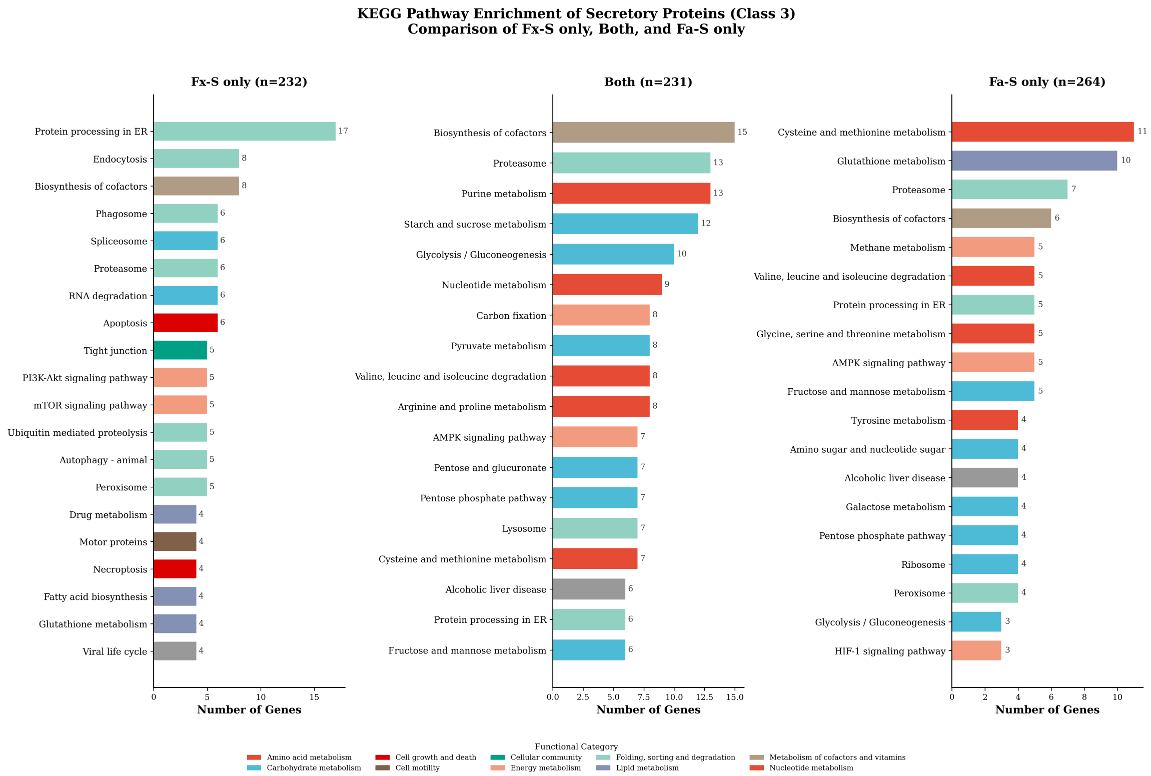


**Fig. S46. KEGG pathway enrichment analysis of secretory proteins comparing Fx-S only, Both (shared), and Fa-S only groups.** Three-panel horizontal bar chart showing the top pathways in each category. Left panel: Fx-S only (n = 232 proteins), dominated by Protein processing in ER (17 genes), Endocytosis (8), and Biosynthesis of cofactors (8). Middle panel: Both/shared (n = 231 proteins), enriched in Biosynthesis of cofactors (15), Proteasome (13), and Purine metabolism (13). Right panel: Fa-S only (n = 264 proteins), characterized by Cysteine and methionine metabolism (11), Glutathione metabolism (10), and Proteasome (7). Bar colors indicate functional categories. Fx-S-specific secretory proteins are enriched in protein folding and secretion pathways, while Fa-S-specific proteins show enrichment in amino acid metabolism and detoxification pathways.

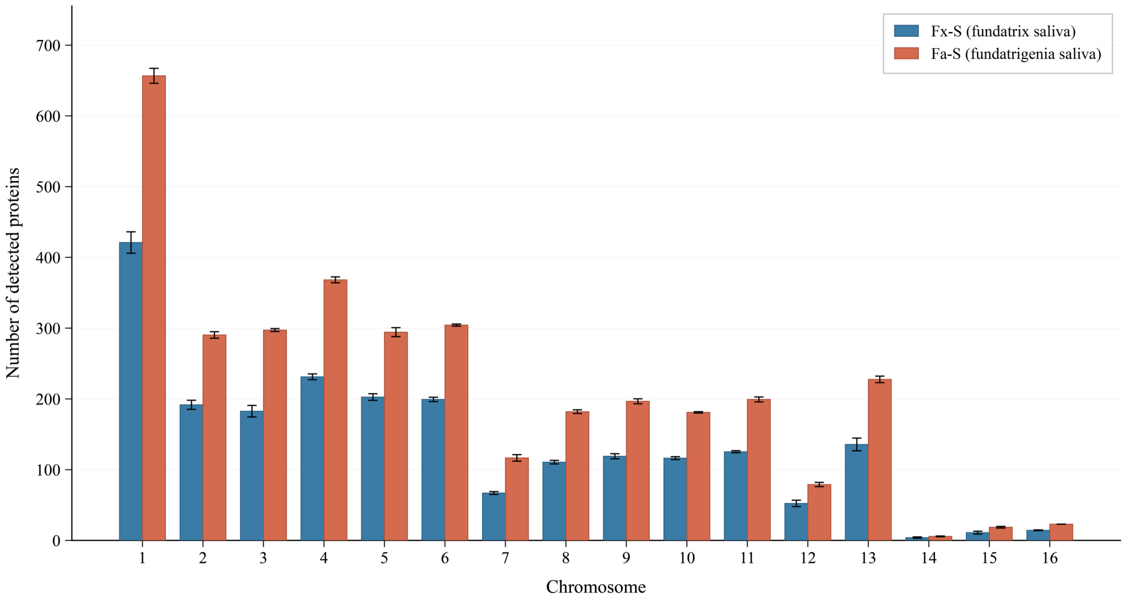


**Fig. S47. Chromosomal distribution of saliva proteins detected in fundatrix and fundatrigenia.** Grouped bar chart showing the number of proteins detected in Fx-S (fundatrix saliva, steel blue) and Fa-S (fundatrigenia saliva, terracotta) across the 16 chromosomes (chr1–chr16) of S. chinensis. Values represent mean ± s.d. of three biological replicates. A total of 2,727 (Fx-S) and 3,715 (Fa-S) proteins were mapped to chr1–chr16. Chromosome 1 harbored the greatest number of detected proteins in both samples (Fx-S: 421.0 ± 15.1; Fa-S: 656.7 ± 10.6), consistent with its disproportionately large gene content (5,986 genes, 33.08% of the genome). Fa-S consistently yielded higher protein counts than Fx-S on every chromosome, reflecting its broader proteomic complexity.

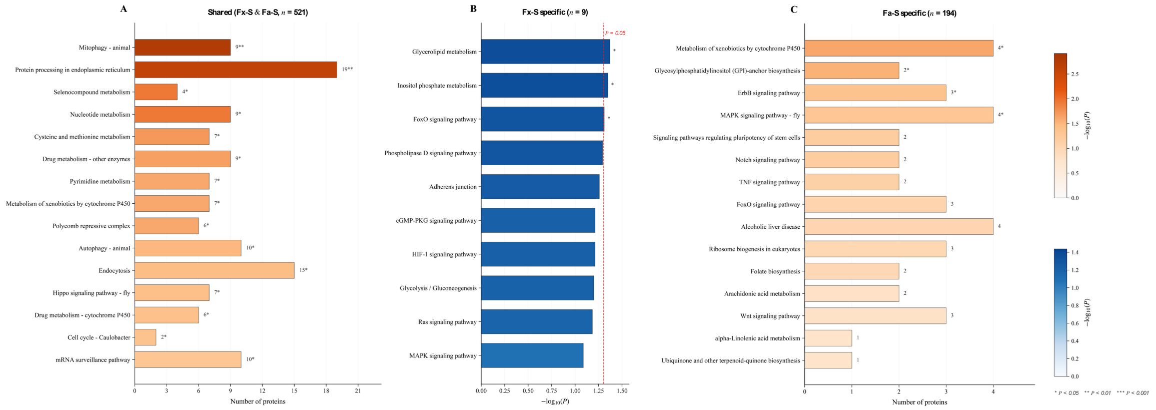


**Fig. S48. KEGG pathway enrichment of chromosome 1–encoded saliva proteins shared between or specific to fundatrix and fundatrigenia.** Horizontal bar charts displaying KEGG pathway enrichment results for three protein subsets encoded on chromosome 1: (A) Proteins shared between Fx-S and Fa-S (n = 521). Bar length indicates the number of proteins; color gradient encodes $-\log_{10}$(P value). Top enriched pathways include Protein processing in endoplasmic reticulum (19 proteins, P = $2.49 \times 10^{-3}$) and Mitophagy—animal (9 proteins, P = $1.68 \times 10^{-3}$). (B) Proteins detected exclusively in Fx-S (n = 9). Three pathways reached nominal significance: Glycerolipid metabolism, Inositol phosphate metabolism, and FoxO signaling pathway. (C) Proteins detected exclusively in Fa-S (n = 194). The most significantly enriched pathways were Metabolism of xenobiotics by cytochrome P450 (4 proteins, P = $2.38 \times 10^{-2}$), GPI-anchor biosynthesis (2 proteins, P = $2.89 \times 10^{-2}$), and ErbB signaling pathway (3 proteins, P = $4.24 \times 10^{-2}$). Enrichment was assessed by one-sided hypergeometric test against the full proteomic dataset (4,141 proteins) as background, with Benjamini–Hochberg correction. *$P < 0.05$, **$P < 0.01$, ***$P < 0.001$.

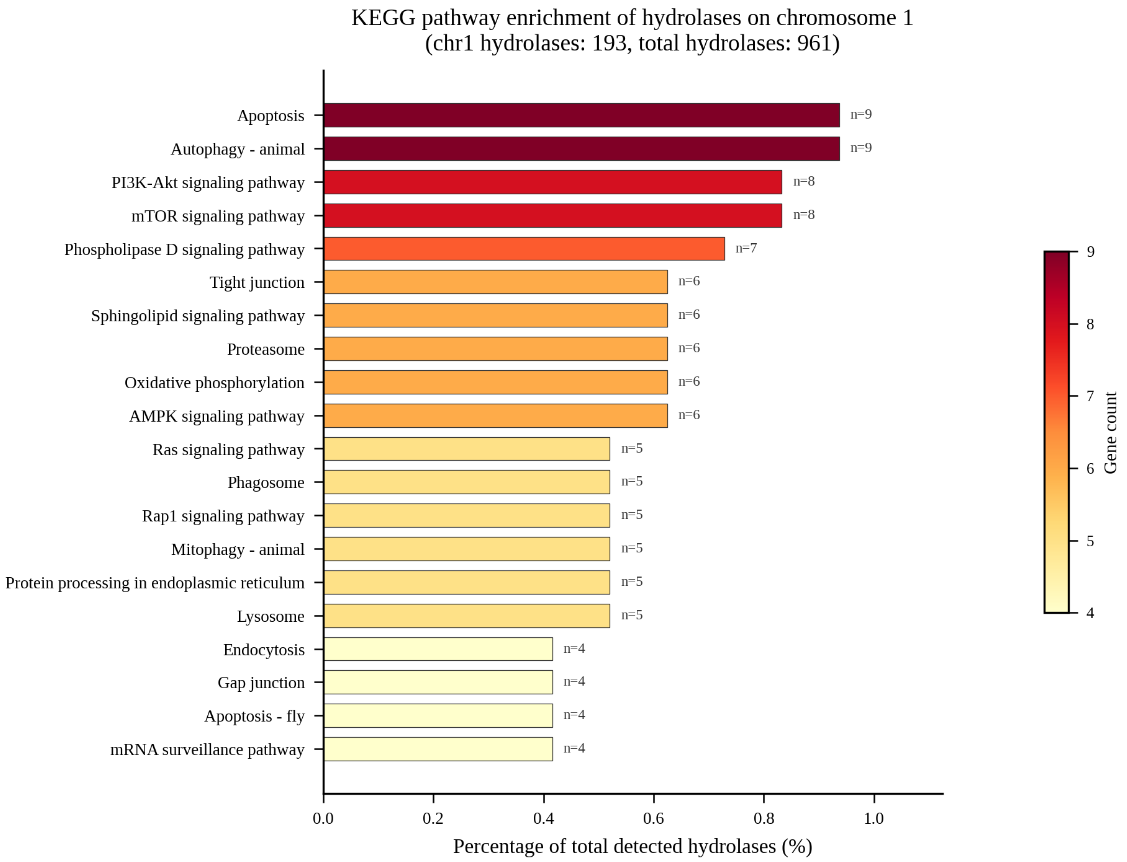


**Figure S49. KEGG pathway enrichment analysis of hydrolases located on chromosome 1 of *Schlechtendalia chinensis*.** The horizontal bar chart displays the top 20 enriched KEGG pathways for the 193 hydrolases identified on chr1 from the proteome. The x-axis represents the percentage of each pathway relative to the total 961 hydrolases detected across the genome, and the bar color gradient (yellow to dark red) indicates gene count. Gene numbers (n) are annotated beside each bar. Apoptosis and autophagy–animal pathways ranked highest (n = 9 each), followed by PI3K-Akt signaling (n = 8), mTOR signaling (n = 8), and phospholipase D signaling (n = 7), indicating that chr1 hydrolases are predominantly associated with programmed cell death regulation, cellular recycling, and signal transduction.

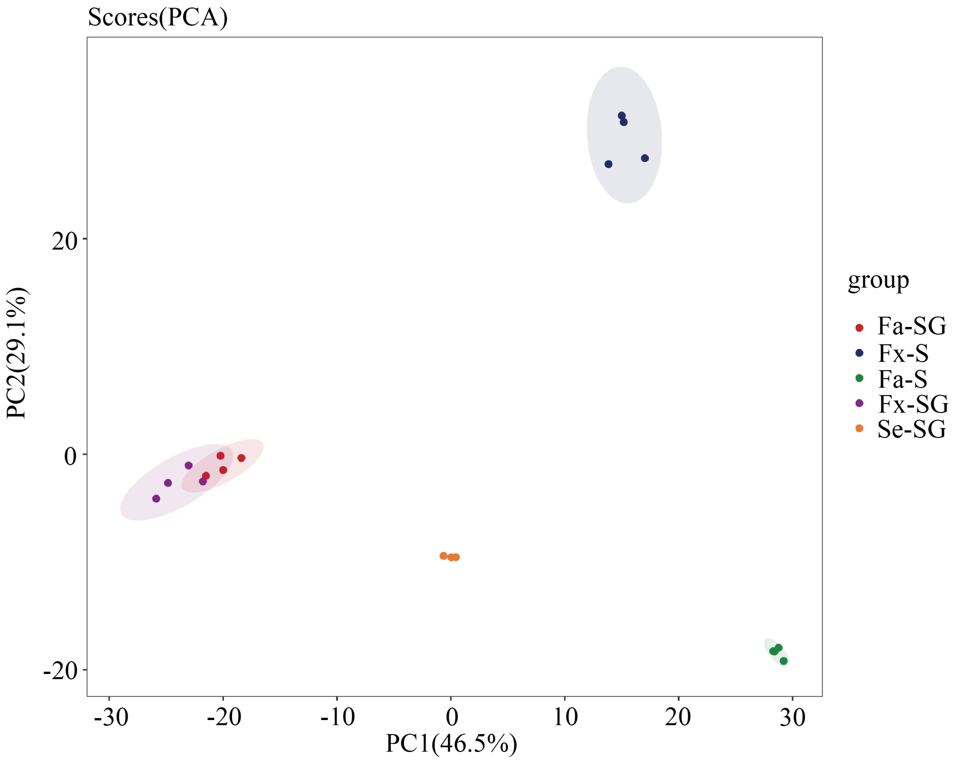


**Fig. S50. PCA of untargeted metabolomics profiles (negative ionization mode).** PCA score plot in negative ionization mode across all 15 samples (3 replicates × 5 groups). Ellipses indicate 95% confidence intervals. Clear separation between salivary gland and saliva samples is observed, consistent with the proteomic PCA (Fig. S20).

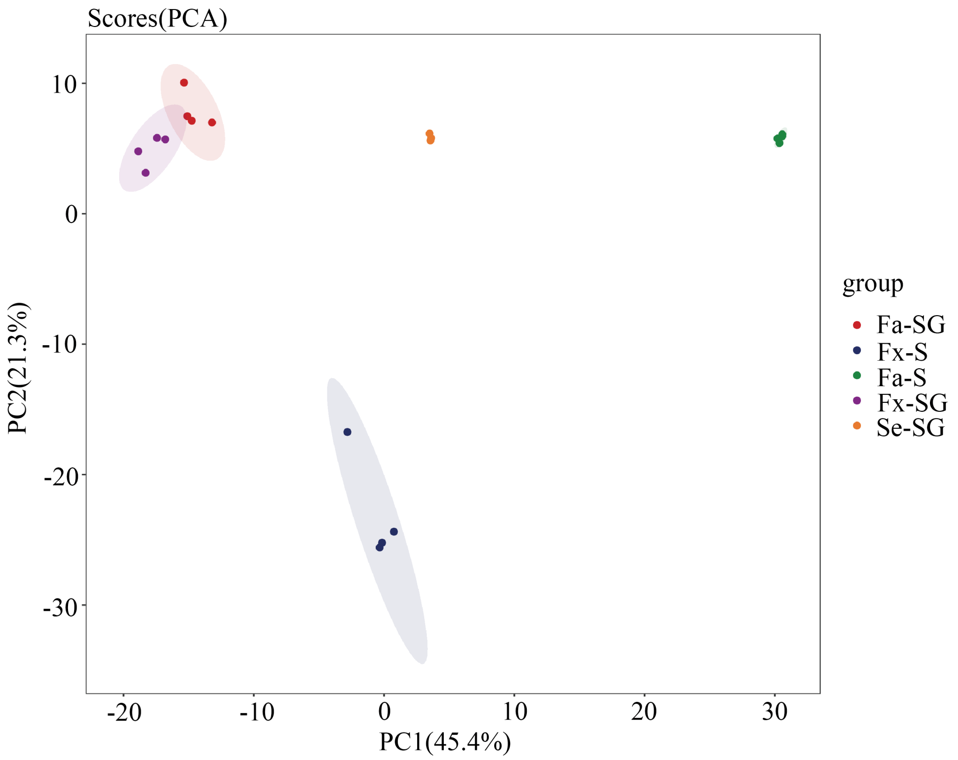


**Fig. S51. PCA of untargeted metabolomics profiles (positive ionization mode).** PCA score plot in positive ionization mode. The separation pattern is consistent with negative mode (Fig. S50), confirming robust metabolomic group differences independent of ionization polarity.

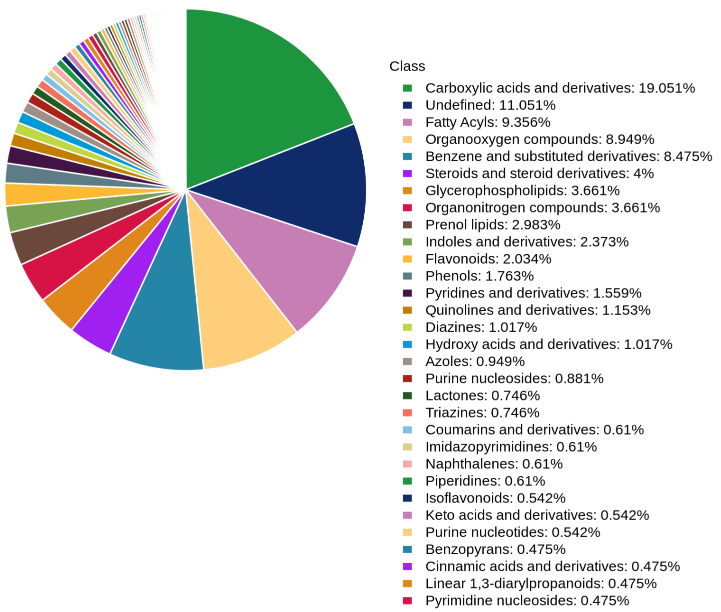


**Fig. S52. Chemical class composition of all detected metabolites.** Pie chart of class-level classification of 1,475 metabolites. Carboxylic acids and derivatives (19.05%), undefined (11.05%), fatty acyls (9.36%), organooxygen compounds (8.95%), and benzene derivatives (8.48%) are the most represented classes.

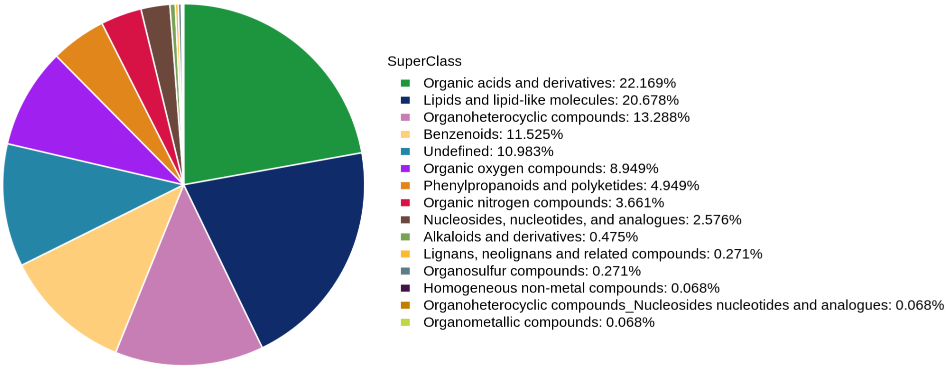


**Fig. S53. Chemical superclass composition of metabolites assigned to a superclass.** Pie chart of superclass-level classification of the 1,313 metabolites assigned to a superclass (of 1,475 detected in total). Organic acids and derivatives (327; 24.90%), lipids and lipid-like molecules (305; 23.23%), organoheterocyclic compounds (196; 14.93%) and benzenoids (170; 12.95%) are the predominant categories, consistent with the main text (Fig. 4A)

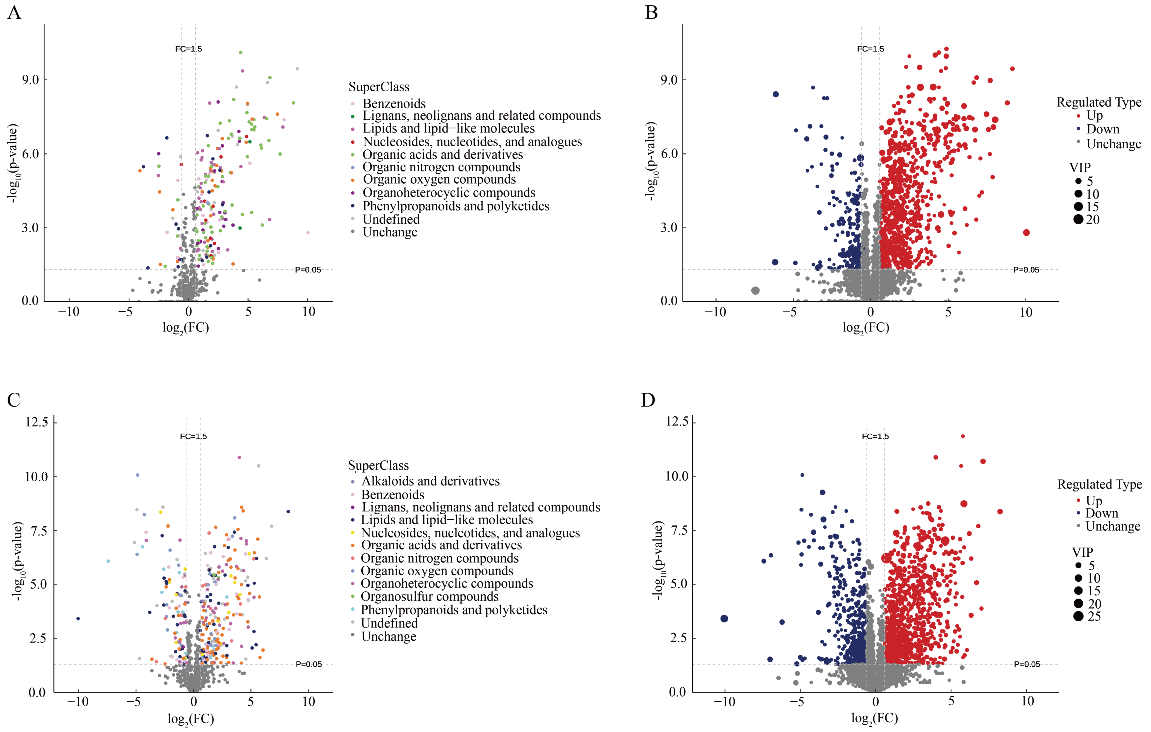


**Fig. S54. Volcano plots of differential metabolites: Fa-SG vs Fx-SG.** Combined volcano plots (4 panels): (A) Negative ionization, VIP-ranked; (B) Negative ionization, superclass-colored; (C) Positive ionization, superclass-colored; (D) Positive ionization, VIP-ranked. Thresholds: $|\log_2 FC| > 1$, $P < 0.05$. Metabolites above both thresholds are highlighted.

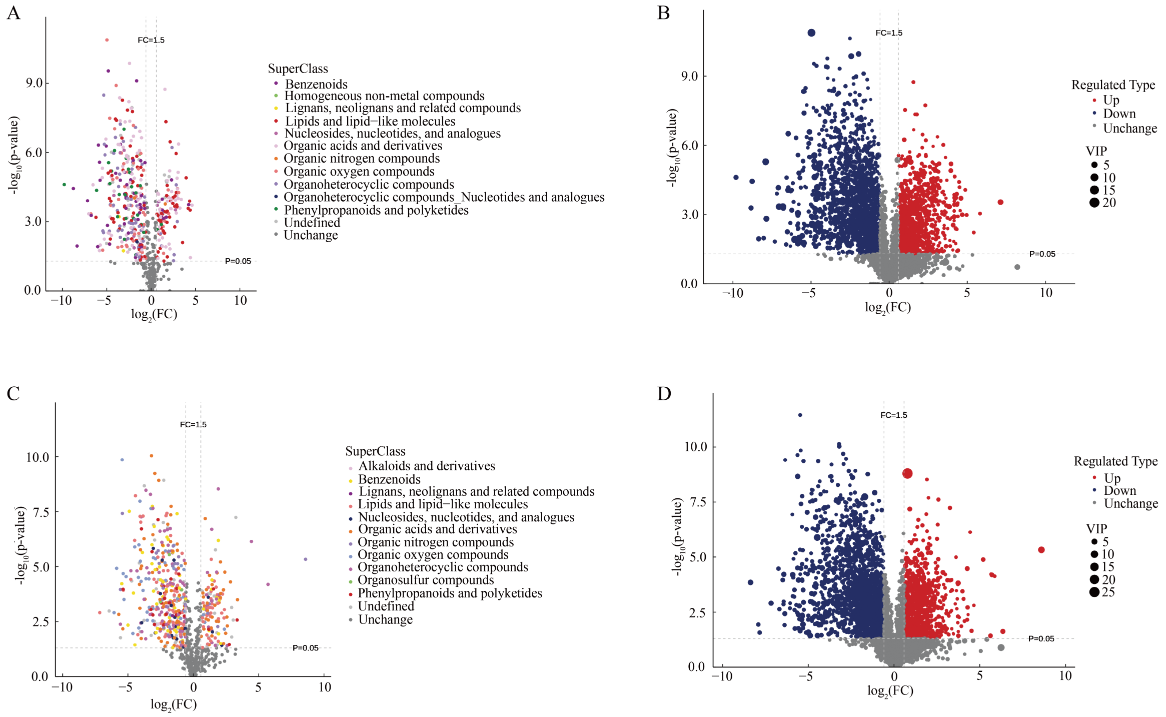


**Fig. S55. Volcano plots of differential metabolites: Fa-SG vs Se-SG.** Combined volcano plots in the same format as Fig. S54, comparing fundatrigenia salivary gland with sexuparae salivary gland. This comparison reveals metabolites specifically enriched in the gall-inducing morph relative to the non-galling morph.

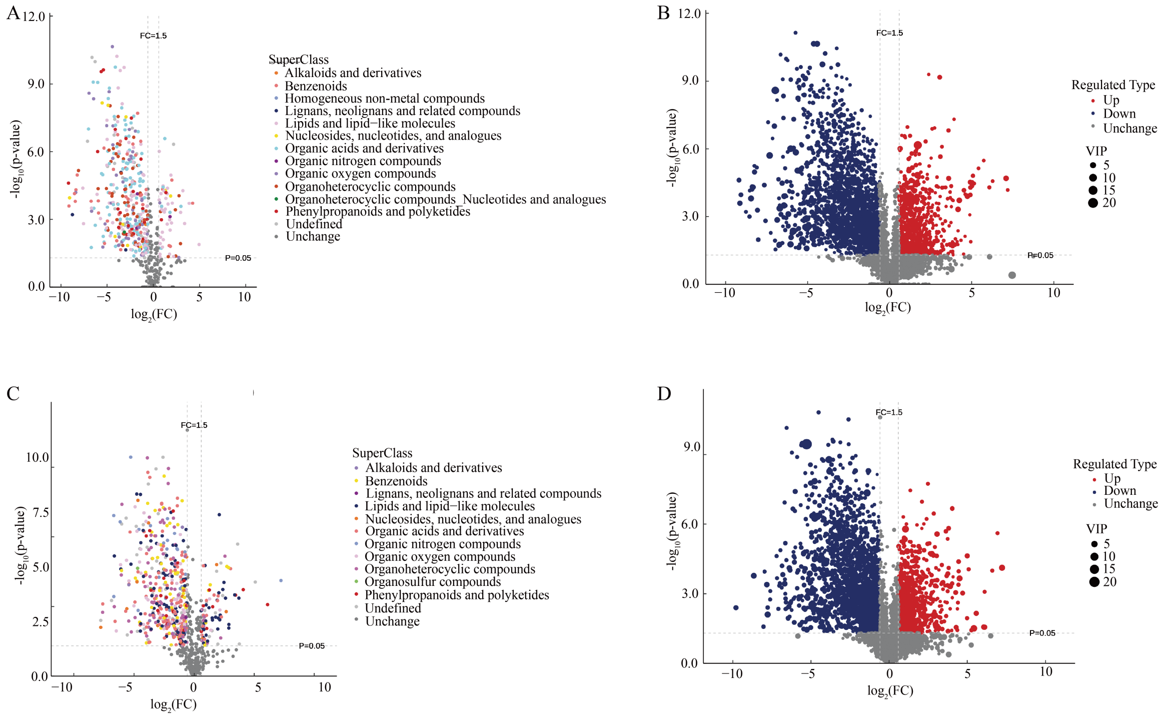


**Fig. S56. Volcano plots of differential metabolites: Fx-SG vs Se-SG.** Combined volcano plots comparing fundatrix salivary gland with sexuparae salivary gland, formatted as in Fig. S54.

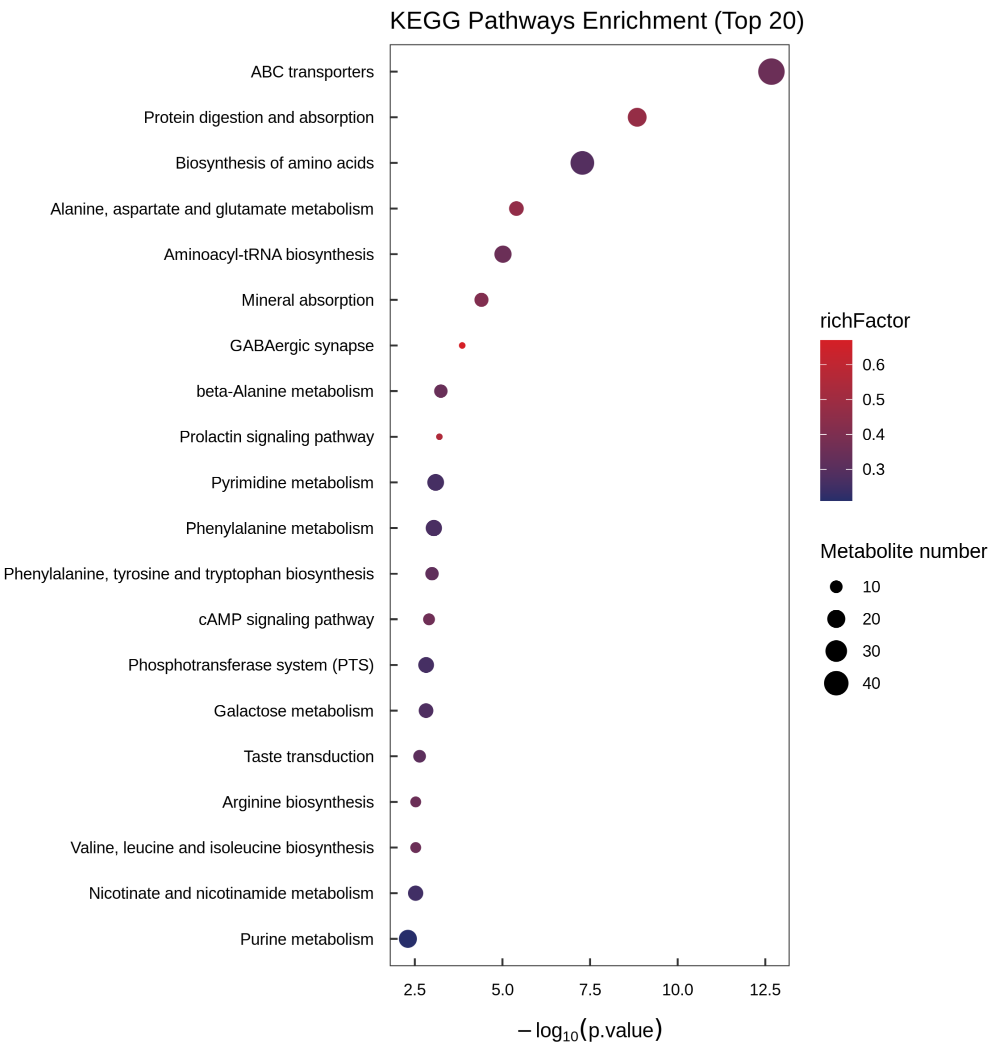


**Fig. S57. KEGG pathway enrichment of differential metabolites: one-way analysis.** KEGG metabolic pathway enrichment for metabolites differentially accumulated across Fa-SG, Fx-SG, and Se-SG by one-way ANOVA. Bubble size indicates gene count; color represents significance level. Amino acid metabolism and biosynthesis pathways are prominently enriched.

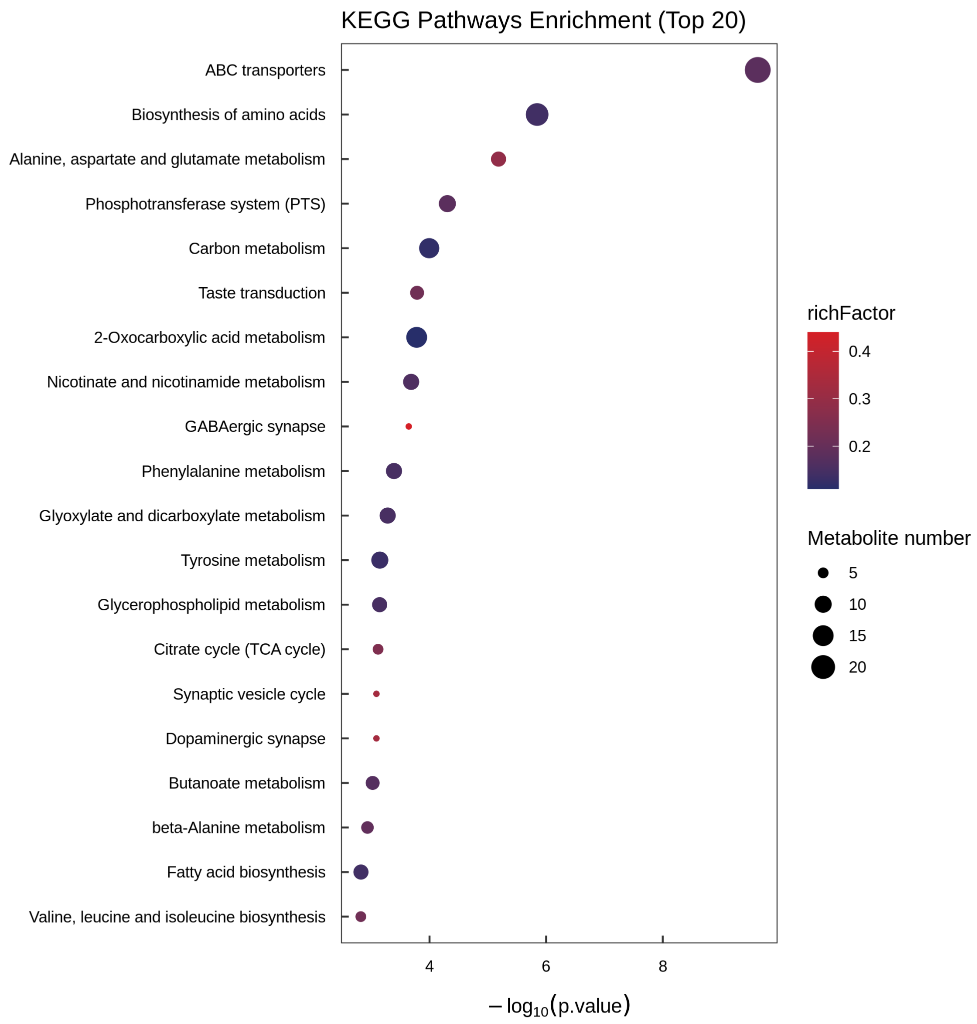


**Fig. S58. KEGG pathway enrichment of differential metabolites: Fa-SG vs Fa-S.** KEGG pathway enrichment bubble plot for metabolites differentially accumulated between fundatrigenia salivary gland and fundatrigenia saliva. This comparison reveals metabolites actively secreted from the gland into the saliva.

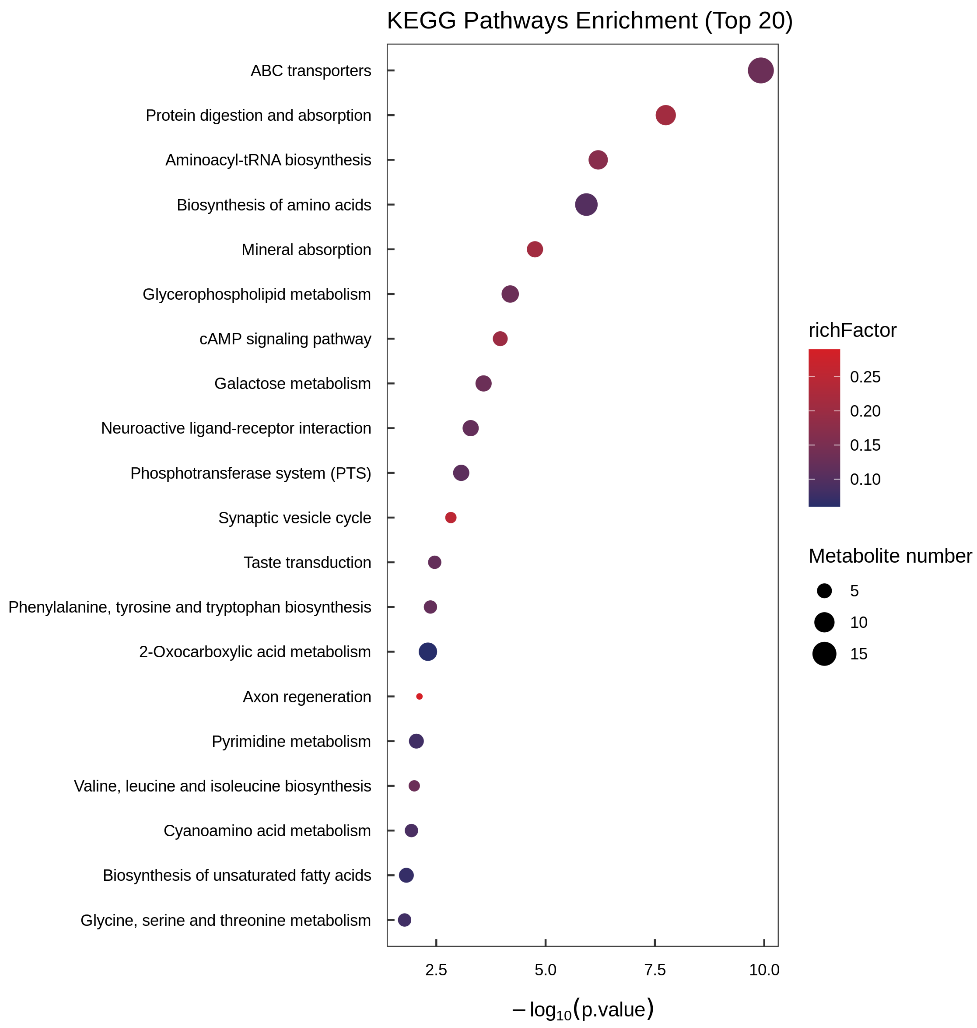


**Fig. S59. KEGG pathway enrichment of differential metabolites: Fa-SG vs Fx-SG.** KEGG pathway enrichment for metabolites differentially accumulated between fundatrigenia and fundatrix salivary glands, highlighting morph-specific metabolic specialization.

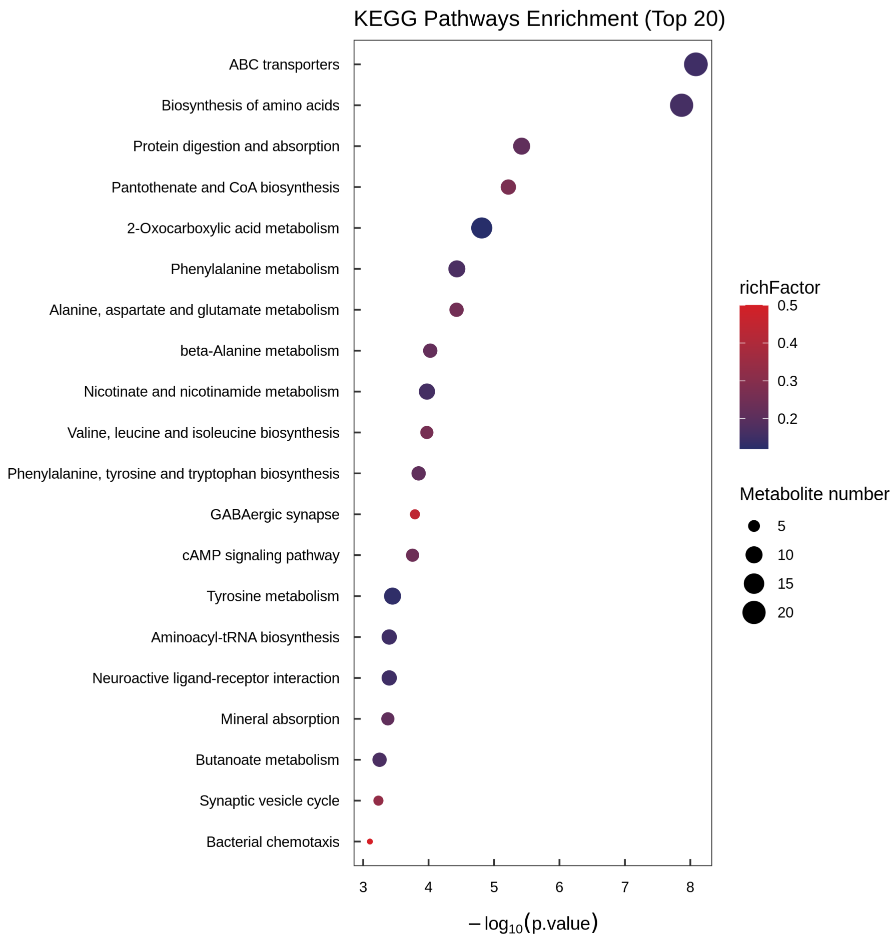


**Fig. S60. KEGG pathway enrichment of differential metabolites: Fa-SG vs Se-SG.** KEGG pathway enrichment comparing fundatrigenia salivary gland with sexuparae salivary gland metabolomes.

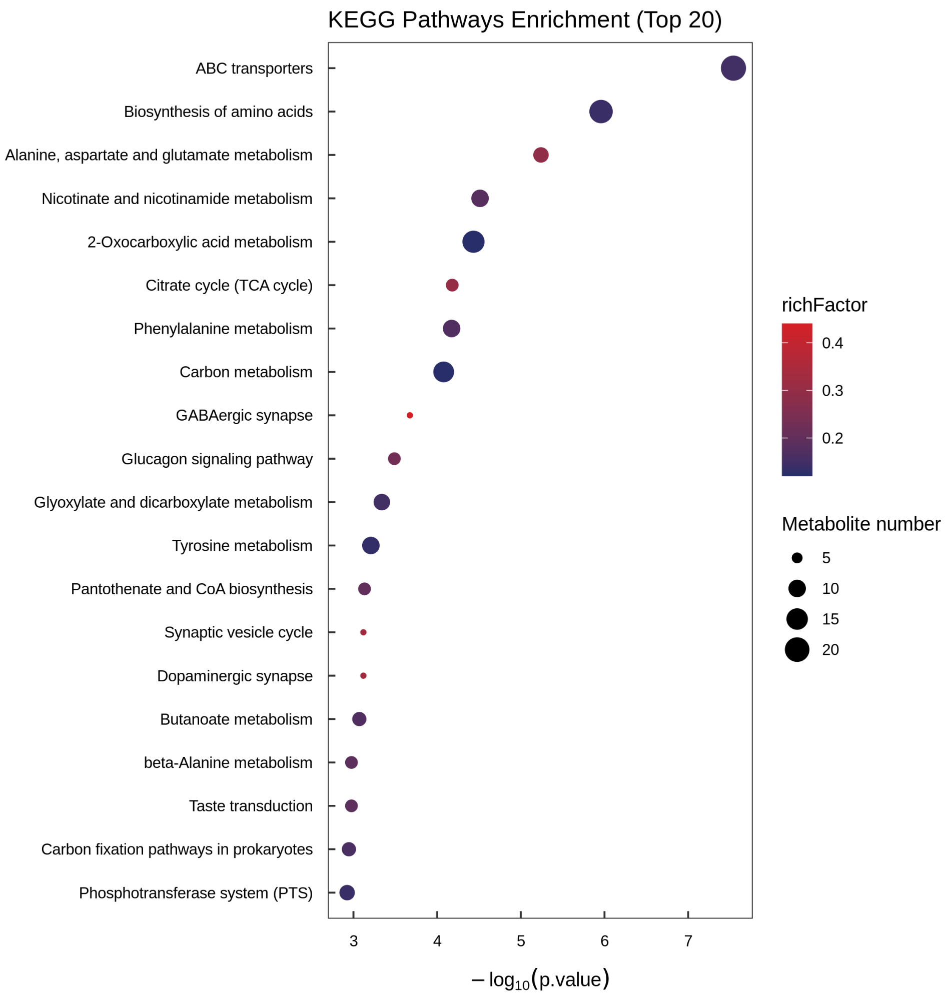


**Fig. S61. KEGG pathway enrichment of differential metabolites: Fx-S vs Fa-S.** KEGG pathway enrichment for metabolites differentially accumulated between fundatrix saliva and fundatrigenia saliva, corresponding to the volcano plot comparison in Fig. 4e.

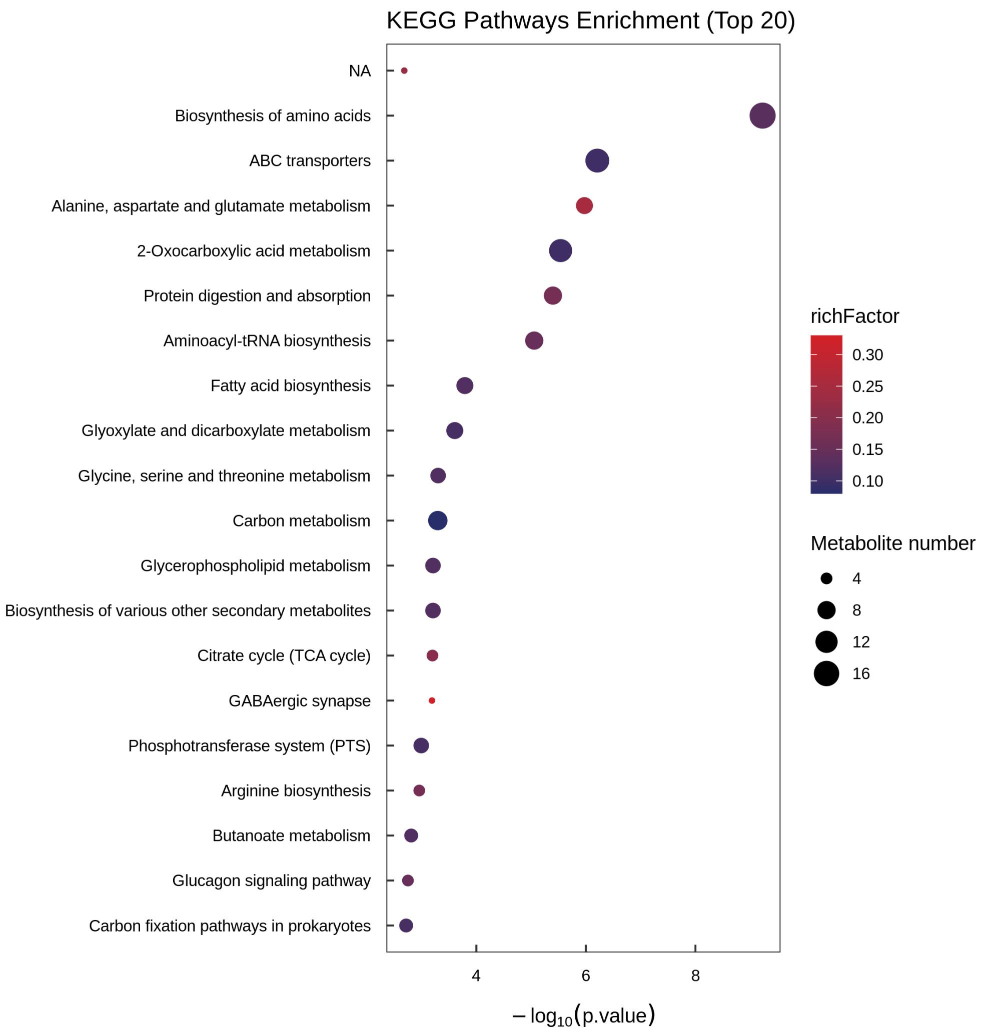


**Fig. S62. KEGG pathway enrichment of differential metabolites: Fx-SG vs Fx-S.** KEGG pathway enrichment comparing fundatrix salivary gland with fundatrix saliva, revealing metabolites retained in the gland versus those actively secreted into the saliva.

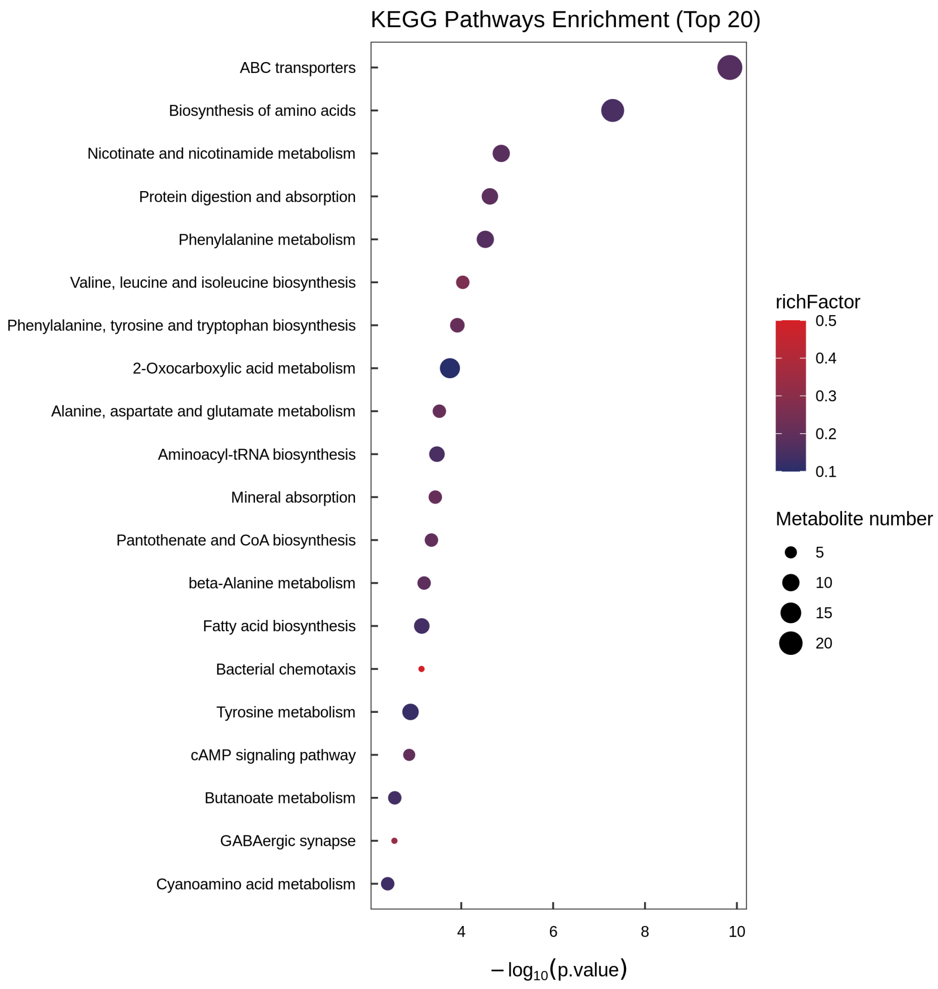


**Fig. S63. KEGG pathway enrichment of differential metabolites: Fx-SG vs Se-SG.** KEGG pathway enrichment comparing fundatrix salivary gland with sexuparae salivary gland metabolomes.

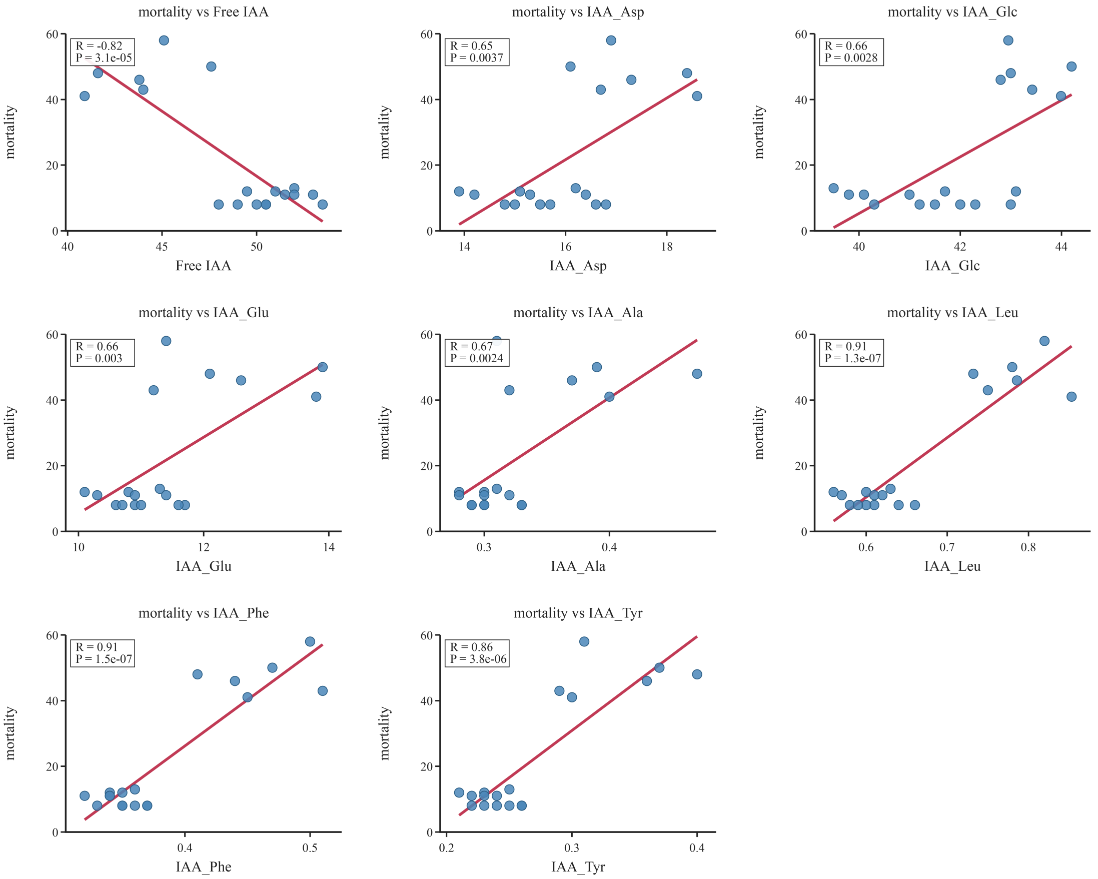


**Fig. S64. ACY1 RNAi phenotypic and auxin metabolite validation.** Comprehensive visualization of *ACY1* knockdown effects. Panels show the dose-dependent relationship between *ACY1* expression, galling phenotype, and auxin metabolite changes at the settlement zone. This figure supplements the main RNAi results presented in Fig. 5.

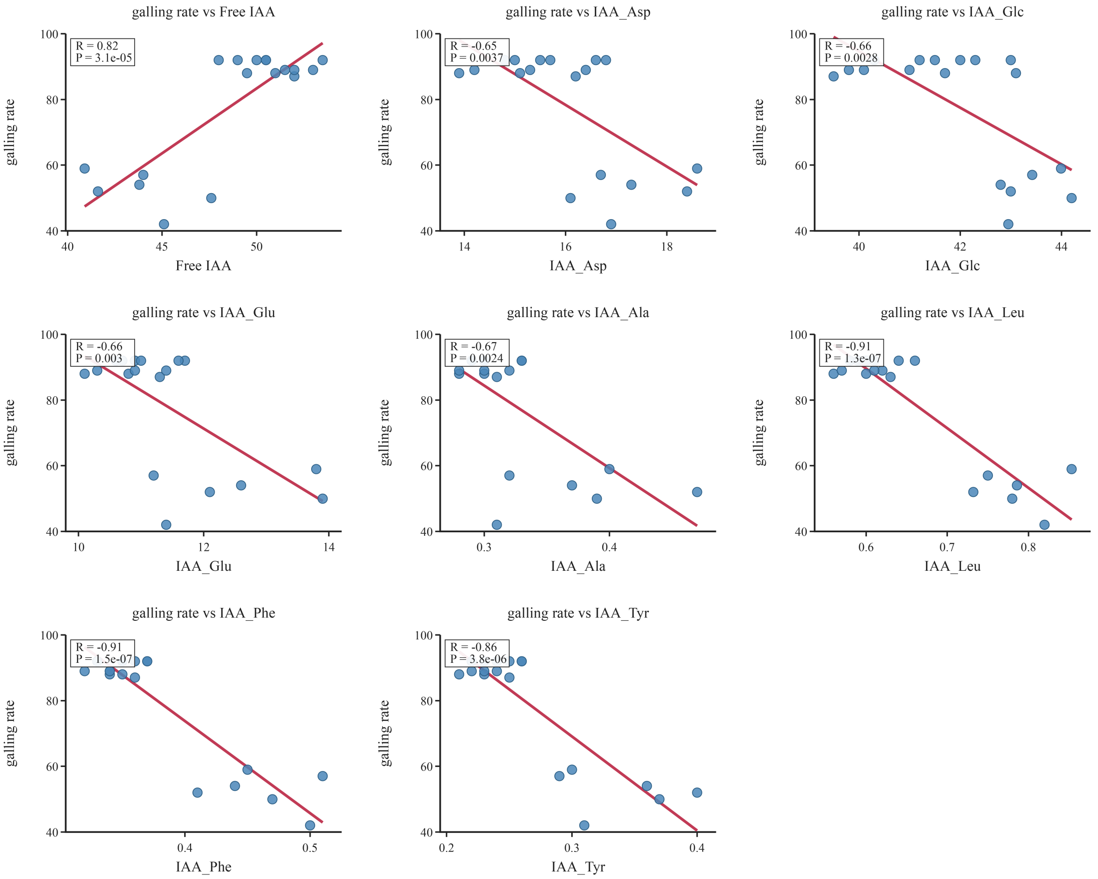


**Fig. S65. Correlations of galling rate with free auxin and individual conjugated auxins.** Galling rate was positively correlated with free auxin content and negatively correlated with the contents of individual conjugated auxins.

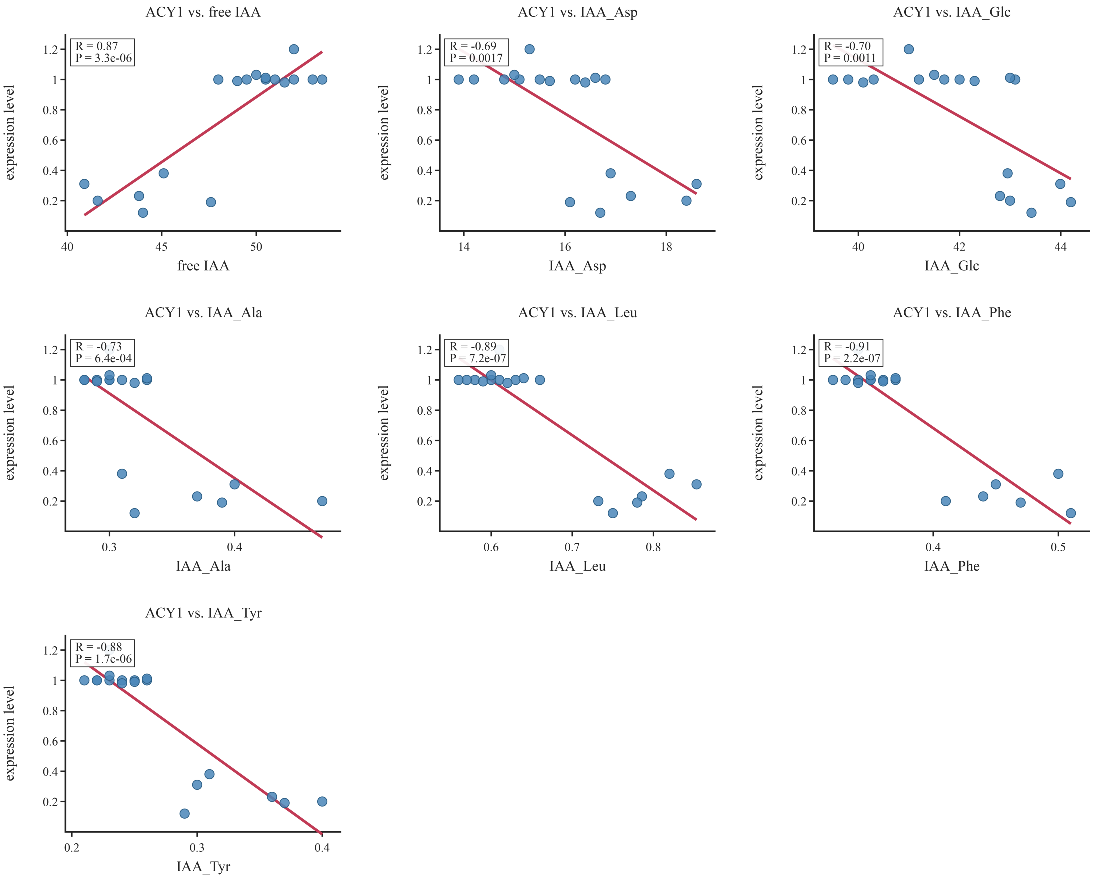


**Fig. S66. Correlation analysis of ACY1 enzyme activity and auxin metabolism**（4 μM）**.** Correlation matrix and regression plots showing the relationships among ACY1 concentration, free IAA levels and conjugated IAA species. Pearson correlation coefficients and significance levels are indicated.

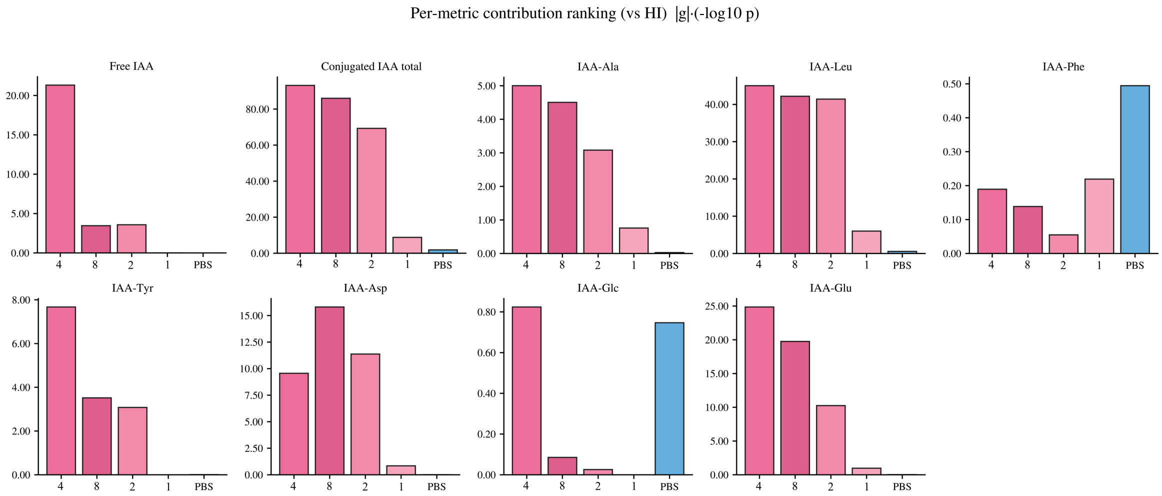


**Fig. S67. Proportion of variance in individual IAA species explained by each ACY1 treatment concentration ($R^2$).** Each axis represents one phenotypic or metabolic indicator; higher $R^2$ values indicate that a greater proportion of the within-group variation in that indicator is accounted for by the treatment. The 4 µM treatment exhibits the highest $R^2$ across most indicators, particularly for IAA-Leu, IAA-Phe, IAA-Tyr, and free IAA, indicating that this concentration exerts the most consistent and predictable effect on auxin redistribution and growth promotion.

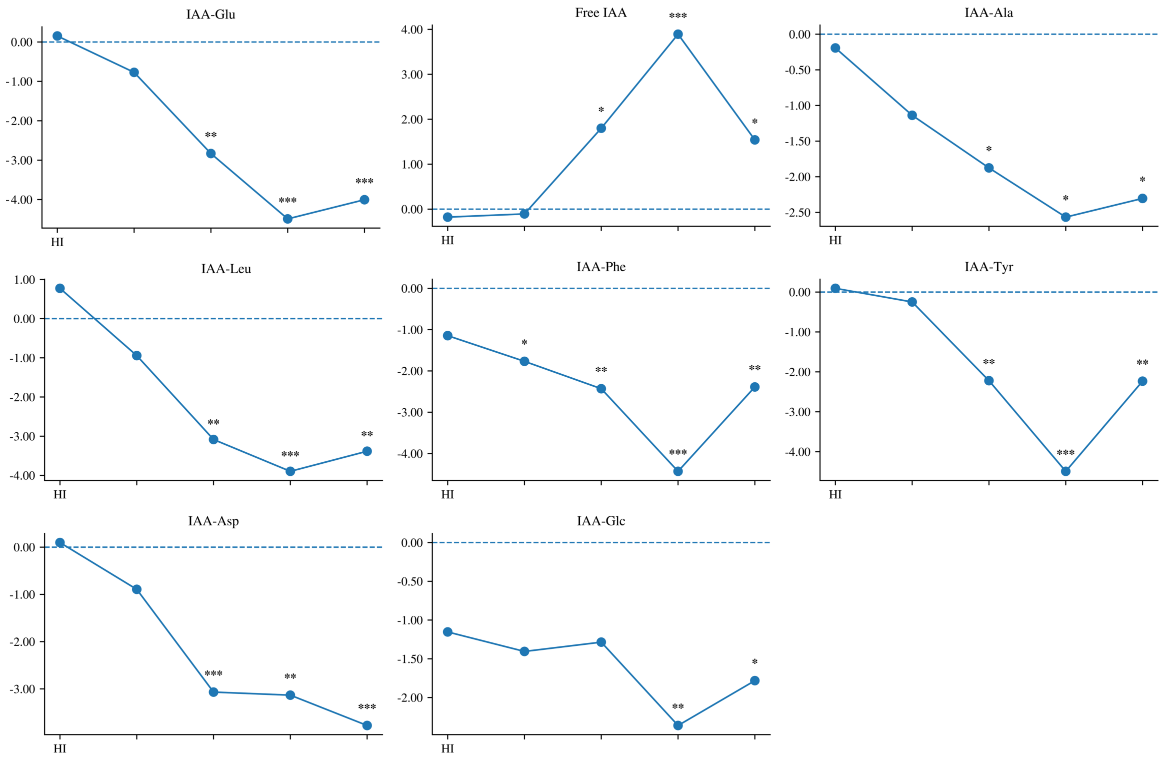


**Fig. S68. Composite impact score of each ACY1 concentration on IAA metabolites.** The 4 µM concentration yields the highest composite impact score, followed by 8 µM and 2 µM, while 1 µM shows the weakest overall effect.
