## Supplementary methods for "Chromosome gigantism and auxin deconjugation underpin gall induction in a horned gall aphid"

### 1. Sample collection

#### 1.1 Aphid collection

*Schlechtendalia chinensis* (Chinese horned gall aphid) samples were collected from naturally occurring galls on *Rhus chinensis* trees in Yunnan Province, China (E102°44′49″, N25°03′50″; elevation ~1,900 m a.s.l.) during July–October over two consecutive years. Three distinct morphs were collected: (i) fundatrices (Fx; gall initiators), harvested from early-stage galls (1–2 weeks post-induction) by carefully opening the gall wall under a stereomicroscope and extracting live aphids with fine forceps; (ii) fundatrigeniae (Fa; gall maintainers), collected from mid-stage galls (4–6 weeks post-induction) containing actively reproducing colonies; and (iii) sexuparae (Se; non-galling winged migrants), captured from mature galls prior to emergence using fine-mesh traps positioned over the gall ostiole. For genomic DNA extraction, adult fundatrigeniae were flash-frozen in liquid nitrogen within 30 s of collection to minimize DNA degradation. For proteomics and metabolomics, salivary glands and saliva were collected as described below (sections 5 and 6). All specimens were stored at −80°C until processing.

1.2 Host plant collection

Young expanding leaves of *R. chinensis* were collected from the same trees bearing aphid galls. Leaves were rinsed with sterile distilled water, blotted dry, flash-frozen in liquid nitrogen and stored at −80°C. For genome sequencing, leaf tissue from a single tree was used to minimize heterozygosity. For Hi-C library preparation, fresh young leaves were cross-linked with 1% formaldehyde immediately after harvest.

### 2. Genome sequencing and assembly

2.1 DNA extraction and library preparation

High-molecular-weight genomic DNA was extracted from flash-frozen *S. chinensis* adult fundatrigeniae (~500 individuals pooled) using a modified CTAB method with additional RNase A and Proteinase K treatment, followed by purification with AMPure PB magnetic beads (Pacific Biosciences). DNA integrity was assessed by pulsed-field gel electrophoresis (PFGE; target fragment size >50 kb) and quantified using a Qubit 3.0 fluorometer (Thermo Fisher Scientific). For *R. chinensis*, genomic DNA was extracted from young leaf tissue (~5 g) using the DNeasy Plant Maxi Kit (Qiagen) with modifications for polyphenol-rich tissue (addition of 2% PVP-40 and 0.1% β-mercaptoethanol to the lysis buffer). PacBio HiFi sequencing libraries were prepared using the SMRTbell Express Template Prep Kit 2.0 (Pacific Biosciences) with a target insert size of 15–20 kb and sequenced on a PacBio Sequel IIe platform in CCS mode, generating ~30× genome coverage for each species. Illumina paired-end libraries (insert size 350 bp) were prepared using the NEBNext Ultra II DNA Library Prep Kit (New England Biolabs) and sequenced on a NovaSeq 6000 platform (2 × 150 bp) to ~100× coverage for each species.

### 2.2 Hi-C library preparation

Hi-C libraries were prepared from cross-linked tissue (aphid: ~200 adult fundatrigeniae; host: ~2 g young leaf tissue) using the Arima-HiC Kit (Arima Genomics) following the manufacturer's protocol. Briefly, chromatin was fixed with 1% formaldehyde for 10 min at room temperature, quenched with glycine, and digested with the restriction enzyme cocktail. Proximity-ligated DNA was sheared to ~300–500 bp using a Covaris M220 focused ultrasonicator, size-selected with AMPure XP beads (Beckman Coulter), and sequenced on a NovaSeq 6000 platform (2 × 150 bp) to generate >100× genome coverage of Hi-C read pairs.

### 2.3 Genome assembly and scaffolding

PacBio HiFi reads were assembled into contigs using hifiasm v0.16[1] with default parameters. Contigs were polished in two rounds: first with PacBio HiFi reads using pbmm2 + DeepVariant, then with Illumina reads using Pilon v1.24[2]. Hi-C reads were aligned to polished contigs using BWA-MEM v0.7.17[3] and processed with the Juicer pipeline v1.6[4]. Chromosome-level scaffolding was performed using 3D-DNA v180922[5] and manually curated using Juicebox Assembly Tools v1.11.08[6].

2.4 Assembly quality assessment

Assembly completeness was evaluated using BUSCO v5.4[7] against the insecta_odb10 (for *S. chinensis*) and eudicots_odb10 (for *R. chinensis*) lineage datasets in genome and protein modes. The *S. chinensis* assembly showed 98.0% BUSCO completeness at the genome level and 94.4% at the annotation level; the *R. chinensis* assembly showed 97.4% and 93.8%, respectively (Table S7). Merqury v1.3[8] was used to estimate consensus quality (QV) and k-mer completeness using Illumina reads. Genome size was estimated independently using GenomeScope 2.0[9] with Illumina k-mer frequency distributions (k = 21).

2.5 Karyotype validation

Chromosome numbers were validated by two independent cytogenetic approaches. (i) DAPI-FISH: metaphase chromosome spreads were prepared from embryonic tissue of parthenogenetic aphids and root-tip meristems of *R. chinensis* seedlings following standard colchicine-arrested protocols. Chromosomes were stained with 4′,6-diamidino-2-phenylindole (DAPI; 1 μg $ml^{-1}$) and imaged on a Zeiss Axio Imager Z2 fluorescence microscope equipped with a 100× oil-immersion objective and a CoolSNAP HQ2 CCD camera. Karyograms were arranged using the ImageJ plugin KaryotypeJ. (ii) Giemsa staining: metaphase spreads were G-banded following standard trypsin-Giemsa protocols and scored independently by two investigators.

**3. Genome annotation**

#### 3.1 Repeat annotation

Repetitive sequences were identified using a combined de novo and homology-based approach. De novo repeat libraries were constructed for each species using RepeatModeler v2.0.3[10] (including the LTR structural search pipeline). The species-specific libraries were merged with RepBase v20181026[11] and used as input for RepeatMasker v4.1.2[12] to annotate and soft-mask repeats genome-wide. Tandem repeats were independently identified using Tandem Repeats Finder v4.09[13].

#### 3.2 Gene prediction

Protein-coding genes were predicted on repeat-masked genomes using an evidence-driven pipeline integrating three lines of evidence. (i) Ab initio prediction: AUGUSTUS v3.4.0[14], GlimmerHMM v3.0.4[15] and Genscan[16] were trained on species-specific training sets derived from full-length transcript alignments. (ii) Transcript-based evidence: RNA-seq reads from multiple tissues (salivary glands, whole body, gall tissue and leaves) were assembled using Trinity v2.13.2[17] and aligned to the genome using PASA v2.5.2[18]. (iii) Homology-based evidence: protein sequences from six related aphid species (*Acyrthosiphon pisum*, *Myzus persicae*, *Rhopalosiphum maidis*, *Viteus vitifoliae*, *Diuraphis noxia*, *Sipha flava*) and, for the plant, five Sapindales genomes (*Citrus sinensis*, *Mangifera indica*, *Pistacia vera*, *Acer yangbiense, Dimocarpus longan*) were aligned to the target genome using GeMoMa v1.8[19]. All evidence was integrated using EvidenceModeler v1.1.1[20] to generate consensus gene models. Genes shorter than 50 amino acids or with premature stop codons were filtered.

#### 3.3 Functional annotation

Predicted proteins were functionally annotated by homology searches against six databases: (i) NR (NCBI non-redundant protein database, BLASTP, e-value < $1 \times 10^{-5}$); (ii) Swiss-Prot (BLASTP, e-value < $1 \times 10^{-5}$); (iii) KOG/eggNOG v5.0[21] (eggNOG-mapper v2.1.9); (iv) KEGG (KofamScan v1.3.0[22] against the KEGG Orthology database); (v) InterPro (InterProScan v5.59-91.0[23], searching Pfam, SMART, CDD, PANTHER, ProSiteProfiles and Gene3D); (vi) Gene Ontology (GO) terms were assigned from InterPro2GO mappings and BLASTP hits to Swiss-Prot. Signal peptides were predicted using SignalP v6.0[24] and transmembrane domains using TMHMM v2.0[25].

**4. Comparative genomics**

4.1 Orthology inference and gene family evolution

Orthogroups were inferred among *S. chinensis*, *R. chinensis* and eight additional insect and plant species using OrthoFinder v2.5.4[26] with default parameters (DIAMOND v2.0.15 for all-vs-all BLASTP; MCL clustering with inflation parameter 1.5). The species tree was inferred from a concatenated alignment of single-copy orthologues using RAxML v8.2.12[27] under the PROTGAMMALG model. Divergence times were estimated using MCMCTree in PAML v4.9[28] with calibration points from the TimeTree database[29]. Gene family expansion and contraction were modelled using CAFE v5[30] with a birth–death model of gene gain and loss, testing significance at $P < 0.05$ (conditional probability per family). Expanded families were defined as those with a significantly higher gene count in the focal lineage; contracted families as those with a significantly lower count. Enrichment of expanded and contracted families was assessed separately by hypergeometric tests against GO and KEGG backgrounds, with Benjamini–Hochberg (BH) FDR correction at $Q < 0.05$.

#### 4.2 Positive selection analysis

Single-copy orthologues shared among *S. chinensis* (or *R. chinensis*) and at least four outgroup species were extracted, aligned at the codon level using PRANK v.170427[31] guided by the species tree, and filtered for alignment quality using Gblocks v0.91b[32] (minimum block length 5 codons; no gap columns allowed). Positive selection was tested using the branch-site model (model = 2, NSsites = 2) in codeml of PAML v4.9[28], with the focal species specified as foreground. The null model fixed $\omega_2 = 1$ was compared with the alternative model ($\omega_2 > 1$) using a likelihood ratio test (LRT; $\chi^2$, df = 1). *P* values were adjusted for multiple testing using BH-FDR. Genes with FDR < 0.05 were designated as positively selected genes (PSGs). KEGG and GO enrichment of PSG sets was performed as for expanded families (section 4.1).

#### 4.3 Synteny and segmental duplication analysis

Intra- and inter-species syntenic blocks were identified using MCScanX[33] with parameters: minimum block size = 5 genes, maximum gap = 25 genes. For inter-species comparison, protein sequences of *S. chinensis* were compared against *Viteus vitifoliae* and *Acyrthosiphon pisum* using BLASTP (e-value < 1 × 10−10), and collinear blocks were extracted. Macrosynteny was visualized using the Python package JCVI v1.1.12[34]. Intraspecies segmental duplications (SDs) were called based on all-vs-all BLASTP within each genome (identity ≥90%, alignment length ≥100 aa, e-value < $1 \times 10^{-10}$), filtered for non-tandem pairs separated by ≥10 intervening genes, and summarized by chromosome. Circos v0.69-9[35] was used for genome-wide visualization of gene density, TE density, GC content, SD positions and inter-chromosomal links.

#### 4.4 Chromosome 1 functional analysis

To characterize the functional specialization of *S. chinensis* chr1, we performed the following analyses: (i) Gene-category density comparison: seven functional gene categories (zinc finger, UGT, opsin, crystallin/sHSP20, PIF1 helicase, SETMAR and TE-associated) were annotated across all chromosomes using InterPro domain assignments. Gene counts per category were normalized by chromosome gene content, and $\chi^2$ tests were used to compare observed versus expected frequencies on chr1. (ii) KEGG enrichment of chr1-resident genes and SD-overlapping genes was performed using hypergeometric tests (BH-FDR $Q < 0.05$). (iii) Re-annotation of the lysine degradation pathway: all 60 genes assigned to ko00310 in expanded families were manually inspected by BLASTP against Swiss-Prot (e-value $< 1 \times 10^{-10}$) and InterPro domain scanning. Each gene was classified into one of five categories: SETMAR fusion protein, canonical Mariner transposase, FLJ37770/GVQW3-like degenerate Mariner, other histone methyltransferase (NSD2, Mes-4) or uncharacterized. (iv) Gene cluster detection: tandem gene arrays on chr1 were defined as ≥3 genes of the same family within a 500 kb window. Opsin (12.75–13.00 Mb), crystallin/sHSP20 (75.78–75.83 Mb and 80.92–80.94 Mb) and PIF1 helicase loci were mapped and visualized using custom R scripts and ggplot2.

### 5. Salivary proteomics

#### 5.1 Salivary gland dissection and saliva collection

Salivary glands were dissected from three morphs under a stereomicroscope (Leica M205 FA) in ice-cold PBS (137 mM NaCl, 2.7 mM KCl, 10 mM $Na_2HPO_4$, 1.8 mM $KH_2PO_4$, pH 7.4) supplemented with 1× protease inhibitor cocktail (Roche cOmplete ULTRA). For each replicate, glands from 500 individuals were pooled. Dissected glands were flash-frozen and stored at −80°C. Five sample groups were generated: Fx-SG (fundatrix salivary gland), Fa-SG (fundatrigenia salivary gland), Se-SG (sexuparae salivary gland), Fx-S (fundatrix saliva) and Fa-S (fundatrigenia saliva). For saliva collection, live fundatrices and fundatrigeniae were placed on an artificial feeding membrane (Parafilm M stretched over a sucrose–amino acid diet) for 12 h. The lower surface of the membrane, onto which saliva was deposited, was rinsed with PBS + 0.1% Tween-20 and concentrated using 3-kDa molecular-weight-cutoff filters (Millipore Amicon Ultra). Three biological replicates were prepared per sample group.

#### 5.2 Protein extraction and digestion

Samples were lysed in SDT buffer (4% SDS, 100 mM Tris-HCl pH 7.6, 1 mM DTT) by ultrasonication (3 × 30 s, 80 W) on ice. Lysates were heated at 95°C for 5 min, cooled, and centrifuged at 16,000 g for 15 min. Protein concentrations were determined using the BCA Protein Assay Kit (Pierce). Proteins were digested using the filter-aided sample preparation (FASP) method[36]: 200 μg protein per sample was loaded onto 30-kDa Microcon filters, washed with UA buffer (8 M urea in 0.1 M Tris-HCl pH 8.5), alkylated with 50 mM iodoacetamide in UA for 30 min in the dark, washed three times with 50 mM $NH_4HCO_3$, and digested with sequencing-grade trypsin (Promega; 1:50 enzyme:protein ratio) at 37°C for 16 h. Peptides were eluted, vacuum-dried and resuspended in 0.1% formic acid.

#### 5.3 LC–MS/MS analysis

Peptides were separated on a reversed-phase $C^{18}$ column (75 µm × 250 mm, 1.8 µm; Thermo Fisher Scientific) using an EASY-nLC 1200 system (Thermo Fisher Scientific) with a 120-min linear gradient from 5% to 35% acetonitrile in 0.1% formic acid at a flow rate of 300 nL $min^{-1}$. Eluted peptides were analysed on an Orbitrap Exploris 480 mass spectrometer (Thermo Fisher Scientific) operated in data-dependent acquisition (DDA) mode: full MS scans were acquired at 60,000 resolution (m/z 350–1,500); the top 20 precursor ions were selected for HCD fragmentation (normalized collision energy 30%) and MS/MS analysis at 15,000 resolution. Dynamic exclusion was set to 30 s.

5.4 Database search and quantification

Raw files were searched against the predicted *S. chinensis* proteome (19,109 entries) appended with common contaminants using MaxQuant v2.1.3.0[37] with the following settings: trypsin/P specificity with up to two missed cleavages; carbamidomethylation of cysteine as fixed modification; oxidation of methionine and N-terminal acetylation as variable modifications; precursor mass tolerance 20 ppm (first search) and 4.5 ppm (main search); fragment mass tolerance 20 ppm; peptide and protein FDR < 1%. Label-free quantification (LFQ) was enabled using the MaxLFQ algorithm[38] with a minimum ratio count of 2 and the "match between runs" option. Proteins quantified in ≥2 of 3 replicates in at least one group were retained for downstream analysis.

5.5 Differential abundance and functional analysis

LFQ intensities were $log_2$-transformed. Proteins with valid values in at least 2 of 3 replicates in at least one group were retained, and the remaining missing values were imputed from a down-shifted normal distribution (width 0.3 SD, downshift 1.8 SD) on a per-sample basis to approximate low-abundance signals. Differential abundance between sample groups was assessed using the limma package, whose empirical Bayes moderation stabilizes per-protein variance estimates and is therefore well suited to experiments with a small number of replicates. For each protein, a linear model was fitted to the $\log_2$ LFQ values with lmFit; pairwise group contrasts were specified with makeContrasts, applied with contrasts.fit, and moderated *t*-statistics and *P* values were obtained with eBayes. *P* values were adjusted across all proteins by the Benjamini–Hochberg method. Proteins with $|\log_2 FC| > 1$ and adjusted $P < 0.05$ were designated as significantly differentially abundant. Subcellular localization was predicted using DeepLoc v2.0[39]. Enzyme-class enrichment was tested by hypergeometric tests comparing enzyme-class frequencies in saliva-enriched proteins against the full proteome background. KEGG and GO enrichment used the same framework as in section 4.1. All analyses were performed in R v4.3.1.

### 6. Untargeted metabolomics

#### 6.1 Sample preparation

Metabolites were extracted from the same five sample groups as proteomics (section 5.1). Each sample (~10 mg for salivary glands; ~50 µL for saliva) was mixed with 400 µL ice-cold 80% methanol, vortexed for 30 s, sonicated on ice for 10 min, incubated at −20°C for 1 h, and centrifuged at 14,000g for 20 min at 4°C. The supernatant was collected, vacuum-dried, and reconstituted in 100 µL 50% methanol for analysis. Quality control (QC) samples were prepared by pooling equal aliquots from all samples.

#### 6.2 UPLC–MS/MS analysis

Metabolomic profiling was performed on a Vanquish UHPLC system coupled to a Q Exactive HFX Orbitrap mass spectrometer (Thermo Fisher Scientific). Chromatographic separation used a Hypersil Gold C18 column (100 × 2.1 mm, 1.9 µm; Thermo Fisher Scientific) at 40°C with mobile phases A (0.1% formic acid in water) and B (0.1% formic acid in methanol). The gradient was: 0–2 min 2% B; 2–12 min 2–98% B; 12–14 min 98% B; 14–14.1 min 98–2% B; 14.1–17 min 2% B. The flow rate was 0.2 mL min−1. The MS operated in both positive and negative electrospray ionization (ESI) modes at 120,000 resolution for full scan (m/z 70–1,050) with data-dependent MS/MS acquisition at 30,000 resolution (top 5, NCE 20/40/60). QC samples were injected every 10 samples.

6.3 Data processing and annotation

Raw data were processed using Compound Discoverer v3.3 (Thermo Fisher Scientific) with the following workflow: peak alignment, peak picking (minimum peak intensity 1 × 105), gap filling, adduct grouping and compound assignment. Metabolite identification was performed at four confidence levels following the Metabolomics Standards Initiative (MSI) guidelines: Level 1 (matched to authentic standards; mzCloud score ≥80), Level 2 (putative annotation; matched to spectral databases including mzCloud, HMDB and MassBank), Level 3 (putative class; matched by molecular formula and class prediction), and Level 4 (unknown). A total of 1,475 metabolites were annotated at Levels 1–3. Chemical classification was assigned using ClassyFire[40]. Metabolite peak areas were normalized to QC sample medians. Differential metabolites were defined using $|\log_2 FC| > 1$ and $P < 0.05$ (Welch's *t*-test, BH-FDR corrected). KEGG pathway enrichment used hypergeometric tests with BH correction ($Q < 0.05$).

**7. RNA interference and auxin profiling**

#### 7.1 dsRNA synthesis

A 400-bp fragment of S. chinensis *ACY1* (nucleotides 451–850 of the coding sequence), spanning the predicted M20 metallopeptidase catalytic domain, was selected as the RNAi target. This region was amplified by RT–PCR from fundatrigenia salivary gland cDNA using gene-specific primers each appended with a T7 promoter sequence at the 5′ end (ds*ACY1*-F: 5′-TAATACGACTCACTATAGATGTCATTTGTTCCTGATGAA-3′; ds*ACY1*-R: 5′-TAATACGACTCACTATAGTATCAAAACCAACTGTTAACT-3′). The purified PCR product was used as template for simultaneous bidirectional in vitro transcription and double-stranded RNA (dsRNA) synthesis using the MEGAscript RNAi Kit (Thermo Fisher Scientific) following the manufacturer's protocol. After transcription, the reaction was treated with DNase and RNase to remove the DNA template and single-stranded RNA. The resulting dsRNA was purified using the kit-supplied nuclease-free water elution procedure, quantified with a NanoDrop 2000 spectrophotometer (Thermo Fisher Scientific) and verified by 1% agarose gel electrophoresis. A dsRNA targeting green fluorescent protein (GFP) of similar length served as the negative control for off-target effects.

#### 7.2 dsRNA delivery by artificial feeding and RT–qPCR

Fundatrix first-instar nymphs (within 24 h of birth) were collected and placed on artificial feeding chambers. Each chamber comprised a glass cylinder sealed at one end with a double Parafilm membrane enclosing ~500 μL of feeding solution (15% sucrose, 0.5× Murashige–Skoog salts, pH 7.0) supplemented with dsRNA at 500 ng/μL. Three treatment groups were established: (i) blank control (diet only, no dsRNA), (ii) GFP (diet + GFP dsRNA negative control), and (iii) *ACY1* dsRNA (diet + *ACY1* dsRNA at 500 ng μL−1). Nymphs fed for 24 h were transferred to young *R. chinensis* leaves maintained on wet cotton in sealed Petri dishes (25°C, 16:8 L:D photoperiod). Six biological replicates per group (200 nymphs per replicate) were established. Total RNA was extracted from 100 aphids per replicate using TRIzol reagent (Invitrogen). cDNA was synthesized using HiScript III RT SuperMix (Vazyme). RT–qPCR was performed on a QuantStudio 5 Real-Time PCR System (Applied Biosystems) using ChamQ Universal SYBR qPCR Master Mix (Vazyme). *ACY1* expression was normalized to the reference gene *β-actin* using the $2^{-\Delta\Delta Ct}$ method. Primer sequences are listed in Table S1.

7.3 Phenotype scoring

Galling rate was scored 25 d after transfer as the proportion of aphids associated with visible gall initiation (epidermal swelling or pocket formation). Mortality was scored as the proportion of aphids that died without initiating galls. The sum of galling rate and mortality accounted for all individuals in each replicate.

7.4 Targeted auxin profiling at the settlement zone

Host tissue (~50 mg) surrounding each settlement site was excised, weighed, and flash-frozen. Auxin metabolites were extracted in 80% methanol containing 1 ng $mL^{-1}$ [$^{13}C_6$]-IAA as internal standard, and analysed by targeted LC–MS/MS on a QTRAP 6500+ system (SCIEX) operated in multiple reaction monitoring (MRM) mode. The following analytes were quantified: free IAA, IAA-Ala, IAA-Leu, IAA-Phe, IAA-Tyr, IAA-Asp, IAA-Glu and IAA-Glc. Calibration curves were prepared using authentic standards (Sigma-Aldrich and OlChemIm). Concentrations were normalized to tissue fresh weight (pmol $g^{-1}$ FW).

#### 7.5 Statistical analysis of RNAi data

All metrics (galling rate, mortality, *ACY1* expression, free IAA and conjugated IAA species) were analysed using six biological replicates per group. Multi-group comparisons used one-way ANOVA followed by Tukey's HSD post hoc test (adjusted $P < 0.05$ considered significant). Data are presented as mean ± s.d. Pearson correlation coefficients ($r$) were computed between *ACY1* expression and each IAA metabolite, galling rate and mortality across all 18 samples (3 groups × 6 replicates). $P$ values for correlations were BH-FDR corrected across the full correlation matrix (8 IAA metrics × 3 traits). The coefficient of determination ($R^2 = r^2$) was used to express the proportion of variance in each IAA metric explained by *ACY1* expression. Correlation heatmaps and radar plots were generated in R v4.3.1 using the packages corrplot and fmsb.

### 8. Cell-free expression and aphid-feeding-mimic delivery

#### 8.1 Cell-free protein expression

The full-length *ACY1* coding sequence was amplified from *S. chinensis* fundatrigenia salivary gland cDNA and cloned into the pEU-E01-MCS-N-His expression vector (CellFree Sciences), which carries the SP6 promoter and the E01 translational enhancer optimized for wheat germ cell-free protein synthesis. The recombinant plasmid was verified by Sanger sequencing. *In vitro* transcription was performed using the SP6 RNA polymerase supplied in the WEPRO7240H Premium Expression Kit (CellFree Sciences) according to the manufacturer's protocol. Translation reactions were carried out in the bilayer-diffusion format at 26°C for 18 h, in which the WEPRO7240H wheat germ extract layer was overlaid with SUB-AMIX SGC feeding buffer to enable continuous supply of substrates and removal of inhibitory byproducts. After translation, the reaction mixtures were collected as crude cell-free lysates without further affinity purification. Expression of full-length His-tagged ACY1 (predicted ~45.8 kDa) was verified by SDS–PAGE (12% gel, Coomassie staining) and confirmed by western blotting with a mouse anti-His tag monoclonal antibody (1:2,000; Cell Signaling Technology) followed by HRP-conjugated goat anti-mouse IgG (1:5,000; Abcam) and ECL detection (Bio-Rad). The total protein concentration of each crude lysate was determined by the BCA Protein Assay Kit (Pierce, Thermo Fisher Scientific) using bovine serum albumin as the standard. Because the crude lysate contains both recombinant ACY1 and the endogenous wheat germ proteins required for cell-free translation, the BCA assay measures the total protein content of the preparation rather than the absolute amount of ACY1 alone. For comparability across treatments, the total-protein concentration of each preparation was converted to an apparent molar value by dividing the BCA-measured mass concentration (mg $mL^{-1}$) by the predicted molecular mass of ACY1. All concentrations reported throughout this study and in Fig. 6 therefore refer to this BCA-based apparent ACY1 concentration of the crude lysate, not to the concentration of purified ACY1. Aliquots of each crude lysate were flash-frozen and stored at −80°C until use. Heat-inactivated controls (HI-ACY1) were prepared from the same crude lysate by incubation at 95°C for 15 min to abolish enzymatic activity while retaining the lysate matrix. Empty-vector control lysate (EV) was prepared by running the identical wheat germ cell-free reaction with the empty pEU-E01-MCS-N-His vector lacking the *ACY1* insert.

8.2 Aphid-feeding-mimic delivery to *R. chinensis* leaves

Young expanding leaves of *R. chinensis* were collected from healthy trees, rinsed with sterile distilled water and blotted dry. Crude lysates expressing recombinant ACY1 were diluted in sterile PBS to four apparent ACY1 concentrations of 1, 2, 4 and 8 μM, defined according to the BCA-based total-protein measurement described in section 8.1 (i.e. total lysate protein mass divided by the predicted molecular mass of ACY1). All control lysates (EV and HI-ACY1) were diluted with PBS to the same total-protein concentration as the corresponding active treatment so that the lysate matrix was held constant across all comparisons. Each leaf was first punctured with a sterile fine needle (gauge 30; BD PrecisionGlide) to create a series of small wounds across the adaxial surface, simulating the penetration of aphid stylets into leaf tissue. Punctured leaves were then submerged in the corresponding enzyme solution and subjected to vacuum infiltration (–0.08 MPa, 2 min, repeated twice with brief release between cycles) to deliver the enzyme preparation into the leaf intercellular apoplastic space. After infiltration, excess solution was gently blotted from the leaf surface. Infiltrated leaves were placed in humidity chambers (sealed Petri dishes lined with moist filter paper) and incubated at 25°C under a 16:8 L:D photoperiod for 12 h before sampling. Six biological replicates per treatment group were used (n = 6). Two control groups were included, each subjected to the same needle puncture and vacuum infiltration: (i) EV (empty-vector lysate control) and (ii) HI-ACY1 (heat-inactivated lysate control), both matched in total-protein concentration to the highest active treatment.

#### 8.3 Auxin metabolite measurement

After 12 h of incubation, treated leaves (~50 mg per replicate) were excised, flash-frozen and processed for targeted auxin profiling as described in section 7.4. Free IAA and seven conjugated species were quantified. The free-to-conjugated IAA ratio was calculated as [Free IAA] / Σ[Conjugated IAA]. Free IAA proportion was calculated as [Free IAA] / [Total IAA] × 100%.

8.4 Statistical analysis of cell-free expression data

All comparisons used six biological replicates per treatment group. Each active-ACY1 apparent concentration (1, 2, 4 and 8 μM, as defined in section 8.1) was compared against two control groups (EV and HI-ACY1) using two-sided Welch's *t*-tests. For each indicator (free IAA, each conjugated IAA, free-to-conjugated ratio, free IAA proportion), multiple comparisons of 4 concentrations against the same control were corrected using the Holm method (family-wise error rate < 0.05). The comparison between EV and HI-ACY1 was reported separately with uncorrected *P* values. Standardized effect sizes were calculated as Hedges' *g* to enable cross-indicator comparison of effect magnitudes. To identify the concentration producing the strongest overall effect, a composite impact score was computed for each concentration $d$: for each indicator $j$, the Z-score $z_{d,j} = (\bar{a}_{d,j} - \bar{a}_{control,j}) / SD_{control,j}$ was calculated, and the composite score $S_d = \Sigma|z_{d,j}|$ was summed across indicators. A higher $S_d$ indicates a stronger overall departure from controls. Data are presented as mean ± s.d. Significance thresholds: * = $P < 0.05$, ** = $P < 0.01$, *** = $P < 0.001$. Analyses were performed in R v4.3.1.

**9. Histological analysis**

To assess the effect of *ACY1* knockdown on host cell morphology, leaf tissue from the settlement zone was collected from control and *ACY1*-RNAi groups at 14 days post-transfer. Tissues were fixed in FAA (formalin–acetic acid–ethanol; 10% formalin, 5%

acetic acid, 50% ethanol) for 24 h, dehydrated through a graded ethanol series (70–100%), embedded in paraffin wax using a Leica ASP300 tissue processor, and sectioned at 8 μm using a Leica RM2255 rotary microtome. Sections were stained with Safranin O–Fast Green for general histology. Cell diameters were measured along the long axis using ImageJ v1.53 software in three tissue layers: epidermis, palisade mesophyll and spongy mesophyll ($n = 30$ cells per tissue type per group). Statistical comparisons between groups used two-sided Mann–Whitney *U* tests.

**10. Scanning electron microscopy**

Mouthpart morphology of fundatrix, fundatrigenia and sexuparae was examined by scanning electron microscopy (SEM). Specimens were fixed in 2.5% glutaraldehyde in 0.1 M phosphate buffer (pH 7.2) at 4°C overnight, post-fixed in 1% osmium tetroxide for 1 h, dehydrated through a graded ethanol series, and critical-point-dried using a Leica EM CPD300. Dried specimens were sputter-coated with gold–palladium (~10 nm) using a Leica EM ACE200 and imaged on a Zeiss Sigma 300 field-emission SEM at 3–5 kV.

**Supplementary Methods References**

[1] Cheng H, Concepcion GT, Feng X, Zhang H, Li H. Haplotype-resolved de novo assembly using phased assembly graphs with hifiasm. Nat Methods 2021;18:170–5. https://doi.org/10.1038/s41592-020-01056-5.

[2] Poplin R, Chang P-C, Alexander D, Schwartz S, Colthurst T, Ku A, et al. A universal SNP and small-indel variant caller using deep neural networks. Nat Biotechnol 2018;36:983–7. https://doi.org/10.1038/nbt.4235.

[3] Vasimuddin Md, Misra S, Li H, Aluru S. Efficient architecture-aware acceleration of BWA-MEM for multicore systems, 2019, p. 314–24. https://doi.org/10.1109/IPDPS.2019.00041.

[4] Durand NC, Shamim MS, Machol I, Rao SSP, Huntley MH, Lander ES, et al. Juicer provides a one-click system for analyzing loop-resolution Hi-C experiments. Cell Syst 2016;3:95–8. https://doi.org/10.1016/j.cels.2016.07.002.

[5] Dudchenko O, Batra SS, Omer AD, Nyquist SK, Hoeger M, Durand NC, et al. De novo assembly of the Aedes aegypti genome using Hi-C yields chromosome-length scaffolds. Science 2017;356:92–5. https://doi.org/10.1126/science.aal3327.

[6] Durand NC, Robinson JT, Shamim MS, Machol I, Mesirov JP, Lander ES, et al. Juicebox provides a visualization system for Hi-C contact maps with unlimited zoom. Cell Syst 2016;3:99–101. https://doi.org/10.1016/j.cels.2015.07.012.

[7] Manni M, Berkeley MR, Seppey M, Simão FA, Zdobnov EM. BUSCO update: novel and streamlined workflows along with broader and deeper phylogenetic coverage for scoring of eukaryotic, prokaryotic, and viral genomes. Mol Biol Evol 2021;38:4647–54. https://doi.org/10.1093/molbev/msab199.

[8] Rhie A, Walenz BP, Koren S, Phillippy AM. Merqury: reference-free quality, completeness, and phasing assessment for genome assemblies. Genome Biol 2020;21:245. https://doi.org/10.1186/s13059-020-02134-9.

[9] Ranallo-Benavidez TR, Jaron KS, Schatz MC. GenomeScope 2.0 and Smudgeplot for reference-free profiling of polyploid genomes. Nat Commun 2020;11:1432. https://doi.org/10.1038/s41467-020-14998-3.

[10] Flynn JM, Hubley R, Goubert C, Rosen J, Clark AG, Feschotte C, et al. RepeatModeler2 for automated genomic discovery of transposable element families. Proc Natl Acad Sci U S A 2020;117:9451–7. https://doi.org/10.1073/pnas.1921046117.

[11] Storer J, Hubley R, Rosen J, Wheeler TJ, Smit AF. The Dfam community resource of transposable element families, sequence models, and genome annotations. Mob DNA 2021;12:2. https://doi.org/10.1186/s13100-020-00230-y.

[12] Smit AFA, Hubley R, Green P. RepeatMasker Open-4.0 2013. http://www.repeatmasker.org.

[13] Benson G. Tandem repeats finder: a program to analyze DNA sequences. Nucleic Acids Res 1999;27:573–80. https://doi.org/10.1093/nar/27.2.573.

[14] Brůna T, Hoff KJ, Lomsadze A, Stanke M, Borodovsky M. BRAKER2: automatic eukaryotic genome annotation with GeneMark-EP+ and AUGUSTUS supported by a protein database. NAR Genomics Bioinforma 2021;3:lqaa108. https://doi.org/10.1093/nargab/lqaa108.

[15] Majoros WH, Pertea M, Salzberg SL. TigrScan and GlimmerHMM: two open source ab initio eukaryotic gene-finders. Bioinformatics 2004;20:2878–9. https://doi.org/10.1093/bioinformatics/bth315.

[16] Burge C, Karlin S. Prediction of complete gene structures in human genomic DNA. J Mol Biol 1997;268:78–94. https://doi.org/10.1006/jmbi.1997.0951.

[17] Grabherr MG, Haas BJ, Yassour M, Levin JZ, Thompson DA, Amit I, et al. Full-length transcriptome assembly from RNA-Seq data without a reference genome. Nat Biotechnol 2011;29:644–52. https://doi.org/10.1038/nbt.1883.

[18] Haas BJ, Delcher AL, Mount SM, Wortman JR, Smith RKJ, Hannick LI, et al. Improving the Arabidopsis genome annotation using maximal transcript alignment assemblies. Nucleic Acids Res 2003;31:5654–66. https://doi.org/10.1093/nar/gkg770.

[19] Keilwagen J, Hartung F, Grau J. GeMoMa: Homology-Based Gene Prediction Utilizing Intron Position Conservation and RNA-seq Data. Methods Mol Biol 2019;1962:161–77. https://doi.org/10.1007/978-1-4939-9173-0_9.

[20] Haas BJ, Salzberg SL, Zhu W, Pertea M, Allen JE, Orvis J, et al. Automated eukaryotic gene structure annotation using EVidenceModeler and the Program to Assemble Spliced Alignments. Genome Biol 2008;9:R7. https://doi.org/10.1186/gb-2008-9-1-r7.

[21] Hernández-Plaza A, Szklarczyk D, Botas J, Cantalapiedra CP, Giner-Lamia J, Mende DR, et al. eggNOG 6.0: enabling comparative genomics across 12 535 organisms. Nucleic Acids Res 2023;51:D389–94. https://doi.org/10.1093/nar/gkac1022.

[22] Aramaki T, Blanc-Mathieu R, Endo H, Ohkubo K, Kanehisa M, Goto S, et al. KofamKOALA: KEGG Ortholog assignment based on profile HMM and adaptive score threshold. Bioinformatics 2020;36:2251–2. https://doi.org/10.1093/bioinformatics/btz859.

[23] Paysan-Lafosse T, Blum M, Chuguransky S, Grego T, Pinto BL, Salazar GA, et al. InterPro in 2022. Nucleic Acids Res 2023;51:D418–27. https://doi.org/10.1093/nar/gkac993.

[24] Teufel F, Almagro Armenteros JJ, Johansen AR, Gíslason MH, Pihl SI, Tsirigos KD, et al. SignalP 6.0 predicts all five types of signal peptides using protein language models. Nat Biotechnol 2022;40:1023–5. https://doi.org/10.1038/s41587-021-01156-3.

[25] Hallgren J, Tsirigos KD, Pedersen MD, Almagro Armenteros JJ, Marcatili P, Nielsen H, et al. DeepTMHMM predicts alpha and beta transmembrane proteins using deep neural networks. bioRxiv 2022. https://doi.org/10.1101/2022.04.08.487609.

[26] Emms DM, Kelly S. OrthoFinder: phylogenetic orthology inference for comparative genomics. Genome Biol 2019;20:238. https://doi.org/10.1186/s13059-019-1832-y.

[27] Minh BQ, Schmidt HA, Chernomor O, Schrempf D, Woodhams MD, von Haeseler A, et al. IQ-TREE 2: new models and efficient methods for phylogenetic inference in the genomic era. Mol Biol Evol 2020;37:1530–4. https://doi.org/10.1093/molbev/msaa015.

[28] Yang Z. PAML 4: phylogenetic analysis by maximum likelihood. Mol Biol Evol 2007;24:1586–91. https://doi.org/10.1093/molbev/msm088.

[29] Kumar S, Suleski M, Craig JM, Kasprowicz AE, Sanderford M, Li M, et al. TimeTree 5: An Expanded Resource for Species Divergence Times. Mol Biol Evol 2022;39:msac174. https://doi.org/10.1093/molbev/msac174.

[30] Mendes FK, Vanderpool D, Fulton B, Hahn MW. CAFE 5 models variation in evolutionary rates among gene families. Bioinformatics 2020;36:5516–8. https://doi.org/10.1093/bioinformatics/btaa1022.

[31] Ranwez V, Douzery EJP, Cambon C, Chantret N, Delsuc F. MACSE v2: toolkit for the alignment of coding sequences accounting for frameshifts and stop codons. Mol Biol Evol 2018;35:2582–4. https://doi.org/10.1093/molbev/msy159.

[32] Steenwyk JL, Buida TJI, Li Y, Shen X-X, Rokas A. ClipKIT: a multiple sequence alignment trimming software for accurate phylogenomic inference. PLoS Biol 2020;18:e3001007. https://doi.org/10.1371/journal.pbio.3001007.

[33] Wang Y, Tang H, DeBarry JD, Tan X, Li J, Wang X, et al. MCScanX: a toolkit for detection and evolutionary analysis of gene synteny and collinearity. Nucleic Acids Res 2012;40:e49. https://doi.org/10.1093/nar/gkr1293.

[34] Tang H, Krishnakumar V, Zeng X, Xu Z, Taranto A, Lomas JS, et al. JCVI: a versatile toolkit for comparative genomics analysis. iMeta 2024;3:e211. https://doi.org/10.1002/imt2.211.

[35] Krzywinski M, Schein J, Birol I, Connors J, Gascoyne R, Horsman D, et al. Circos: an information aesthetic for comparative genomics. Genome Res 2009;19:1639–45. https://doi.org/10.1101/gr.092759.109.

[36] Wiśniewski JR, Zougman A, Nagaraj N, Mann M. Universal sample preparation method for proteome analysis. Nat Methods 2009;6:359–62. https://doi.org/10.1038/nmeth.1322.

[37] Tyanova S, Temu T, Cox J. The MaxQuant computational platform for mass spectrometry-based shotgun proteomics. Nat Protoc 2016;11:2301–19. https://doi.org/10.1038/nprot.2016.136.

[38] Cox J, Hein MY, Luber CA, Paron I, Nagaraj N, Mann M. Accurate proteome-wide label-free quantification by delayed normalization and maximal peptide ratio extraction, termed MaxLFQ. Mol Cell Proteomics 2014;13:2513–26. https://doi.org/10.1074/mcp.M113.031591.

[39] Thumuluri V, Almagro Armenteros JJ, Johansen AR, Nielsen H, Winther O. DeepLoc 2.0: multi-label subcellular localization prediction using protein language models. Nucleic Acids Res 2022;50:W228–34. https://doi.org/10.1093/nar/gkac278.

[40] Djoumbou Feunang Y, Eisner R, Knox C, Chepelev L, Hastings J, Owen G, et al. ClassyFire: automated chemical classification with a comprehensive, computable taxonomy. J Cheminformatics 2016;8:61. https://doi.org/10.1186/s13321-016-0174-y.
